# High resolution profiling of MHC-II peptide presentation capacity, by Mammalian Epitope Display, reveals SARS-CoV-2 targets for CD4 T cells and mechanisms of immune-escape

**DOI:** 10.1101/2021.03.02.433522

**Authors:** Franz Josef Obermair, Florian Renoux, Sebastian Heer, Chloe Lee, Nastassja Cereghetti, Giulia Maestri, Yannick Haldner, Robin Wuigk, Ohad Iosefson, Pooja Patel, Katherine Triebel, Manfred Kopf, Joanna Swain, Jan Kisielow

**Affiliations:** Repertoire Immune Medicine, Cambridge, USA; Tepthera subsidiary, Schlieren, Switzerland; Institute of Molecular Health Sciences, ETH Zürich, Switzerland

## Abstract

Understanding the mechanisms of immune evasion is critical for formulating an effective response to global threats like SARS-CoV2. We have fully decoded the immune synapses for multiple TCRs from acute patients, including cognate peptides and the presenting HLA alleles. Furthermore, using a newly developed mammalian epitope display platform (MEDi), we determined that several mutations present in multiple viral isolates currently expanding across the globe, resulted in reduced presentation by multiple HLA class II alleles, while some increased presentation, suggesting immune evasion based on shifting MHC-II peptide presentation landscapes. In support, we found that one of the mutations present in B1.1.7 viral strain could cause escape from CD4 T cell recognition in this way. Given the importance of understanding such mechanisms more broadly, we used MEDi to generate a comprehensive analysis of the presentability of all SARS-CoV-2 peptides in the context of multiple common HLA class II molecules. Unlike other strategies, our approach is sensitive and scalable, providing an unbiased and affordable high-resolution map of peptide presentation capacity for any MHC-II allele. Such information is essential to provide insight into T cell immunity across distinct HLA haplotypes across geographic and ethnic populations. This knowledge is critical for the development of effective T cell therapeutics not just against COVID-19, but any disease.

## Introduction

Severe acute respiratory syndrome coronavirus 2 (SARS-CoV2) is the infectious agent responsible for the severe acute respiratory syndrome^1,2^, which caused the worldwide COVID-19 pandemic with over two million fatalities. Several companies are now providing vaccines inducing humoral and cellular responses against SARS-CoV2, but for long lasting protection, generation of T cell memory will be required^3^, even if pre-existing T cell immunity to common cold coronavirus might play a role^4,5,6^. Because protection by antibodies is related to protein function (e.g. blocking receptors that are required for viral cell entry), and/or protein localization (surface expression to allow opsonizing antibodies to bind), it has limited potential target space increasing selection pressure for pathogen escape. Protection by T cells, on the other hand, relies entirely on TCR recognition of pathogen-derived peptides presented by MHC and is mostly independent of physiological function or localization of the target protein. Consequently, while only particular epitopes of surface proteins allow targeting by neutralizing antibodies, many peptides can serve as T cell targets, providing a much bigger epitope space for therapeutic development. Clearly, a high-resolution map of all SARS-CoV-2 presentable peptides resolved on different HLA alleles would greatly help these efforts.

The main approaches used currently, analysis of MHC-eluted peptides by liquid chromatography with tandem mass spectrometry (LC-MS/MS) and in-silico prediction algorithms, have contributed to the understanding of peptide presentation. However, they do not provide complete presentability landscapes across many HLAs. LC-MS/MS analysis allows the identification of thousands of naturally presented peptides, but it is technically challenging and requires very large numbers of cells (i.e. 10^8^ to 10^10^) for good coverage^7,8^. Moreover, presentation of peptides for which T cell reactivity was confirmed by ELISPOT, can be missed^8^. The limited sensitivity of LC-MS/MS is especially problematic when working with small tissue samples like human biopsies. To solve this problem, dendritic cells can be pulsed with a pathogen or protein of interest and MHC-associated peptide proteomics (MAPPs) can be performed^9^. This method is particularly useful for HLA class II, but due to the expression of multiple HLA alleles on DCs, determination of the individual restriction requires additional experiments. Attempting to circumvent these problems, computational prediction methods have been developed and are relatively reliable in identifying strong (IC50 < 50 nM) MHC I-binders^10^. While for MHC II the algorithms are also improving^11^, the efficiency in predicting MHC-binding peptides is quite variable and limited. In this respect, the recently improved NetMHCIIpan4 shows better performance than conventional binding prediction algorithms^12^, but is accurate only for a limited number of alleles, owing to the lack of suitable peptide datasets for training. To circumvent this, a recently published study improved algorithm performance using yeast-display peptide libraries^13^. Still, there is a big gap from the several HLAs with high-quality in-silico prediction scores and the thousands of unique HLA alleles present in the human population.

Predicting antigen presentation by MHC is further complicated by the fact that it is a dynamic process and can change depending on the physiological state of the cell. It is also regulated by tightly controlled chaperones like HLA-DM^14^, dysregulation of which has been linked to autoimmune disease progression^15,16^, while high expression of HLA-DM correlated with improved survival in cancer patients^17^. Thus, an unbiased method, testing pure peptide presentation capacity of the MHC not obscured by other physiological factors, would help getting the complete picture of all possible pMHC ligands present in a given protein. This reductionist approach would provide a basic set of allele-specific peptides (the presentable peptide space) ready for the generation of peptide libraries for screening of T cell reactivities or the generation of pMHC tetramers. Taking this set as a basis, subsets of presented peptides could be derived by incorporating protein processing and chaperone functions, dependent on cellular state and chaperone expression levels.

In this work, we studied T cell recognition of the SARS-CoV-2 virus by de-orphaning TCRs from acute COVID-19 patients. We also tested the potential of mutations from SARS-CoV-2 B1.1.7 strain to influence peptide presentation by utilizing a novel mammalian epitope display system called MEDi. This platform allows unbiased, affordable testing of the presentability of all possible peptides derived from a protein in the context of any MHC class II allele. We describe validation experiments and use MEDi to provide a comprehensive presentability map of all SARS-CoV-2 peptides, WT and mutated, in the context of common HLA class II molecules. We found that several mutations resulted in reduced peptide presentation by multiple HLA alleles, a few increased it, and one caused escape from CD4 T cell recognition by altering peptide presentation. While further experiments are needed to fully appreciate the biological consequences of these observations, our results suggest that immune evasion based on shifting peptide presentation away from well recognized CD4 epitopes could be one of them.

Given the importance of CD4 T cells in controlling B cell and CD8 T cell responses in COVID-19 patients, the results described here may help guide the generation of vaccines or therapeutics designed to elicit efficient cellular immunity.

## Results

### De-orphaning TCRs from the BAL of acute COVID-19 patients

Although T cell SARS-CoV-2 reactivities against peptides scattered across the viral genome have been reported, analyses that comprehensively decode “immune synapses”, including TCR alpha and beta chain sequences, the recognized peptide and the presenting HLA, are sparse. To overcome these limitations, we used the MCR2 technology^18^ (Fig.1A) and single chain trimers^19^ linked to the intracellular domain of the TCR zeta chain (SCTz)^20,21^, to de-orphan TCRs of enriched clonotypes from the bronchoalveolar lavages of COVID-19 patients, described recently by Liao et al^22^. Liao et al provided high resolution single cell data indicating aberrant cellular responses and identified expanded T cell clonotypes, but they neither decoded their antigenic specificity, nor the HLA restriction. To address this, we cloned 109 most enriched TCRs (supplementary data excel file S1), expressed them in a T cell line and performed an unbiased epitope screening. This included MCR2 libraries containing all possible 23aa SARS-CoV-2-derived peptides (1aa shifts through all proteins) and libraries containing all possible 10aa SARS-CoV2-derived peptides presented in the context of SCTz. This setup allowed for an unprecedented, complete screen of all SARS-CoV-2 peptides in the context of all HLAs from every patient (Table 1). Screening these patient specific MCR2 libraries of approximately 120.000 different peptide-MCR2 combinations and 60.000 peptide-SCTz combinations required at least 4 rounds of enrichment (Fig1B and not shown) before single cell clones revealed the specific peptides and the presenting HLA alleles (Fig 1C and not shown). As expected not all TCRs showed reactivity against SARS-CoV-2 antigens, but we identified the cognate peptides and the HLA restriction for 8 CD4, and 3 CD8 TCRs (Fig.1CD).

**Table 1.**
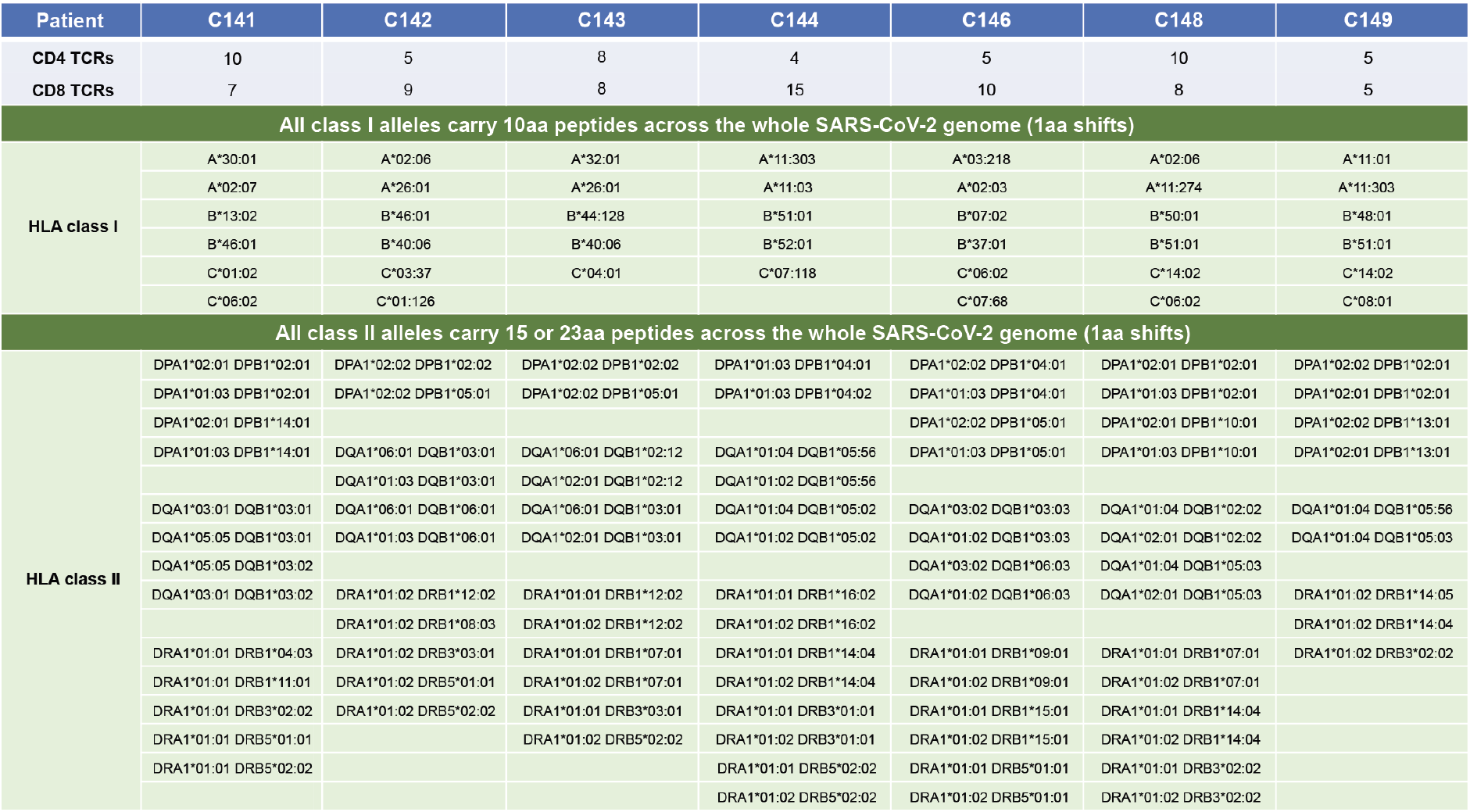
Patient HLAs.

**Figure 1.**
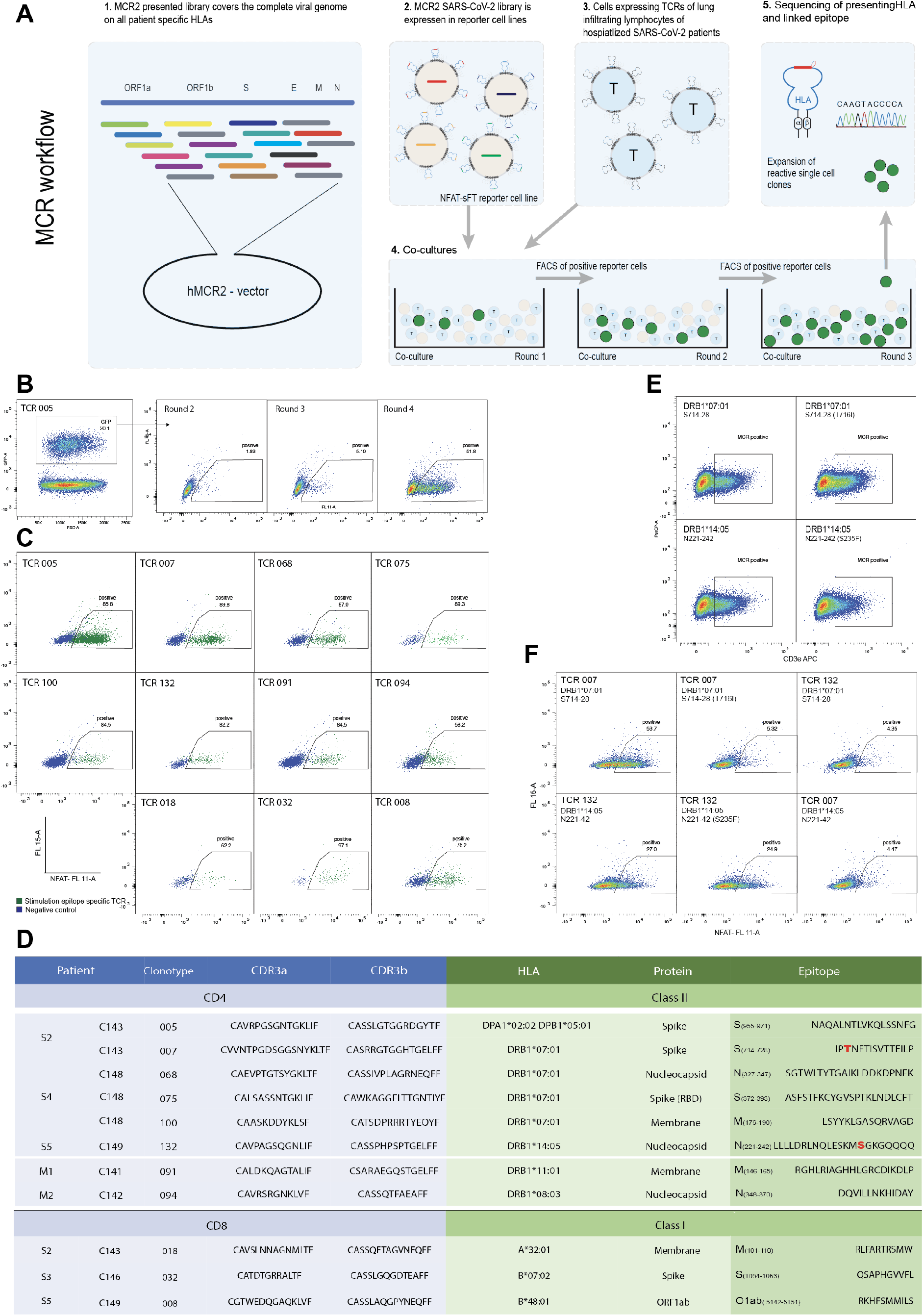
De-orphaning TCRs from the BAL of COVID-19 patients by MCR2 screening. **A.** Schematic representation of the TCR de-orphaning process. **B.** MCR2-SARS-CoV-2^+^ or SCTz-SARS-CoV-2^+^ 16.2X reporter cells (GFP+), carrying all possible SARS-CoV-2-derived peptides in the context of all 12 patient-specific HLA alleles (complexity up to 120.000 individual pMHC combinations) were co-cultured with 16.2A2 cells transduced with individual TCRs from patients. Responding (NFAT^+^) reporter cells were sorted, expanded and co-cultured 4 times. **C,** Individual responding reporter clones were isolated and re-analyzed by an additional co-culture. **D,** Sequences of the de-orphaned TCR chains, specific peptides and HLA restriction. **E**, 16.2X reporter cells carrying the MCR2-S_714-728_ or MCR2-N_221-242_ were analyzed on FACS for MCR2 expression (by aCD3 staining). **F**, 16.2X reporter cells carrying the MCR2-S_714-728_ or MCR2-S_714-728(F716I)_(top) and MCR2-N_221-242_ or MCR2-N_221-242(S235F)_ (bottom) were co-cultured with 16.2A2 cells transduced with TCR007 or TCR132 respectively and NFAT activation was measured on FACS.

Three CD4 T cell clones from severely affected patient C148 recognized peptides from the immunogenic SARS-CoV-2 proteins spike (S), membrane (M) and nucleocapsid (N), all presented by DRB1*07:01. T cells from other patients recognized peptides presented by other HLAs. For example, TCR091 from patient C141, reacted with several membrane glycoprotein-derived peptides, all presented by DRB1*11:01 and centered around the core epitope M_146-165_. In line with a high immunogenicity of this epitope, Peng et al.^3^ reported that 32% of patients contained T cells that recognize an overlapping peptide M_141-158_. Interestingly, two of the CD4 T cell specific peptides identified in our study (S_714-728_ and N_221-242_) were mutated in the SARS-CoV-2 B1.1.7 variant first identified in Britain, which rapidly spreads due to up to 70% increased transmission rates^23^. We transduced reporter cells with MCR2 carrying the WT and mutated S_714-728_ or N_221-242_ peptides (Fig.1E) and discovered that S_714-728_(T716I) was not recognized by the TCR007(Fig.1F). Recognition of N_221-242_ peptide was unaffected by the mutation, suggesting that Ser^236^ was not part of the minimal epitope (Fig.1F). We reasoned that the T716I mutation could abrogate TCR recognition by either of two mechanisms: the mutation might alter presentation on DRB1*07:01, or it could abolish TCR007 binding directly. To distinguish these two possibilities and to look more broadly at SARS-CoV-2 peptide presentation by different HLA alleles, we took advantage of our newly developed MEDi platform described below.

### MEDi, a mammalian epitope display platform based on MCR

Using the MCR system in our previous study, we identified the murine leukemia virus envelope protein-derived mutant peptide (MLVenv^S126R,D127V^, aka envRV) as being the cognate specificity of the mouse TIL-derived hybridoma, TILoma-1.4^18^. Interestingly, while MCR2 carrying envRV was expressed well on the surface of the reporter cells, the one carrying the nonmutated WT peptide (env) could not be detected, consistent with netMHCIIpan affinity predictions (Fig.2A). Given that MHC molecules without a bound peptide are very unstable^24^, this observation led to the hypothesis that peptides fitting well into the peptide-binding groove and therefore being efficiently presented by the MHC, will also effectively stabilize the MCR molecules on the surface of cells. In contrast, peptides not well presented by the MHC destabilize the MCR2 molecules and therefore little, if any, cell surface expression will be detected (Fig.2B). We therefore set out to test our hypothesis and cloned a number of peptides with biochemically tested I-Ab binding affinity ranging from 7.5nM to 10,000nM (Table 2), transduced them into our 16.2X reporter cell line, and determined the MCR2 expression by flow cytometry and staining with I-Ab and CD3 specific antibodies (Fig.2C and not shown). As expected from our previous study, there was a clear linear correlation between both stainings, but CD3 allowed a better separation of the positive and negative populations (Fig.2D). We therefore used anti-CD3 staining in all further MEDi analysis, with the added advantage of being MHC-agnostic and therefore universally usable with all mouse H-2 and human HLA haplotypes. We analyzed MCR2 expression dependence on peptide-I-Ab binding affinity by the mean fluorescence intensity (MFI) of CD3 staining. Consistent with our expectations, the MCR2s carrying peptides with a good I-Ab binding affinity were expressed on the surface at high levels, while MCR2s presenting low affinity peptides showed lower surface expression (Fig.2E). Peptides with an affinity below 1μM (IC_50_) are considered good MHC-binders and all MCRs carrying such peptides were expressed well on the cell surface. In addition, some peptides with lower MHC binding affinity appeared on the surface, indicating that linking peptides directly to the MHC beta chain stabilizes low-affinity peptide-MHC interactions. Being able to test the presentation of such peptides is important, as self-peptides known as targets in autoimmune diseases often bind MHC with low affinity^25^. 6 out of 6 peptides with an I-Ab binding affinity below 5μM (IC_50_) stabilized MCR2 surface expression, while for peptides with lower binding affinity, MCR expression was variable and generally much lower. Some of the MCR2s carrying peptides with an apparently low affinity (e.g., 8.39μM) were expressed on the surface at good levels, suggesting that additional factors apart from pure binding affinity (measured in vitro), regulate peptide-MHC interactions. Similarly, the envRV peptide could stabilize MCR2 expression, even if its I-Ab binding affinity was predicted by netMHCIIpan to be very low at 7.7μM and we needed to add high amounts of envRV peptide for in-vitro T cell stimulations by dendritic cells^18^.

**Table 2.**
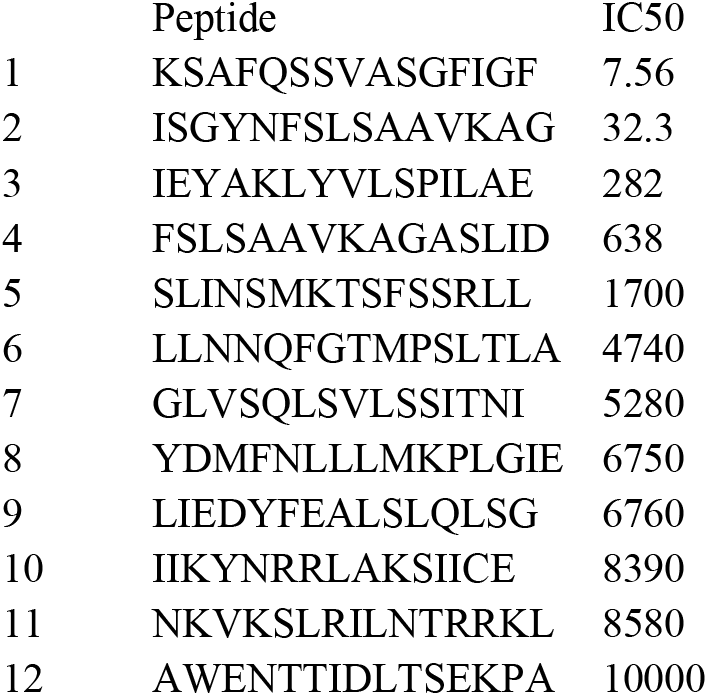
I-A^b^ presented peptides cloned in MCR2 vector

**Figure 2.**
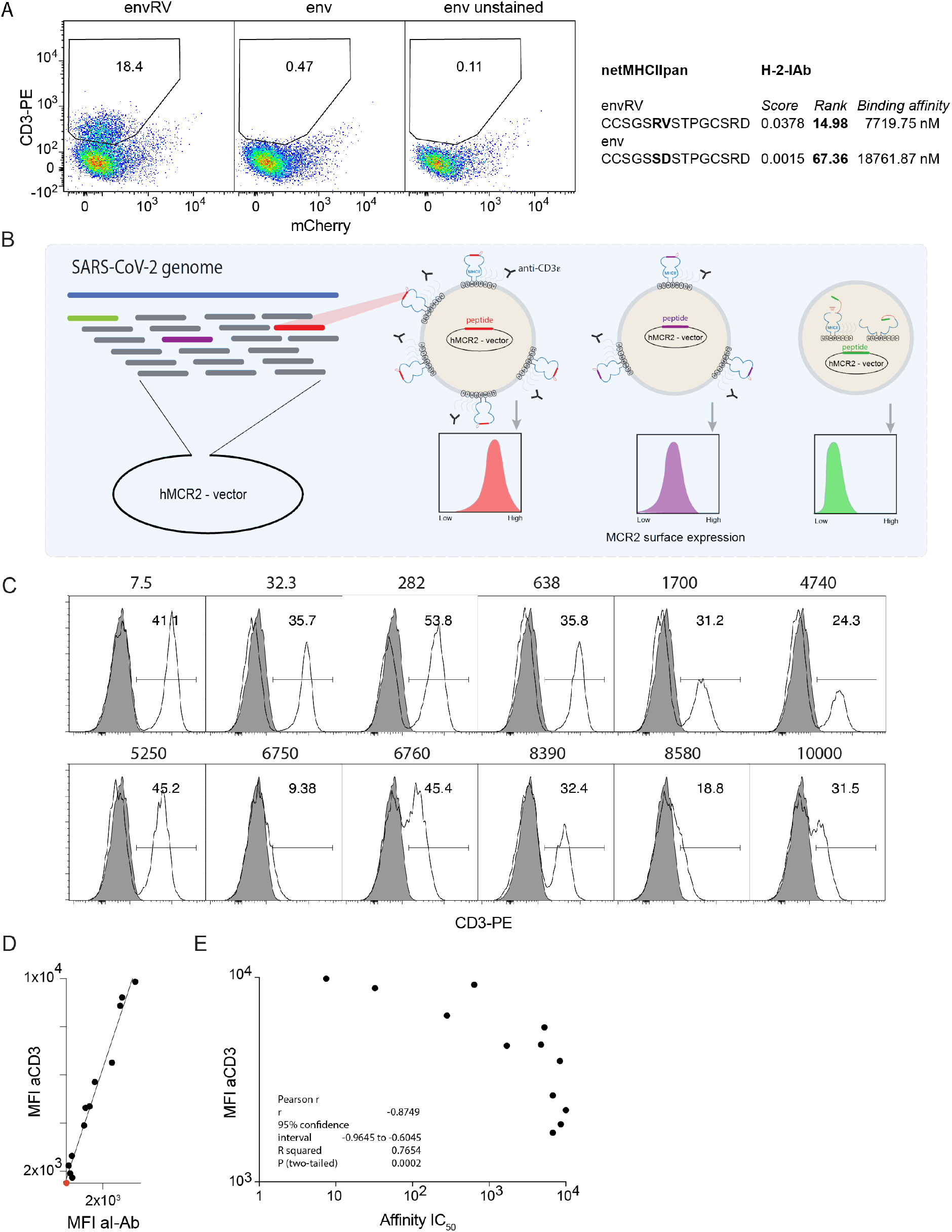
Cell surface expression of MCR2 depends on peptide-MHC binding. **AC**, Flow cytometric analysis of CD3e expression on 16.2c11 cells transduced with (**A**) MCR2-envRV or MCR2-env or (**C**) MCR2 constructs carrying peptides binding I-Ab with different affinity. **B,** Schematic representation of the peptide library cloning into the hMCR2 vector and of the MEDi principle. **D**, comparison of mean fluorescent intensity (MFI) of anti-CD3e and anti-I-Ab stainings. **E**, Correlation of MFI of anti-CD3e staining with the peptide MHC binding affinity of peptides carried by the MCR2.

### Analysis of SARS-CoV-2 peptides presentability by common HLA alleles

Considering the recent interest in SARS-CoV-2 T cell epitopes effectively presented across the possibly highest number of HLA alleles, we used MEDi to determine the presentability of all peptides encoded in the SARS-CoV-2 genome in the context of some of the most common HLA class II haplotypes. The critical role of CD4 T cell help in supporting B cell and CD8 T cell responses is undisputed and also crucial for COVID-19 protection^26,27,28^. However, a complete picture of the important MHC class II epitopes is missing, as they are more difficult to predict by computer algorithms than MHC class I ligands. To achieve a good resolution, we cloned all possible 15aa peptides derived from the SARS-CoV-2 genome (Fig.3A), shifted by 1 aa, into MCR2 vectors containing extracellular domains of the HLAs: DRB1*04:04, DRB1*07:01, DRB1*08:03, DRB1*11:01, DRB1*14:05, DRB1*15:01 and DPA1*02:02/DPB1*05:01 (Fig.3 and not shown). We transduced these libraries into the 16.2X reporter cell line, stained for CD3 and sorted the cells into 4 fractions (neg, low, mid and hi) based on the surface expression level of the MCR2 (Fig.3A). We then determined the peptides carried by the MCR2s in the different fractions by RT-PCR and deep sequencing. For each peptide a MEDi score was calculated using the formula sum_i[(Fr_index_i*Fr_count_i)/sum_i(Fr_count_i)] and plotted against the position of the starting amino acid of the peptide within the protein (see Methods). Figure 3B shows plots of the MEDi score moving average (MEDi-MA, average of 5 consecutive peptides) for the SARS-CoV-2 Spike peptide presentability by a set of 5 HLA alleles. Clearly, peptides derived from particular regions of the protein stabilize surface expression of the MCR better than others i.e., are being better presented by the MHC. Such peptides grouped in regions (“peaks/waves”), indicating that a core MHC-binding epitope was present in a number of peptides starting at several consecutive amino acids (Fig.3C,D). This observation is consistent with the fact that, owing to its open peptide-binding groove, MHC class II molecules present peptides of different length^7^. Usually the minimal MHC-binding core is composed of 9aa as shown by the commonly described binding motifs^29^, even if residues outside of it also contribute to the MHC-binding affinity^30^. As expected, MEDi graphs derived from these analyses showed a diverse presentation pattern. Each HLA molecule was unique, with regions of specific and promiscuous peptide presentation.

**Figure 3.**
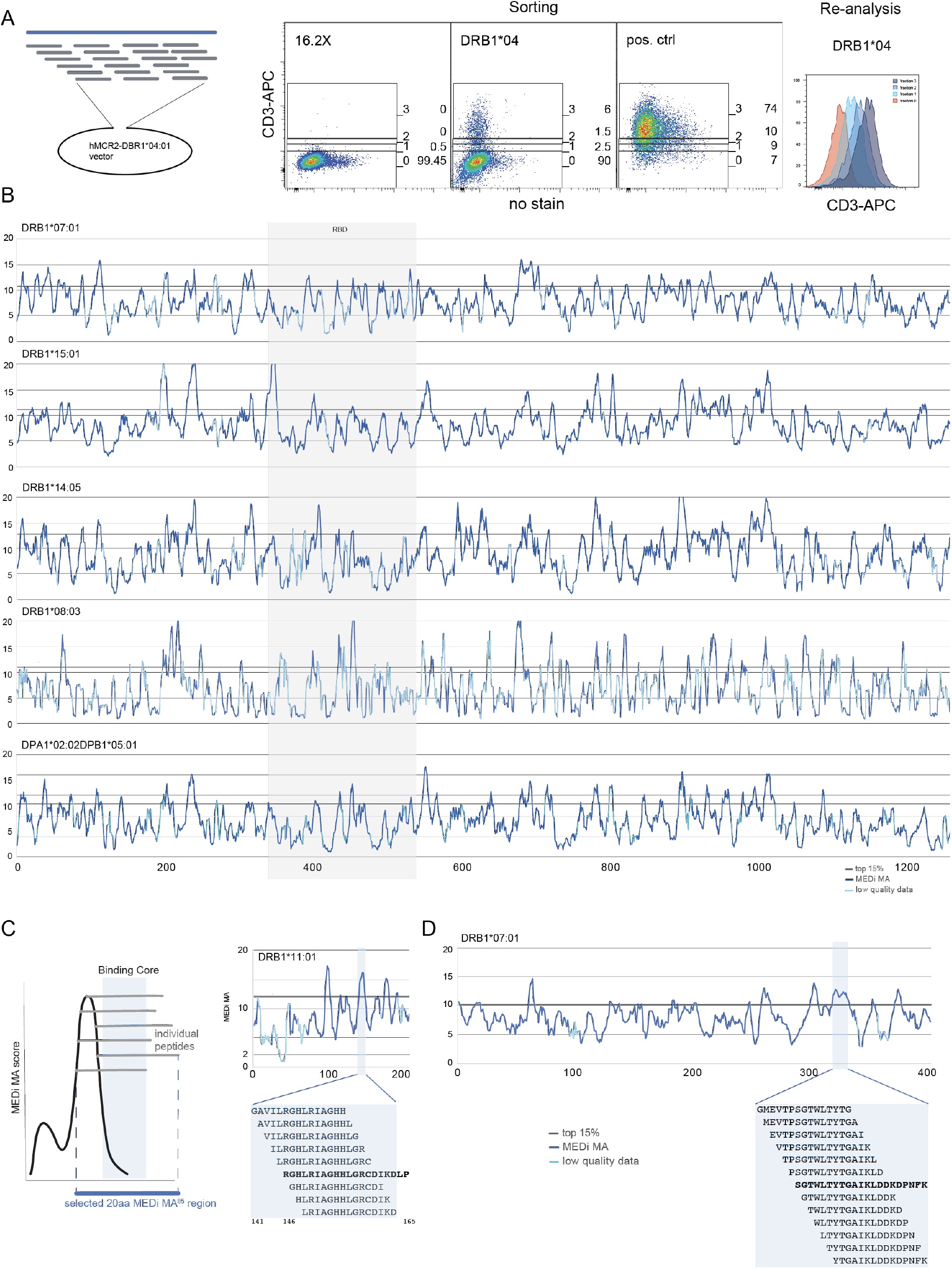
MEDi analysis of Spike peptide presentation by different HLAs. **A**, Example flow cytometric analysis and sorting of MCR2^+^ reporter cells, transduced with an MCR2 library and stained for CD3e. Based on the surface expression of the MCR2, four fractions (neg, low, mid and hi) were sorted and re-analyzed. Positive and negative controls are indicated. **B**, Shown are MEDi-MA score graphs for all Spike-derived peptides presented by 5 different HLAs (dark blue line). The light blue color indicates datapoints on the MEDi-MA graphs below the quality threshold (see M&M). **CD,** Schematics and interpretation of the MEDi graphs. MEDi analysis for the membrane (**C**) and nucleocapsid (**D**) proteins with indicated 15aa peptides falling into example MEDi-MA85 peaks. The extended peptides are recognized by COVID-19 specific TCRs analyzed in this study.

While the precision of this complex analysis is dependent on efficient cloning of all peptides and sufficient numbers of cells being sorted, there may be limitation in the number of sequencing reads per fraction and uneven coverage. To account for such data quality differences, we introduced a MEDi-MA quality metric composed of a minimal read count and the coefficient of variation (see M&M). From the graphs in figure 3B it is clear that the best results were obtained for DRB1*07:01 and DRB1*15:01 and DPA1*02:02/DPB1*05:01, while DRB1*14:05 and DRB1*08:03 showed lower quality. We therefore focused most of the MEDi platform testing on DRB1*07:01 and DRB1*15:01.

To distill the best HLA-binding peptides from this data, we selected peptides composed of the amino acids scoring above the 85^th^ percentile of all peptides from a given antigen (MEDi-MA85). As an example, in Table 3 we provide a list of potentially presentable peptides derived from the Spike protein and in the supplement we extend this analysis to all peptides derived from the SARS-CoV-2 genome in the context of 3 HLAs (supplementary data excel file S2). Of note, the spike list contains peptides greatly overlapping with the immunogenic peptides described in recent literature^3,4^.

**Table 3.**
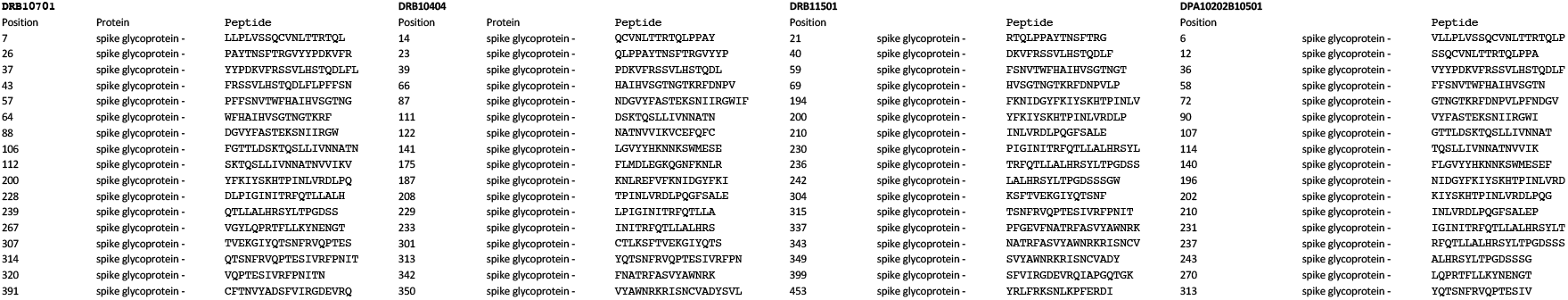
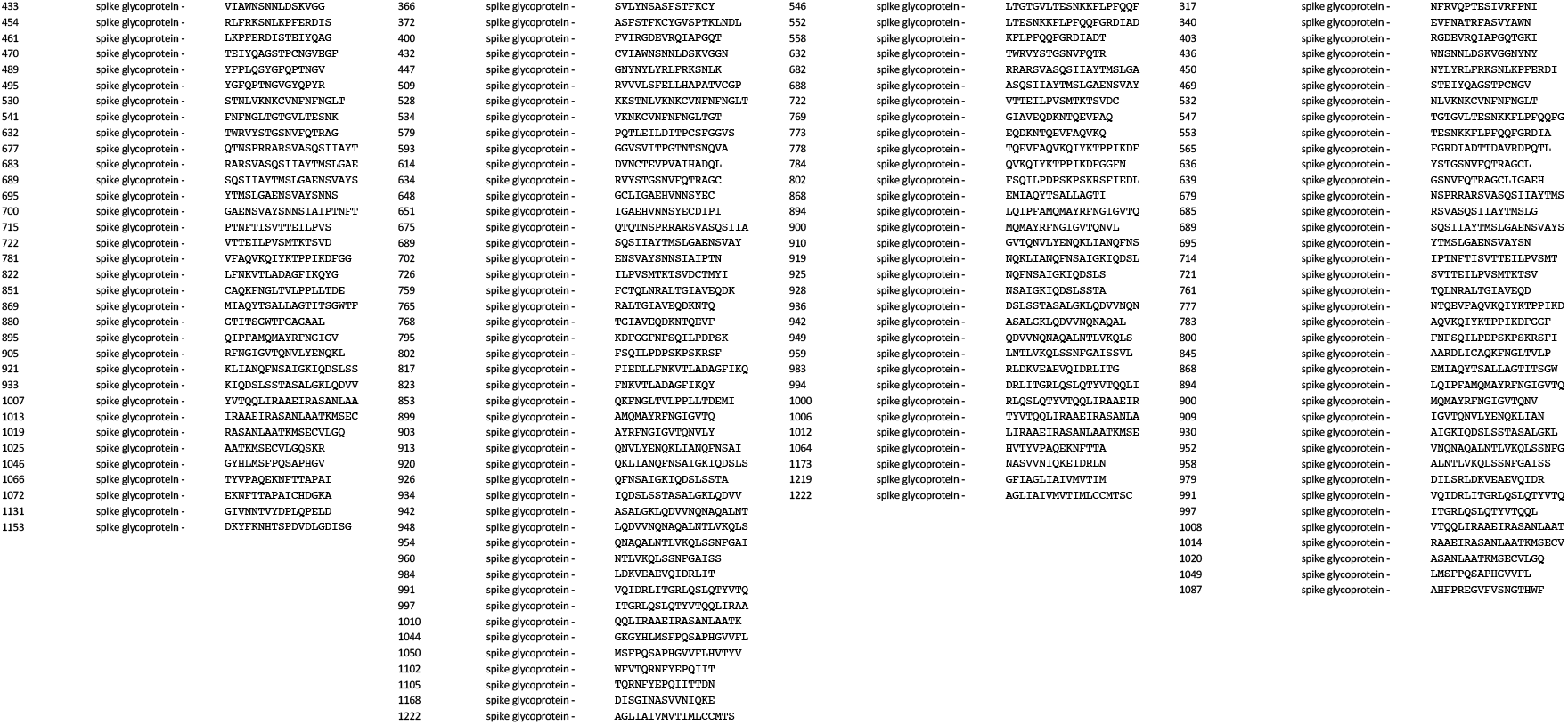
MEDi MA85 selected spike peptides.

### Validation of MEDi by a competitive peptide binding assay

Next, we analyzed peptides from the Spike protein in major MEDi-MA85 peaks for the presence of a binding motif and indeed found an enrichment of known^29^, appropriately spaced anchor residues in most of the selected peptides (Fig.4A DRB1*07:01 and Fig.S1A DRB1*15:01), validating our assay. Still, because tethering peptides to the MCR might stabilize some low affinity interactions not efficiently presented *in vivo*, we wanted to independently validate and quantify the HLA binding of the peptides defined by MEDi. To this end, we performed measurements of competitive peptide binding by fluorescence polarization^31^ for a set of Spike peptides for DRB1*07:01. We selected 33 peptides representing MEDi-MA peaks and 10 peptides representing valleys (Fig.4B) and considered peptides with IC_50_ below 10 μM as binders. When IC_50_ calculation was impossible due to very low peptide binding it was set arbitrarily to 20 μM.

**Figure 4.**
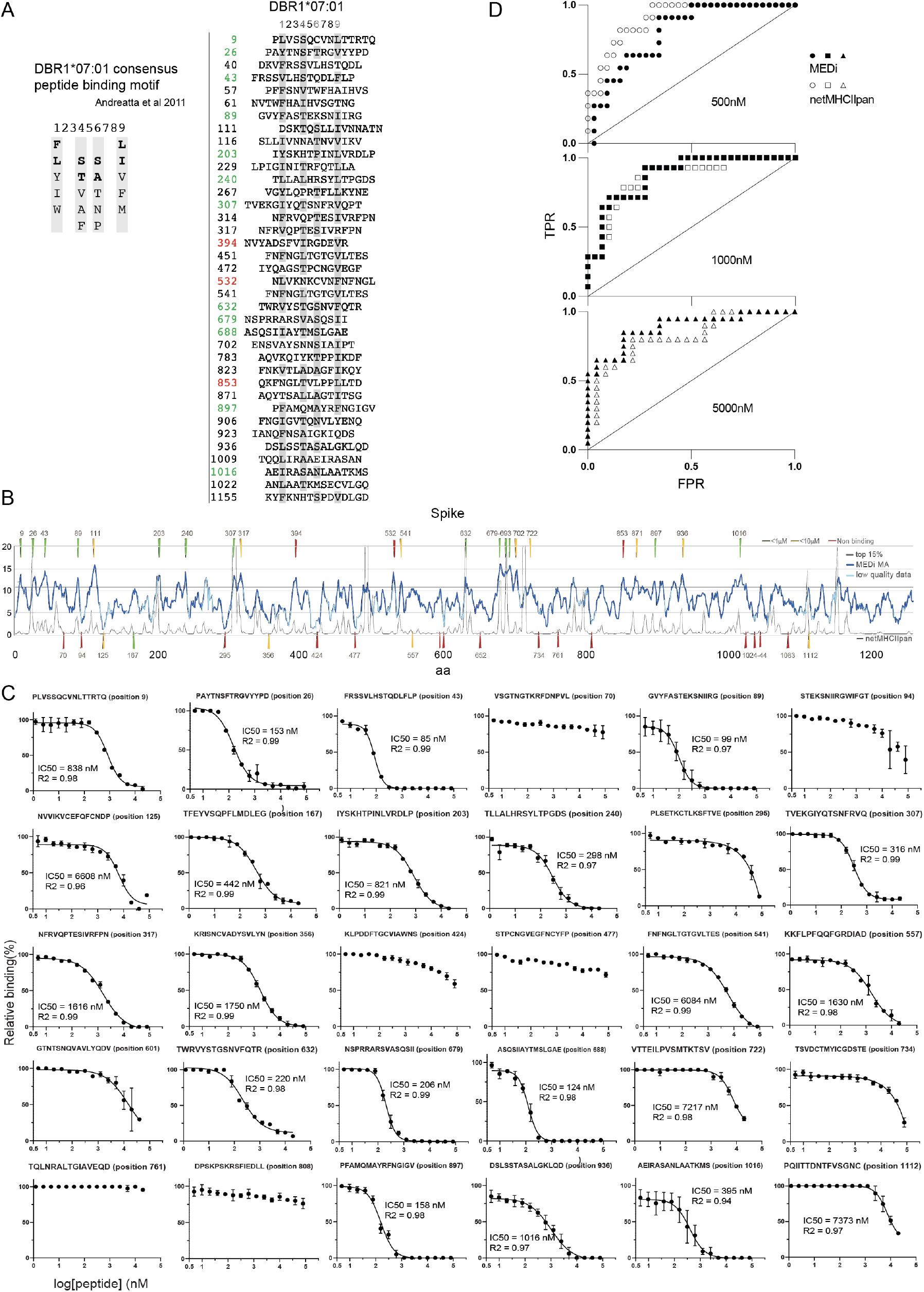
MEDi analysis of Spike peptide presentation by DRB1*07:01 compared to netMHCIIpan and MHC binding IC_50_. **A**, Sequence comparison of Spike peptides representative for the individual MEDi-MA85 peaks containing at least 3 peptides. Residues matching the HLA binding consensus are highlighted in grey **B**, MEDi-MA score graphs (dark blue line) for all Spike-derived peptides presented by DRB1*07:01. The light blue color indicates datapoints on the MEDi-MA graphs below the quality threshold (see M&M). Arrows indicate peptides chosen for HLA-binding IC_50_ calculation by the fluorescence polarization assay, color-coded dependent on the result of the binding assay. **C**, Results of the competitive peptide binding fluorescence polarization assay for individual peptides. IC_50_ and R^2^ values are shown. **D**, ROC curves of the MEDi-MA and netMHCIIpan scores qualifying peptides as HLA-binders. Calculations were done for peptides analyzed in C, positive binding thresholds at IC_50_ of 500nM, 1μM or 5μM.

20 out of the 23 peptides (87%) from the MEDi-MA85 peaks bound to the HLA with IC_50_ between 85nM to 10μM (Fig.4B,C), 13 of them below 1μM. From the remaining 10 peaks, three peptides bound to the HLA (IC_50_ 442nM, 1630nM and 7.3μM) but missed MEDi-MA85 cut-off by a small margin (Fig.4B,C) and for the rest no binding could be shown. Of note, confirmed HLA binding peptides contained 3 correct anchors, while the ones for which binding could not be confirmed had fewer. For peptides from the valleys, 2 out of 10 (20%) bound to the HLA with low affinity, while the rest did not bind (Fig.4C). This data set allowed us to analyze the ability of the MEDi assay to qualify peptides for HLA presentation and compare it to netMHCIIpan. We plotted receiver operating characteristic curves (ROC) for different presentation IC_50_ cut-offs (Fig.4D). The same analysis was done using the netMHCIIpan predicted EL rank for the same peptides (Fig.4D). Overall, the performance of both methods was comparable, with MEDi performing better for low affinity peptides (1μM and 5μM IC_50_ cut-offs: AUC 87.5% to 86% and AUC 88% to 82% respectively), while netMHCIIpan was better for the 500nM IC_50_ cut-off (AUC 89.8% to 80.2%).

Next, using 30 of the same peptides, we performed an unbiased analysis for DRB1*15:01 (Fig.S1). Here, because the peptides were chosen according to MEDi data for DRB1*07:01, most peptides corresponded to MEDi scores below the 85^th^ percentile threshold and were not in major peaks (FigS1AB). Therefore, they should not be well presented. Indeed, the majority did not bind the HLA with sufficient affinity (FigS1C). Nevertheless, 10 of the peptides were in peaks above the threshold and 7 bound to HLA. Reassuringly, the two peptides with the highest IC_50_ (122nM and 241nM) corresponded to 2 of the 5 highest MEDi-MA85 peaks and were on the top of the MEDi ranking. NetMHCIIpan placed them lower at the 3^rd^ and 30^th^ rank. On the other hand, 4 of 7 HLA binding peptides (IC50 from 310nM to 663nM) missed the MEDi 85^th^ percentile threshold. Two of them by a small margin, possibly due to low quality data in these regions. NetMHCIIpan also did not qualify 3 of the 7 binding peptides as good HLA binders but placed them at higher positions in the overall ranking (Fig.S1D). We conclude that both methods perform similarly for these common HLA alleles.

These results validate the MEDi platform as a means to select peptides highly presentable by an HLA allele. They also underscore the need for sufficient coverage in cell numbers and sequencing depth, for the precision of MEDi.

### MEDi indicates efficient presentation of immunogenic CD4 T cell epitopes

Next, we looked at the presentability of the CD4 T cell targets identified in our MCR screens. In line with the results presented in figure 1, MEDi data indicated good presentability of the TCR091 target peptide region by DRB1*11:01(Fig.5A and Fig.S2). Furthermore, consistent with high reactivity among patients shown by Peng et al.^3^, MEDi suggested presentability of this region by other HLA alleles like DRB1*04:04 and DRB1*15:01, and to a lower extent by DRB1*07:01 (Fig.5A). NetMHCIIpan only predicts DRB1*11:01, but the competitive peptide binding assay confirmed the MEDi results: DRB1*11:01 showed the highest IC_50_ (236nM-561nM), followed by DRB1*04:04(1.7-9.5μM) and DRB1*15:01(3.2-5.4μM) and the lowest DRB1*07:01 (4.7-14μM) (Fig.5A and Fig.S2). Even if these values do not precisely indicate differences in binding affinity, because the competing fluorescent peptides bind the HLAs with different affinity, the results highlight the advantages of MEDi over netMHCIIpan for discovering low-affinity peptide presentation.

Next, we analyzed MEDi scores of the other immunogenic peptides found in this study, and compared them to netMHCIIpan predictions (Fig.5BC, indicated in red). All of the CD4 T cell immunogenic peptides were found in the MEDi peaks, with S_955-971_ presented by DPA1*02:02/DPB1*05:01 and N_221-242_ presented by DRB1*14:05 being uniquely identified by MEDi. Also, 7 of the 8 peptides passed the MEDi-MA85 threshold. Only S_372-393_ showed a peak with lower MEDi scores, suggesting lower affinity HLA binding. Thus, selecting all immunogenic peptides for screening applications might require reduction of the MEDi threshold.

**Figure 5.**
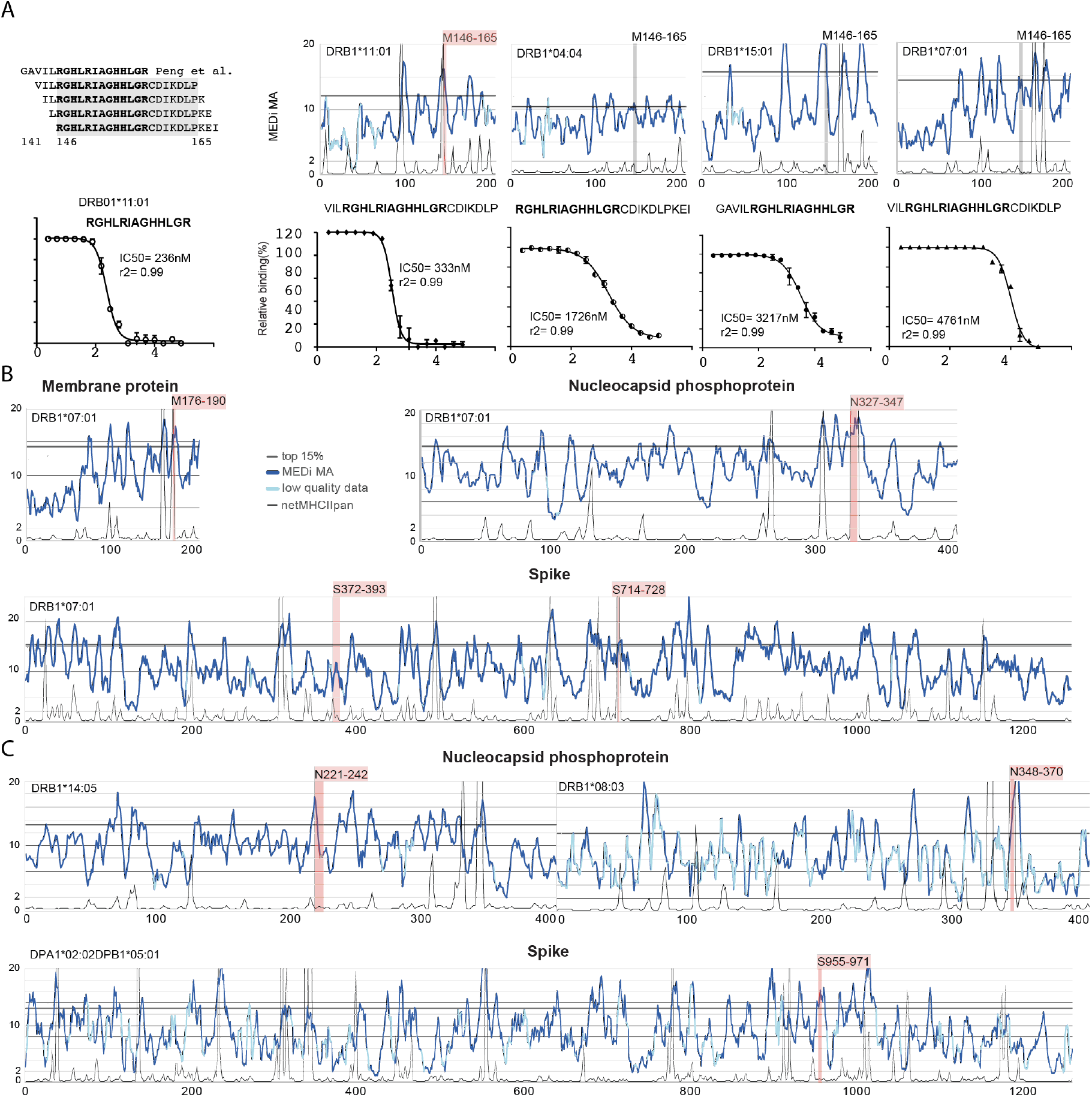
Presentation of immunogenic peptides by MEDi. **A**, MEDi-MA graphs (dark blue) for the membrane protein presented by the indicated HLA alleles. Results of the competitive peptide binding assay for the indicated peptides and alleles are shown below. M_146-165_ peptide (recognized by the TCR091 in the context of DRB1*11:01) is indicated in red for DRB1*11:01 and gray for the other alleles. **BC**, MEDi-MA score profiles compared to netMHCIIpan prediction scores (scaled to fit on the same plot: 20/Rank_EL, thin black) for the HLAs presenting CD4 T cell specific peptides found in this study (Fig.1D). MEDi-MA85 threshold is indicated as a black line, T cell specific peptides are indicated as red shades and netMHCIIpan threshold for weak presentation corresponds to the score of 2. Light blue color indicates datapoints on the MEDi-MA graphs below the quality threshold.

Taken together, these results indicate that MEDi selected peptides are enriched for immunogenic epitopes and that MEDi has an advantage over in-silico predictions for MHC class II alleles, where no high-quality mass spec results or other training data are available.

### MEDi also reveals candidate immune-escape mutants

These results illustrate the ability of our technology to determine presentable peptides for different HLA alleles. We therefore used MEDi to address the presentability of the new B1.1.7 virus-derived epitopes. As shown by the FACS analysis in figure 1E, mutations apparently did not affect cell surface expression levels of MCR2 presenting S_714-728_ or N_221-242_. We then systematically analyzed the effects of the B1.1.7 mutations on the presentation capacity of several HLA alleles. We generated MCR2 libraries containing all the mutation-overlapping 15mer peptides in the context of 8 different HLA alleles and performed MEDi analysis. As shown in figure 6, there was a notable HLA-dependent difference in B1.1.7 mutant peptide presentability. Several mutated peptides from nucleocapsid, ORF1a and ORF8 were inefficiently presented by DRB1*04:04, DRB1*04:01 and DPA1*02:02/DPB1*05:01 suggesting the possibility of immune escape of the virus in patients with these alleles. Particularly interesting in this regard were mutations I2230T and Y73C which disturbed the N-terminal hydrophobic amino acid stretches constituting a preferred binding motif for DRB1*04:04^29^ (Fig 6AB). Also, the spike HV69 deletion disturbed presentation by DRB1*07:01. The other alleles showed no difference between WT and mutated peptides, with a few exceptions where presentability of mutated peptides was enhanced. In particular, the spike D118H mutation appeared to stabilize binding of several peptides to DRB1*14:05, DRB1*15:01 and DRB1*07:01.

**Figure 6.**
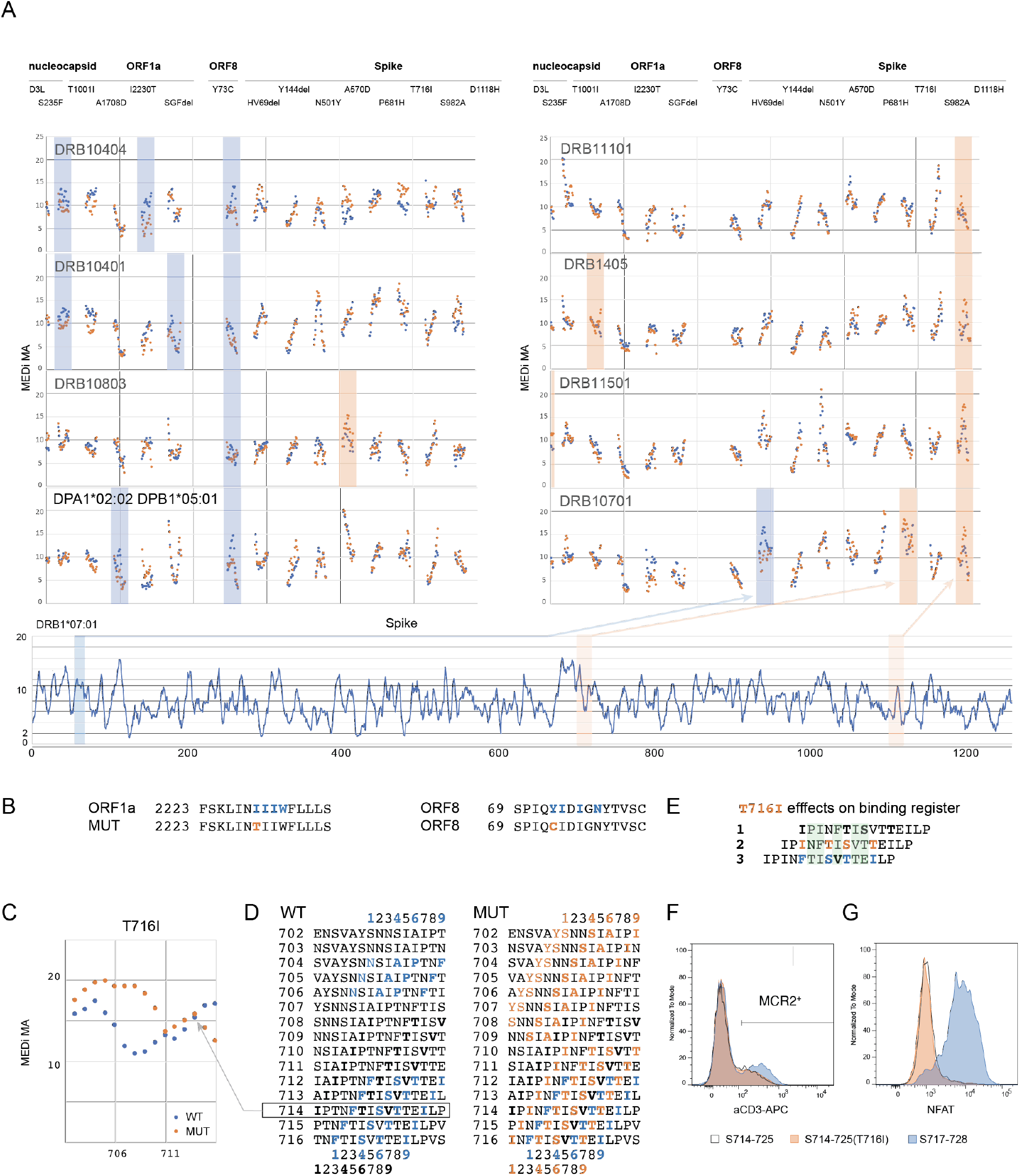
MEDi reveals candidate immune-escape mutants. **A**, Micro MCR2 libraries, containing all 15 (15aa long) peptides spanning the individual mutations were cloned for each HLA and transduced into the 16.2X reporter cells. Shown are individual MEDi-MA scores for the WT (blue dots) and mutated (orange dots) peptides. For context the MEDi scores for the full Spike protein are shown below. Blue and orange shaded squares indicate differences seen in all 3 repeat experiments. **B**, Example peptide sequences from ORF8 and ORF1a with indicated starting residues and the MHC binding motif for DRB1*04:04, are shown. **C**, Detailed view of the MEDi-MA scores for the WT and T716I Spike mutated peptides in the context of DRB1*07:01. **D**, 15mer peptides spanning the T716I mutation with indicated starting residues and the different DRB1*07:01binding motifs. **E**, S_714-728_ peptide sequences with indicated different binding registers forced by several DRB1*07:01 binding motifs present in the WT and/or mutated peptide. TCR facing residues are shown in green. **F**, FACS analysis and sorting of reporter cells transduced with DRB1*07:01-MCR2 carrying the 12mer peptides: S_714-725_, S_714-725(T716I)_ and S_717-728_. **G**, Reporter cells from F, were co-cultured with 16.2A2 cells transduced with TCR007 and NFAT activation was measured on FACS.

Furthermore, while MEDi-MA scores confirmed that the 15mer S_714-728(T716I)_ was presented as well as the WT, they also indicated that mutated peptides starting from Asp^702^ to Asn^710^ would be presented substantially better than WT (Fig.6C). Indeed, the T716I mutation introduced a perfect P9 anchor residue at position 716 complementing residues Tyr^707^/Ser^708^, Ser^711^ and Ala^713^ to form a good DRB1*07:01 binding motif (Fig.6D and 4A). Furthermore, T716I mutation introduced additional DRB1*07:01-binding motifs potentially allowing three different presentation registers for peptide S_714-728(T716I)_ (Fig.6E): first, comprising a weak HLA-binding motif starting at Ile^714^, with Thr^716^ directly accessible by the TCR; second starting with the mutated Ile^716^ as a new anchor residue; and third binding motif where the T716I mutation would presumably not be part of the minimal epitope for TCR007. Thus, mutation T716I could abrogate TCR recognition by either of two mechanisms: it could alter the presentation register on DRB1*07:01 (Fig.6E), or it could abolish TCR007 binding directly.

To answer this question, we cloned 12mer peptides S_714-725_, S_714-725(T716I)_ and S_717-728_ into the DRB1*07:01-MCR2 and cocultured MCR2^+^ reporter cells with TCR007 T cells. As shown in figure 6F all constructs were expressed well with S_717-728_ reaching the highest levels indicating best presentation. Intriguingly, TCR007 recognized S_717-728_, but not S_714-725_ (Fig.6G). This indicates that T716I abrogated TCR recognition of the S_714-728_ 15mer by altering peptide presentation rather than by directly affecting the TCR binding epitope. Change of the presentation register appears as the likely reason, but steric hindrance cannot be completely excluded at this point.

These results indicate several mechanisms of peptide presentation modulation and highlight the ability of the MEDi platform to decipher molecular details underlaying possible viral immune escape strategies. Comprehensive analyses of the arising viral mutants, studying the relation of presentability and immunogenicity, will be important for the development of future therapeutics.

## Discussion

Identifying the specificity of SARS-CoV-2 reactive lymphocytes is crucial for the fields of therapeutics and vaccine development. While protection from viral infections is mostly attributed to B cell and CD8 T cell effector functions, the balance between enabling and restricting them decides about life and death of the host. Thus, understanding the CD4 T cell reactivity, which orchestrates these responses, is critical.

While several methods exist to “de-orphan” TCRs^32^, MCR2 or SCT-based screens have the great advantage to provide fully decoded “immune synapses”, including TCR alpha and beta chain sequences, the recognized peptide and the presenting HLA. In this work we studied the reactivity of 109 enriched TCRs found in the bronchoalveolar lavages of acute COVID-19 patients and precisely analyzed their peptide specificity and HLA restriction. Highlighting the importance of CD4 T cells, we discovered that 8/47 (17.0%) of the CD4 TCRs and 3/63 (4.7%) of the CD8-derived TCRs recognized SARS-CoV-2 peptides in the lungs of SARS-CoV-2 patients.

The appearance of mutated SARS-CoV-2 with higher transmissibility raises important questions about the selective pressure that gave rise to the fitter variants and the role of immune escape in their evolution. While viral escape from antibody-mediated neutralization has been well documented for many diseases^33,34^, much less is known about a potential selective pressure to evade T cell reactivity. Understanding HLA presentation and TCR recognition of mutant and WT epitopes is critical in this regard. Recently, Redd et al^35^ found only one of the mutations to overlap with CD8 epitopes and it was considered unlikely to affect TCR recognition. In this study we found that several mutations present in the emerging SARS-CoV-2 B1.1.7 strain reduced presentability of the affected peptides by several HLA class II alleles. Furthermore, a CD4 T cell targeted 15mer peptide was affected by the spike T716I mutation and was no longer recognized by the cognate TCR. We tested two different mechanisms of escape from CD4 T cell recognition and found that Thr^716^ was not directly bound by the TCR, but that the T716I mutation altered peptide presentation, presumably leading to presentation in a different register. This evasion strategy would affect all T cells recognizing this peptide, so the T716I mutation might provide a bigger advantage for the virus than appreciated so far. But how likely is it to be relevant in vivo if it resides outside the main nonameric HLA-binding core? Previous studies have shown the influence of so-called peptide-flanking residues (PFRs) on HLA-binding and subsequent T cell recognition^36^. Therefore, given the optimal peptide length for MHC class II being 18-20 amino acids^30^, it is very likely that most peptides, comprising the HLA binding core starting at Phe^718^, will include the Thr/Ile^716^.

Methods such as detection of natural peptides eluted from MHC by mass spectrometry, or in silico prediction, have both contributed to the understanding of peptide presentation. However, they do not provide complete presentability landscapes across multiple alleles, owing to the low sensitivity of mass spectroscopy and varying, often limited, accuracy of the in-silico methods depending on the HLA allele. With MEDi, we provide an alternative/complementary approach, in the form of a novel mammalian epitope display platform. It is based on functional cell surface expression of the MCR2 molecules. It is HLA agnostic thanks to the association of MCR2 with the CD3 chains and allows unbiased, fast and affordable testing of all antigen-derived peptides for their ability to stabilize MHC(MCR2) expression on the surface. We show proof of concept experiments, indicating that antigenic peptides usually reside within the MEDi score high regions, provide a list of presentable SARS-CoV-2 peptides for several different HLA alleles and describe different possibilities for viral immune evasion.

Testing MEDi-MA85 selected peptides by a competitive fluorescence polarization assay has confirmed HLA binding for most peptides. Interestingly, some of the peptides bound the HLA with low, or very low IC_50_, suggesting low affinity. However, the competitive binding assay is an in-vitro assay with recombinant HLAs and its limitations have to be also considered. Nevertheless, it is expected that direct tethering of peptides to the HLA stabilizes low-affinity interactions. This allows testing of such potentially short-lived peptide-HLA complexes for presentation and T cell reactivity. The key question was whether MEDi selected peptides are relevant in vivo. We therefore analyzed whether immunogenic epitopes correspond to MEDi scoring peaks. Indeed, all 8 HLA class II restricted immunogenic peptides were identified by MEDi. NetMHCIIpan missed two of them, but overall performed well for the common HLA alleles used in this study. MEDi has the advantage of being easily scalable to the thousands of alleles present in man, and to describe peptide presentability by patient-specific HLA alleles for which no good training data are available. Consistently, the immunogenic spike S_955-969_ peptide presented by DPA1*02:02/DPB1*05:01 and N_221-242_ presented by DRB1*14:05, both MEDi high, were not well predicted by netMHCIIpan. Furthermore, with MEDi we can quickly provide the presentability information for any immunogenic peptide across multiple HLA alleles. This is exemplified in this study by the very immunogenic membrane protein peptide M_146-165_, recognized by TCR091 in the context of DRB1*11:01 and shown by MEDi to be also presentable by several other HLAs, also not predicted by netMHCIIpan. However, the information gained from MEDi can support further training of predictive models similar to Rappazzo et al^13^.

Notably, when performing CD8 T cell screening experiments with SCTz, we realized that, in contrast to the MCR2, SCTz fused to different peptide libraries were all expressed on the cell surface, irrespective of correct folding (data not shown). Unfortunately, monoclonal antibodies specifically distinguishing the native HLA conformation from the misfolded one are not available for many alleles, precluding the broad use of SCT for MEDi-type applications at present.

The results presented in this study validate the MEDi platform and provide insights into the molecular mechanisms of SARS-CoV-2 peptide presentation and potential escape from T cell recognition. MEDi should help closing the gaps in peptide-presentation landscape for thousands of HLA alleles and be useful for the development of novel therapeutical approaches beyond prevention of COVID-19 or treatment of SARS-CoV-2 patients.

## Material and Methods

### Cell lines, molecular cloning and retroviral transduction

Cell lines, molecular cloning procedures and retroviral transduction used in this study were described previously^18^

### MEDi procedure and score calculation

Libraries carrying 15-amino acid (aa) long peptides, spanning the entire sequences of the SARS-CoV-2 virus, were cloned as oligonucleotides (Twist) into MCR vectors carrying different HLA alleles. 16.2X reporter cell line was transduced with these libraries and surface expression of the MCR molecules was analyzed by flow cytometry. Four fractions were sorted: Fr.0 (cells expressing no detectable MCRs on the surface), Fr.1 (cells expressing low levels of MCRs), Fr.2 (cells expressing intermediate levels of MCRs), Fr.3 (cells expressing high levels of MCRs) – see figure 3A. Peptides carried by the MCRs from sorted cells were amplified from cDNA by RT-PCR using the peptide flanking regions and sequenced on a miniSeq (Illumina). Sequences from the Illumina output files were trimmed, merged and translated with the help of the CLC genomic workbench program. Counting was done with FilemakerPro18 and all further analysis with Excel (Microsoft).

The individual peptide counts in each fraction were normalized to the total counts in the fraction. For each peptide a MEDi score was calculated with the following formula: sum_i [(Fr_index_i*Fr_count_i)/sum_i(Fr_count_i)]. Fr_indexes: Fr1=1, Fr2=2, Fr3=4, Fr4=28. MEDi-MA was calculated by averaging MEDi scores for 5 peptides (-2/-1/0/1/2) and assigned to the middle(0) peptide. MEDi-MA85 indicates the threshold calculated as the 85^th^ percentile of the MEDi-MA score for the individual protein.

### MEDi MA score quality threshold

MEDi MA score for a given peptide was considered of good quality if at least 40 reads were collected for a peptide and the MEDi MA value had a coefficient of variation (CV=Std.Deviation/Average) lower than 0.75.

### Local maximum of MEDi-MA peak definition

Local maximum of 7 MEDi-MA scores was determined (-3/-2/-1/0/1/2/3) and assigned to the middle(0) peptide.

### MCR2 screening

Libraries were generated by cloning all SARS-CoV2-derived peptides in MCR2 molecules carrying the complete viral genome in 23mers shifted by 1 aa. For screening we pooled the libraries at equal ratios, generating a combined patient-specific library of roughly 120.000 different peptide-MCR2 combinations (Table 1).

MCR2 screening was performed as described previously^18^. Briefly, MCR2 expressing 16.2X cells have been co-cultured with cell clones expressing one specific TCR selected from Liao et al. in a ratio of 1:5 to 1:10. Cells were mixed and co-cultured for 8-12 hours in a standard tissue culture medium, in the presence of 13ug/ml anti-mouse FasL antibodies (BioXcell) to inhibit induction of cell death during incubation. After harvesting, reporter cells positive for NFAT signaling have been sorted on a BD FACS Aria Fusion Cell Sorter as bulk or into 96 well plates for further expansion. On average 4-5 enrichment round per TCR have been performed before single reporter cells have been sorted. Expanded single cells were harvested, DNA was isolated (Kapa Express Extract) followed by sanger sequencing of the MCR2 alpha and beta chain including the linked antigen. Whenever overlapping peptides were found in the screen (e.g fig5A), in figure 1D, we listed the common part as the specific peptide recognized by the TCR.

### Single Chain Trimer-zeta (SCTz) screening

Single chain trimers of class I HLAs of all seven patients have been generated by linking the leader sequence, epitope, b2m and HLA alpha chain with 3 x G4S linkers and addition of the intracellular domain from the CD247(zeta-chain) molecules. Each alpha chain was modified/ mutated to open the groove of class I by introducing the Y84A mutation in every alpha chain^37^. For SCTz screening we again used libraries covering the whole SARS-CoV-2 genome with 10mers shifted by 1aa, cloned as oligonucleotides into the SCTz vectors.

### Fluorescence Polarization Assay

#### Fluorescence Polarization Assay

The MHC II α- and β-chain extracellular domains were recombinantly expressed with C-terminal Myc and His tag sequences, respectively. For DRB1*15:01 the Myc tag was replaced with a V5 tag. The N-terminus of the β-chain was fused to CLIP peptide followed by a flexible Factor Xa-cleavable linker. Both α- and β-chains were co-expressed in CHO cells and secreted into the expression medium as a stable CLIP-loaded heterodimer. Heterodimerization of the α- and β-chains of DRB1*07:01 and DRB1*1501 was forced using a fusion of an engineered human IgG1-Fc protein to each chain^38^. Following CHO expression, the heterodimer was purified by immobilized metal ion affinity chromatography and size exclusion chromatography (SEC). The fluorescence polarization assay was performed as described in^31^ with few modifications. Following Factor Xa cleavage, 100 nM of HLA were incubated overnight with 25 nM fluorescent probe and various concentrations of the indicated peptide competitor in 100 mM Sodium citrate pH 5.5, 100 mM NaCl, 0.1% octylglucoside and 1x protease inhibitors (SigmaFast) at 30° C. The fluorescent probe for DRB1*04:04, DRB1*07:01 and DRB1*11:01 was PRFV(K/Alexa488)QNTLRLAT. The fluorescent probe for DRB1*15:01 was ENPVVHFF(C/Alexa488Mal)NIVTPR.

## Supplementary Figures

**FigureS1.**
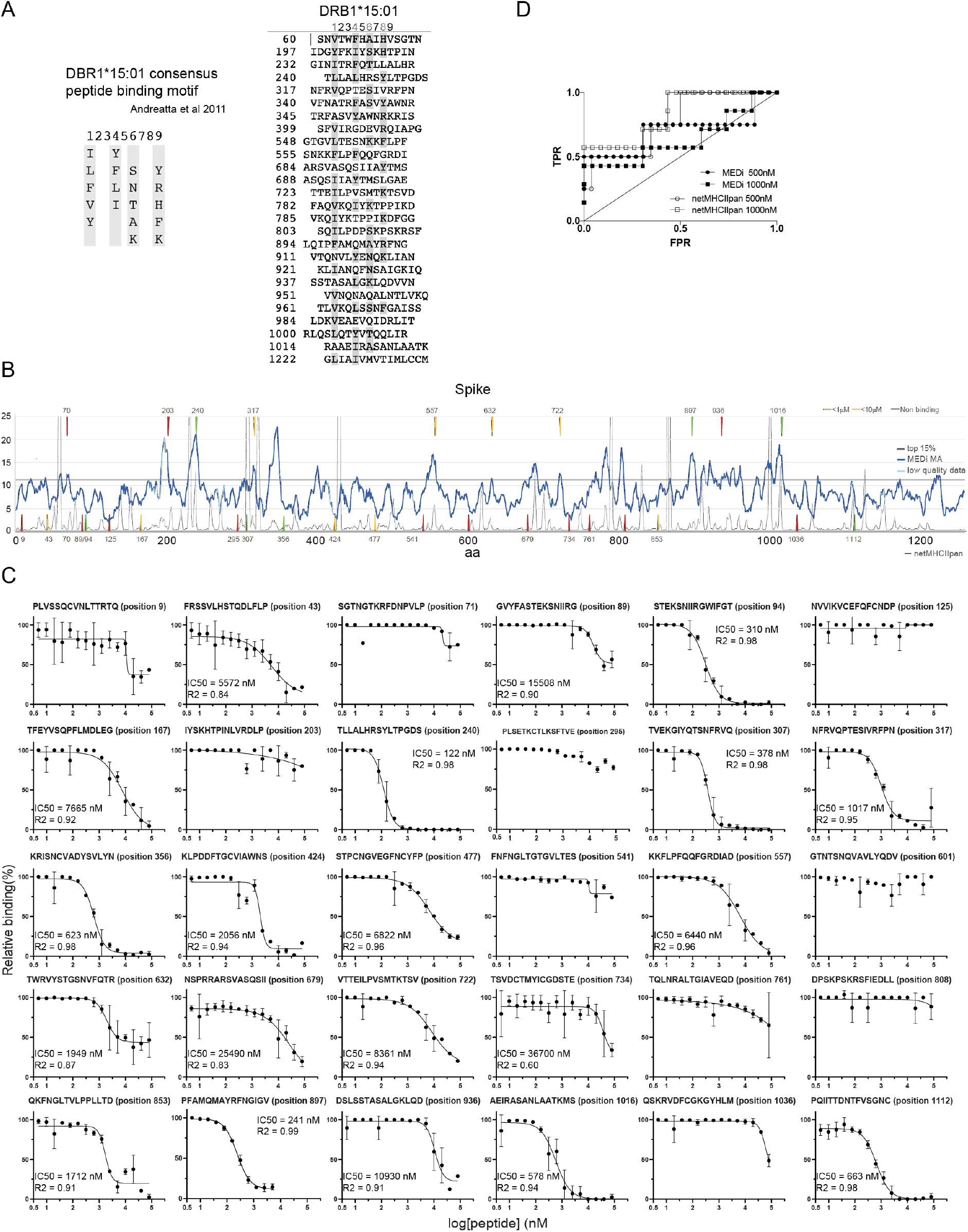

**Suppl.Figure S2.**
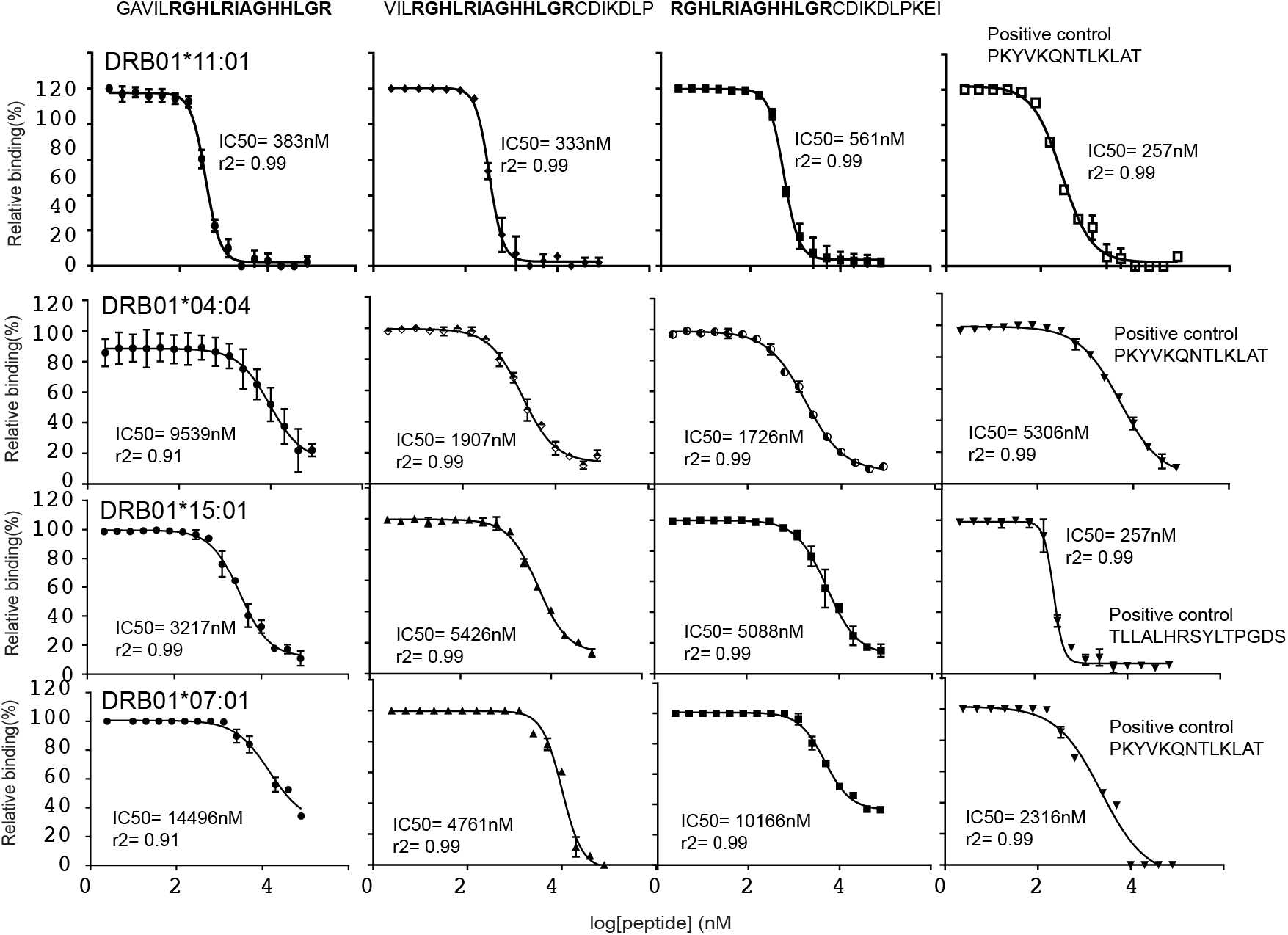

## Supplementary Files

excel file S1: TCR data.

excel file S2: Presentable peptides derived from the SARS-CoV-2 genome by MEDi MA85.

## Author contribution

**Conceptualization**, F.J.O., and J.K.; **Methodology**, F.J.O., M.K., J.S. and J.K.; **Investigation**, F.J.O., F.R., S.H., C.H.L., N.C., G.M., Y.H., R.W., O.I., P.P., K.T., and J.K.; **Resources**, M.K., J.S. **Writing – Original Draft**, J.K.; **Writing – Review & Editing,** F.J.O., M.K., J.S. and J.K; **Supervision,** F.J.O., J.S. and J.K.

## Acknowledgements

We thank Dr. D.Acker (Repertoire Immune Medicine) for discussions related to MEDi score analysis. We thank Dr. A.Coyle and Dr. T.Harris (Repertoire Immune Medicine) for critical reading of the manuscript. A.Schütz and Dr. M.Kisielow (ETHZ Flow Cytometry Core Facility) for help with cell sorting, D.Kollegger and J.Meier for technical assistance and Dr. M.Loi (Tepthera Ltd, Zürich) for discussions.

