## Supplemental file S1 for "High resolution profiling of MHC-II peptide presentation capacity, by Mammalian Epitope Display, reveals SARS-CoV-2 targets for CD4 T cells and mechanisms of immune-escape"

| patient | original_id | summary | t_cell_type | name of file for x-ref | count | type | TRAV | TRAJ | alpha junction | TRBV | TRBJ | beta junction |
|---|---|---|---|---|---|---|---|---|---|---|---|---|
| C141 | clonotype008 | No hit | CD4+ T cells | TCR-Liao-C141-008 | 13 | CD4 | TRAV12-1*01 | TRAJ52*01 | CVVTVNAGGTSYGKLTF | TRBV9*01 | TRBJ1-2*01 | CASSAQDGYTF |
| C141 | clonotype039a | No hit | CD4+ T cells | TCR-Liao-C141-039a | 4 | CD4 | TRAV13-1*01 | TRAJ39*01 | CAASIGNAGNMLTF | TRBV30*01 | TRBJ2-2*01 | CAWRIRDRVTGELFF |
| C141 | clonotype039b | No hit | CD4+ T cells | TCR-Liao-C141-039b | 4 | CD4 | TRAV24*01 | TRAJ45*01 | CASPNSGGGADGLTF | TRBV30*01 | TRBJ2-2*01 | CAWRIRDRVTGELFF |
| C141 | clonotype045 | No hit | CD4+ T cells | TCR-Liao-C141-045 | 4 | CD4 | TRAV13-2*01 | TRAJ26*01 | CAENNYGQNFVF | TRBV7-9*01 | TRBJ1-1*01 | CASSFGGSNTEAFF |
| C141 | clonotype062 | No hit | CD4+ T cells | TCR-Liao-C141-062 | 3 | CD4 | TRAV10*01 | TRAJ49*01 | CVVSRTGNQFYF | TRBV5-1*01 | TRBJ2-7*01 | CASSGGQEGGEQYF |
| C141 | clonotype091 | Hit | CD4+ T cells | TCR-Liao-C141-091 | 2 | CD4 | TRAV6*01 | TRAJ15*01 | CALDKQAGTALIF | TRBV20-1*01 | TRBJ2-2*01 | CSARAEGQSTGELFF |
| C141 | clonotype119 | No hit | CD4+ T cells | TCR-Liao-C141-119 | 2 | CD4 | TRAV13-1*01 | TRAJ31*01 | CAASWGNNARLMF | TRBV20-1*01 | TRBJ2-7*01 | CSARDRDYPYEQYF |
| C141 | clonotype150 | No hit | CD4+ T cells | TCR-Liao-C141-150 | 1 | CD4 | TRAV27*01 | TRAJ47*01 | CAGVRYGNKLVF | TRBV4-2*01 | TRBJ2-5*01 | CASSQEAGAPETQYF |
| C141 | clonotype156 | No hit | CD4+ T cells | TCR-Liao-C141-156 | 1 | CD4 | TRAV8-2*01 | TRAJ44*01 | CVVSRGGTASKLTF | TRBV5-1*01 | TRBJ1-2*01 | CASSFREGTYGYTF |
| C141 | clonotype418 | No hit | CD4+ T cells | TCR-Liao-C141-418 | 1 | CD4 | TRAV20*01 | TRAJ6*01 | CAVQAGGSYIPTF | TRBV3-1*01 | TRBJ1-2*01 | CASSQDHNGYTF |
| C141 | clonotype001 | No hit | CD8+ T cells | TCR-Liao-C141-001 | 26 | CD8 | TRAV13-1*01 | TRAJ6*01 | CAASTSGGSYIPTF | TRBV12-4*01 | TRBJ2-5*01 | CASSPSLLSGRARETQYF |
| C141 | clonotype003 | No hit | CD8+ T cells | TCR-Liao-C141-003 | 19 | CD8 | TRAV14/DV4*01 | TRAJ33*01 | CAMREDDSNYQLIW | TRBV20-1*01 | TRBJ2-1*01 | CSATVAGGSYNEQFF |
| C141 | clonotype004 | No hit | CD8+ T cells | TCR-Liao-C141-004 | 15 | CD8 | TRAV27*01 | TRAJ33*01 | CAGEGNSNYQLIW | TRBV12-4*01 | TRBJ1-1*01 | CASSLRATDEAFF |
| C141 | clonotype006a | No hit | CD8+ T cells | TCR-Liao-C141-006a | 15 | CD8 | TRAV12-3*01 | TRAJ49*01 | CAHRRNQFYF | TRBV7-8*01 | TRBJ2-1*01 | CASSLGYTEQFF |
| C141 | clonotype007 | No hit | CD8+ T cells | TCR-Liao-C141-007 | 14 | CD8 | TRAV13-1*01 | TRAJ30*01 | CAASPLRGNRDDKIIF | TRBV19*01 | TRBJ2-3*01 | CASSINPDFTDTQYF |
| C141 | clonotype009 | No hit | CD8+ T cells | TCR-Liao-C141-009 | 12 | CD8 | TRAV19*01 | TRAJ4*01 | CALSGFSGGYNKLIF | TRBV19*01 | TRBJ1-1*01 | CASKGVGAEDTEAFF |
| C141 | clonotype010 | No hit | CD8+ T cells | TCR-Liao-C141-010 | 12 | CD8 | TRAV12-2*01 | TRAJ4*01 | CAVNLFIGGYNKLIF | TRBV15*01 | TRBJ2-2*01 | CATSRAGTYNTGELFF |
| C142 | clonotype094 | Hit | CD4+ T cells | TCR-Liao-C142-094 | 2 | CD4 | TRAV41*01 | TRAJ47*01 | CAVRSRGNKLVF | TRBV7-2*01 | TRBJ1-1*01 | CASSQTFAEAFF |
| C142 | clonotype143 | No hit | CD4+ T cells | TCR-Liao-C142-143 | 1 | CD4 | TRAV9-2*01 | TRAJ57*01 | CARREMGGSEKLVF | TRBV3-1*01 | TRBJ2-1*01 | CASSPHRNEQFF |
| C142 | clonotype330 | No hit | CD4+ T cells | TCR-Liao-C142-330 | 1 | CD4 | TRAV19*01 | TRAJ57*01 | CALSEAGGGSEKLVF | TRBV5-1*01 | TRBJ2-7*01 | CASSLDTRQDYSYEQYF |
| C142 | clonotype335 | No hit | CD4+ T cells | TCR-Liao-C142-335 | 1 | CD4 | TRAV21*01 | TRAJ8*01 | CAVNRNTGFQKLVF | TRBV7-2*01 | TRBJ2-1*01 | CASSLRGLSSYNEQFF |
| C142 | clonotype340 | No hit | CD4+ T cells | TCR-Liao-C142-340 | 1 | CD4 | TRAV5*01 | TRAJ31*01 | CAVGGARLMF | TRBV6-1*01 | TRBJ2-1*01 | CASRRGEAGGRARYEQFF |
| C142 | clonotype001 | No hit | CD8+ T cells | TCR-Liao-C142-001 | 22 | CD8 | TRAV17*01 | TRAJ10*01 | CATVDILTGGGNKLTF | TRBV10-3*01 | TRBJ2-1*01 | CAISEWGEGPNEQFF |
| C142 | clonotype002 | No hit | CD8+ T cells | TCR-Liao-C142-002 | 20 | CD8 | TRAV14/DV4*01 | TRAJ29*01 | CAMRLNSGNTPLVF | TRBV27*01 | TRBJ2-1*01 | CASSLTRGLTYNEQFF |
| C142 | clonotype003 | No hit | CD8+ T cells | TCR-Liao-C142-003 | 19 | CD8 | TRAV5*01 | TRAJ6*01 | CADAPASGGSYIPTF | TRBV5-4*01 | TRBJ2-3*01 | CASSPRGENPDTQYF |
| C142 | clonotype004 | No hit | CD8+ T cells | TCR-Liao-C142-004 | 18 | CD8 | TRAV12-2*01 | TRAJ10*01 | CASYTGGGNKLTF | TRBV30*01 | TRBJ1-2*01 | CAWSVGGVMNGYTF |
| C142 | clonotype005 | No hit | CD8+ T cells | TCR-Liao-C142-005 | 18 | CD8 | TRAV10*01 | TRAJ45*01 | CVVRGGSGGGADGLTF | TRBV9*01 | TRBJ2-2*01 | CASSVSRLAGPNTGELFF |

| | | | | | | | | | | | | |
|---|---|---|---|---|---|---|---|---|---|---|---|---|
| C142 | clonotype006 | No hit | CD8+ T cells | TCR-Liao-C142-006 | 18.5 | CD8 | TRAV12-1*01 | TRAJ43*01 | CVVHDMRF | TRBV12-4*01 | TRBJ2-1*01 | CASSLDVDNEQFF |
| C142 | clonotype007 | No hit | CD8+ T cells | TCR-Liao-C142-007 | 17 | CD8 | TRAV17*01 | TRAJ10*01 | CATALLFTGGGNKLTF | TRBV28*01 | TRBJ1-5*01 | CASSLYDGEEHSNQPQHF |
| C142 | clonotype008 | No hit | CD8+ T cells | TCR-Liao-C142-008 | 13 | CD8 | TRAV17*01 | TRAJ10*01 | CATVDILTGGGNKLTF | TRBV10-3*01 | TRBJ1-2*01 | CAISDPASRGSPNGYTF |
| C142 | clonotype016 | No hit | CD8+ T cells | TCR-Liao-C142-016 | 7 | CD8 | TRAV9-2*01 | TRAJ12*01 | CALWMDSSYKLIF | TRBV28*01 | TRBJ2-7*01 | CASSPTGTGARGYEQYF |
| C143 | clonotype004 | No hit | CD4+ T cells | TCR-Liao-C143-004 | 18 | CD4 | TRAV12-1*01 | TRAJ5*01 | CVVNYYGDTGRRALTF | TRBV28*01 | TRBJ1-5*01 | CASSPMWGGTSLSNQPQHF |
| C143 | clonotype006 | No hit | CD4+ T cells | TCR-Liao-C143-006 | 14 | CD4 | TRAV13-1*01 | TRAJ41*01 | CASGYALNF | TRBV11-2*01 | TRBJ2-7*01 | CASSLGTGGVYEQYF |
| C143 | clonotype007 | Hit | CD4+ T cells | TCR-Liao-C143-007 | 13 | CD4 | TRAV12-1*01 | TRAJ53*01 | CVVNTPGDSGGSNYKLTF | TRBV7-8*01 | TRBJ2-2*01 | CASRRGTGGHTGELFF |
| C143 | clonotype011 | No hit | CD4+ T cells | TCR-Liao-C143-011 | 8 | CD4 | TRAV17*01 | TRAJ54*01 | CATDAAQKLVF | TRBV20-1*01 | TRBJ2-1*01 | CSARTSGRVRSYNEQFF |
| C143 | clonotype016 | No hit | CD4+ T cells | TCR-Liao-C143-016 | 7 | CD4 | TRAV9-2*01 | TRAJ20*01 | CALRGSNDYKLSF | TRBV27*01 | TRBJ2-1*01 | CASSSLAGGKWNEQFF |
| C143 | clonotype022 | No hit | CD4+ T cells | TCR-Liao-C143-022 | 6 | CD4 | TRAV6*01 | TRAJ8*01 | CALDMSNTGFQKLVF | TRBV7-8*01 | TRBJ2-1*01 | CASTPGGQGVWGQFF |
| C143 | clonotype1068 | No hit | CD4+ T cells | TCR-Liao-C143-1068 | 3 | CD4 | TRAV8-3*01 | TRAJ4*01 | CAVGLGGYNKLIF | TRBV12-3*01 | TRBJ1-5*01 | CASGGNSNQPQHF |
| C143 | clonotype001 | No hit | CD8+ T cells | TCR-Liao-C143-001 | 24 | CD8 | TRAV12-2*01 | TRAJ20*01 | CAVRDDYKLSF | TRBV7-9*01 | TRBJ2-1*01 | CASSFGTGAREQFF |
| C143 | clonotype002 | No hit | CD8+ T cells | TCR-Liao-C143-002 | 23 | CD8 | TRAV20*01 | TRAJ9*01 | CAVQARSRLGGFKTIF | TRBV11-2*01 | TRBJ2-1*01 | CASGGPGGRPYNEQFF |
| C143 | clonotype003 | No hit | CD8+ T cells | TCR-Liao-C143-003 | 20 | CD8 | TRAV38-2/DV8*01 | TRAJ56*01 | CAYHNSTGANSKLTF | TRBV20-1*01 | TRBJ1-2*01 | CSARTTRGYTF |
| C143 | clonotype012 | No hit | CD8+ T cells | TCR-Liao-C143-012 | 8 | CD8 | TRAV13-2*01 | TRAJ17*01 | CAENSEAAGNKLTF | TRBV20-1*01 | TRBJ1-5*01 | CSARPLKGGLQPQHF |
| C143 | clonotype013 | No hit | CD8+ T cells | TCR-Liao-C143-013 | 8 | CD8 | TRAV5*01 | TRAJ5*01 | CAESDADTGRRALTF | TRBV29-1*01 | TRBJ2-1*01 | CSVEVIGYNEQFF |
| C143 | clonotype015 | No hit | CD8+ T cells | TCR-Liao-C143-015 | 7 | CD8 | TRAV3*01 | TRAJ31*01 | CAVRDVRNNNARLMF | TRBV7-2*01 | TRBJ2-5*01 | CASSLVVGGSGETQYF |
| C143 | clonotype018 | Hit | CD8+ T cells | TCR-Liao-C143-018 | 7 | CD8 | TRAV1-2*01 | TRAJ39*01 | CAVSLNNAGNMLTF | TRBV4-2*01 | TRBJ2-1*01 | CASSQETAGVNEQFF |
| C143 | clonotype020 | No hit | CD8+ T cells | TCR-Liao-C143-020 | 6 | CD8 | TRAV3*01 | TRAJ13*01 | CAVRDLSGGYQKVTF | TRBV6-5*01 | TRBJ1-2*01 | CASSSRGYGYTF |
| C143 | clonotype005 | Hit | CD4+ T cells | TCR-Liao-C143-005 | 17 | CD4 | TRAV21*01 | TRAJ37*01 | CAVRPGSGNTGKLIF | TRBV11-2*01 | TRBJ1-2*01 | CASSLGTGGRDGYTF |
| C144 | clonotype024 | No hit | CD4+ T cells | TCR-Liao-C144-024 | 1 | CD4 | TRAV12-3*01 | TRAJ41*01 | CALTLSNSGYALNF | TRBV19*01 | TRBJ2-1*01 | CATRRLGENEQFF |
| C144 | clonotype064a | No hit | CD4+ T cells | TCR-Liao-C144-064a | 1 | CD4 | TRAV1-1*01 | TRAJ32*01 | CAVDYGGATNKLIF | TRBV20-1*01 | TRBJ2-5*01 | CSARPGGETQYF |
| C144 | clonotype064b | No hit | CD4+ T cells | TCR-Liao-C144-064b | 1 | CD4 | TRAV12-1*01 | TRAJ11*01 | CAVGTGYSTLTF | TRBV20-1*01 | TRBJ2-5*01 | CSARPGGETQYF |
| C144 | clonotype001 | No hit | CD8+ T cells | TCR-Liao-C144-001 | 17 | CD8 | TRAV12-2*01 | TRAJ31*01 | CAVNNARLMF | TRBV11-2*01 | TRBJ2-5*01 | CASSLKPPTDRGRETQYF |
| C144 | clonotype002 | No hit | CD8+ T cells | TCR-Liao-C144-002 | 4 | CD8 | TRAV12-3*01 | TRAJ3*01 | CAMRQYSSASKIIF | TRBV27*01 | TRBJ2-7*01 | CASSLGSSGYEQYF |
| C144 | clonotype003 | No hit | CD8+ T cells | TCR-Liao-C144-003 | 4 | CD8 | TRAV36/DV7*01 | TRAJ47*01 | CAVPSYGNKLVF | TRBV27*01 | TRBJ1-1*01 | CASSLGQLNTEAFF |
| C144 | clonotype004 | No hit | CD8+ T cells | TCR-Liao-C144-004 | 3 | CD8 | TRAV1-2*01 | TRAJ39*01 | CAVRDRSNNAGNMLTF | TRBV4-1*01 | TRBJ1-2*01 | CASTPTDSSYGYTF |
| C144 | clonotype005a | No hit | CD8+ T cells | TCR-Liao-C144-005a | 3 | CD8 | TRAV5*01 | TRAJ13*01 | CAETSNSGGYQKVTF | TRBV28*01 | TRBJ2-5*01 | CASSSIPGDGETQYF |

| | | | | | | | | | | | | |
|---|---|---|---|---|---|---|---|---|---|---|---|---|
| C144 | clonotype005b | No hit | CD8+ T cells | TCR-Liao-C144-005b | 3 | CD8 | TRAV12-2*01 | TRAJ37*01 | CAADHSGNTGKLIF | TRBV28*01 | TRBJ2-5*01 | CASSSIPGDGETQYF |
| C144 | clonotype006 | No hit | CD8+ T cells | TCR-Liao-C144-006 | 3 | CD8 | TRAV21*01 | TRAJ11*01 | CAVGPGYSTLTF | TRBV7-9*01 | TRBJ1-1*01 | CASSSDRNTEAFF |
| C144 | clonotype007 | No hit | CD8+ T cells | TCR-Liao-C144-007 | 2 | CD8 | TRAV12-2*01 | TRAJ9*01 | CAVHTGGFKTIF | TRBV15*01 | TRBJ2-1*01 | CATSRDLRLAVYNEQFF |
| C144 | clonotype008 | No hit | CD8+ T cells | TCR-Liao-C144-008 | 2 | CD8 | TRAV3*01 | TRAJ31*01 | CAVRDYRNNNARLMF | TRBV27*01 | TRBJ2-7*01 | CASRPGQGSPYEQYF |
| C144 | clonotype009 | No hit | CD8+ T cells | TCR-Liao-C144-009 | 2 | CD8 | TRAV38-1*01 | TRAJ16*01 | CAFKFSDGQKLLF | TRBV4-1*01 | TRBJ2-2*01 | CASSHLSGGAFAGELFF |
| C144 | clonotype011 | No hit | CD8+ T cells | TCR-Liao-C144-011 | 2 | CD8 | TRAV1-2*01 | TRAJ28*01 | CAVTEQGAGSYQLTF | TRBV20-1*01 | TRBJ2-7*01 | CSARTGGGYEQYF |
| C144 | clonotype014 | No hit | CD8+ T cells | TCR-Liao-C144-014 | 2 | CD8 | TRAV8-3*01 | TRAJ21*01 | CAVGAADNFNKFYF | TRBV12-4*01 | TRBJ2-3*01 | CASRPRTTDTQYF |
| C144 | clonotype015 | No hit | CD8+ T cells | TCR-Liao-C144-015 | 2 | CD8 | TRAV12-2*01 | TRAJ49*01 | CAVSNTGNQFYF | TRBV6-5*01 | TRBJ2-3*01 | CASTRWGDTDTQYF |
| C144 | clonotype036 | No hit | CD8+ T cells | TCR-Liao-C144-036 | 1 | CD8 | TRAV12-1*01 | TRAJ10*01 | CAVNIGGAPGGGNKLTF | TRBV20-1*01 | TRBJ2-1*01 | CSARPPRRGSYNEQFF |
| C144 | clonotype069 | No hit | CD8+ T cells | TCR-Liao-C144-069 | 1 | CD8 | TRAV1-2*01 | TRAJ16*01 | CARELDGQKLLF | TRBV4-1*01 | TRBJ1-2*01 | CASSQDAGFGYTF |
| C144 | clonotype025 | No hit | CD4+ T cells | TCR-Liao-C144-025 | 1 | CD4 | TRAV3*01 | TRAJ4*01 | CAVRDLSGGYNKLIF | TRBV10-2*01 | TRBJ2-3*01 | CASSEGLGNTDTQYF |
| C146 | clonotype005 | No hit | CD4+ T cells | TCR-Liao-C146-005 | 1 | CD4 | TRAV5*01 | TRAJ37*01 | CAEHPPPGNTGKLIF | TRBV5-6*01 | TRBJ2-3*01 | CASSLASGRKDTQYF |
| C146 | clonotype013 | No hit | CD4+ T cells | TCR-Liao-C146-013 | 1 | CD4 | TRAV12-2*01 | TRAJ29*01 | CAFPNSGNTPLVF | TRBV18*01 | TRBJ2-3*01 | CASSRTSVGTDTQYF |
| C146 | clonotype031 | No hit | CD4+ T cells | TCR-Liao-C146-031 | 1 | CD4 | TRAV8-3*01 | TRAJ32*01 | CAVGAYGGATNKLIF | TRBV15*01 | TRBJ2-1*01 | CATSRDGGNNEQFF |
| C146 | clonotype040 | No hit | CD4+ T cells | TCR-Liao-C146-040 | 1 | CD4 | TRAV9-2*01 | TRAJ47*01 | CALSLDGNKLVF | TRBV20-1*01 | TRBJ2-1*01 | CSASGRRVNEQFF |
| C146 | clonotype043 | No hit | CD4+ T cells | TCR-Liao-C146-043 | 1 | CD4 | TRAV38-2/DV8*01 | TRAJ52*01 | CANAGGTSYGKLTF | TRBV2*01 | TRBJ2-7*01 | CASRGDRAIAYEQYF |
| C146 | clonotype001 | No hit | CD8+ T cells | TCR-Liao-C146-001 | 6 | CD8 | TRAV27*01 | TRAJ35*01 | CAGFMGFGNVLHC | TRBV2*01 | TRBJ2-7*01 | CASSFAPYEQYF |
| C146 | clonotype002 | No hit | CD8+ T cells | TCR-Liao-C146-002 | 4 | CD8 | TRAV8-4*01 | TRAJ27*01 | CAVTIGGGKSTF | TRBV4-1*01 | TRBJ2-5*01 | CASSGAGQETQYF |
| C146 | clonotype012 | No hit | CD8+ T cells | TCR-Liao-C146-012 | 1 | CD8 | TRAV16*01 | TRAJ8*01 | CALSGRGFQKLVF | TRBV7-8*01 | TRBJ2-7*01 | CASSLGPGINNEQYF |
| C146 | clonotype014 | No hit | CD8+ T cells | TCR-Liao-C146-014 | 1 | CD8 | TRAV29/DV5*01 | TRAJ4*01 | CAASKTGGYNKLIF | TRBV7-8*01 | TRBJ2-1*01 | CASSQGPGINNEQFF |
| C146 | clonotype016 | No hit | CD8+ T cells | TCR-Liao-C146-016 | 1 | CD8 | TRAV23/DV6*01 | TRAJ52*01 | CAASMYAGGTSYGKLTF | TRBV11-2*01 | TRBJ2-3*01 | CASSRLRGGITDTQYF |
| C146 | clonotype020b | No hit | CD8+ T cells | TCR-Liao-C146-020b | 1 | CD8 | TRAV12-2*01 | TRAJ34*01 | CAVNNGDTDKLIF | TRBV5-6*01 | TRBJ2-2*01 | CASSLGGDLAGELFF |
| C146 | clonotype021 | No hit | CD8+ T cells | TCR-Liao-C146-021 | 1 | CD8 | TRAV21*01 | TRAJ49*01 | CAVRLATGNQFYF | TRBV5-1*01 | TRBJ2-1*01 | CASSLAGGPLNEQFF |
| C146 | clonotype030 | No hit | CD8+ T cells | TCR-Liao-C146-030 | 1 | CD8 | TRAV35*01 | TRAJ29*01 | CAGRPSHSGNTPLVF | TRBV9*01 | TRBJ2-7*01 | CASSPWASGSYEQYF |
| C146 | clonotype032 | Hit | CD8+ T cells | TCR-Liao-C146-032 | 1 | CD8 | TRAV17*01 | TRAJ5*01 | CATDTGRRALTF | TRBV5-1*01 | TRBJ1-1*01 | CASSLGQGDTEAFF |
| C146 | clonotype046a | No hit | CD8+ T cells | TCR-Liao-C146-046a | 1 | CD8 | TRAV12-3*01 | TRAJ6*01 | CAKPSGGSYIPTF | TRBV19*01 | TRBJ2-1*01 | CASSISGNEQFF |
| C148 | clonotype018 | No hit | CD4+ T cells | TCR-Liao-C148-018 | 1 | CD4 | TRAV23/DV6*01 | TRAJ11*01 | CAASYLIQGTLTF | TRBV5-1*01 | TRBJ2-1*01 | CASSAPGQGYNEQFF |
| C148 | clonotype044 | No hit | CD4+ T cells | TCR-Liao-C148-044 | 1 | CD4 | TRAV8-3*01 | TRAJ56*01 | CAVGTGANSKLTF | TRBV5-1*01 | TRBJ2-5*01 | CASSLEGFGSQETQYF |

| | | | | | | | | | | | | |
|---|---|---|---|---|---|---|---|---|---|---|---|---|
| C148 | clonotype053 | No hit | CD4+ T cells | TCR-Liao-C148-053 | 1 | CD4 | TRAV29/DV5*01 | TRAJ47*01 | CAARTMEYGNKLVF | TRBV5-5*01 | TRBJ2-3*01 | CASSPPGVYTQYF |
| C148 | clonotype068 | Hit | CD4+ T cells | TCR-Liao-C148-068 | 1 | CD4 | TRAV13-2*01 | TRAJ52*01 | CAEVPTGTSYGKLTF | TRBV19*01 | TRBJ2-1*01 | CASSIVPLAGRNEQFF |
| C148 | clonotype075 | Hit | CD4+ T cells | TCR-Liao-C148-075 | 1 | CD4 | TRAV16*01 | TRAJ37*01 | CALSASSNTGKLIF | TRBV30*01 | TRBJ1-3*01 | CAWKAGGELTTGNTIYF |
| C148 | clonotype084 | No hit | CD4+ T cells | TCR-Liao-C148-084 | 1 | CD4 | TRAV13-2*01 | TRAJ17*01 | CAENKGAAGNKLTF | TRBV28*01 | TRBJ1-2*01 | CASGRRGARAGGYTF |
| C148 | clonotype100 | Hit | CD4+ T cells | TCR-Liao-C148-100 | 1 | CD4 | TRAV13-1*01 | TRAJ20*01 | CAASKDDYKLSF | TRBV24-1*01 | TRBJ2-7*01 | CATSDPRRRTYEQYF |
| C148 | clonotype107 | No hit | CD4+ T cells | TCR-Liao-C148-107 | 1 | CD4 | TRAV21*01 | TRAJ4*01 | CAAMFSGGYNKLIF | TRBV6-5*01 | TRBJ1-6*01 | CASSYSLEGPNSSPLHF |
| C148 | clonotype125 | No hit | CD4+ T cells | TCR-Liao-C148-125 | 1 | CD4 | TRAV23/DV6*01 | TRAJ34*01 | CAASSNTDKLIF | TRBV20-1*01 | TRBJ2-7*01 | CSARRQGFSYEQYF |
| C148 | clonotype001 | No hit | CD8+ T cells | TCR-Liao-C148-001 | 6 | CD8 | TRAV38-2/DV8*01 | TRAJ41*01 | CAYRSPVSALNF | TRBV4-3*01 | TRBJ2-7*01 | CASSQGTSGTYEQYF |
| C148 | clonotype003 | No hit | CD8+ T cells | TCR-Liao-C148-003 | 2 | CD8 | TRAV21*01 | TRAJ30*01 | CAVILGDDKIIF | TRBV10-3*01 | TRBJ2-2*01 | CAITEQGSGELFF |
| C148 | clonotype005 | No hit | CD8+ T cells | TCR-Liao-C148-005 | 2 | CD8 | TRAV9-2*01 | TRAJ27*01 | CPRGGSTNAGKSTF | TRBV19*01 | TRBJ2-7*01 | CASSIRGQDEQYF |
| C148 | clonotype008 | No hit | CD8+ T cells | TCR-Liao-C148-008 | 2 | CD8 | TRAV12-1*01 | TRAJ6*01 | CVVKGGSYIPTF | TRBV28*01 | TRBJ1-1*01 | CASRESGTNTEAFF |
| C148 | clonotype106 | No hit | CD4+ T cells | TCR-Liao-C148-106 | 1 | CD4 | TRAV23/DV6*01 | TRAJ54*01 | CAANIQGAQKLVF | TRBV12-4*01 | TRBJ2-7*01 | CASSHDPGRRNQVF |
| C148 | clonotype004 | No hit | CD8+ T cells | TCR-Liao-C148-004 | 2 | CD8 | TRAV14/DV4*01 | TRAJ26*01 | CAMRERGYGQNFVF | TRBV5-1*01 | TRBJ2-2*01 | CASSPETSANTGELFF |
| C148 | clonotype015 | No hit | CD8+ T cells | TCR-Liao-C148-015 | 1 | CD8 | TRAV5*01 | TRAJ31*01 | CAERDPRDARLMF | TRBV7-2*01 | TRBJ2-2*01 | CASSLDRHTGELFF |
| C148 | clonotype029 | No hit | CD8+ T cells | TCR-Liao-C148-029 | 1 | CD8 | TRAV2*01 | TRAJ33*01 | CAVEDSNYQLIW | TRBV4-1*01 | TRBJ1-1*01 | CASSLTDGFIGEAFF |
| C148 | clonotype092 | No hit | CD8+ T cells | TCR-Liao-C148-092 | 1 | CD8 | TRAV35*01 | TRAJ41*01 | CAGQEYSNSGYALNF | TRBV4-1*01 | TRBJ2-1*01 | CASSQPHGTSGSSEQFF |
| C149 | clonotype105 | No hit | CD4+ T cells | TCR-Liao-C149-105 | 1 | CD4 | TRAV3*01 | TRAJ7*01 | CAVRSVYGNNRLAF | TRBV18*01 | TRBJ2-3*01 | CASSPTDASTDTQYF |
| C149 | clonotype042 | No hit | CD4+ T cells | TCR-Liao-C149-042 | 1 | CD4 | TRAV8-1*01 | TRAJ15*01 | CAVNSNQAGTALIF | TRBV20-1*01 | TRBJ1-6*01 | CSAVPGQGSNSPLHF |
| C149 | clonotype132 | Hit | CD4+ T cells | TCR-Liao-C149-132 | 1 | CD4 | TRAV20*01 | TRAJ42*01 | CAVPAGSQGNLIF | TRBV18*01 | TRBJ2-2*01 | CASSPHPSPTGELFF |
| C149 | clonotype252 | No hit | CD4+ T cells | TCR-Liao-C149-252 | 1 | CD4 | TRAV13-1*01 | TRAJ42*01 | CAAILVYGGSQGNLIF | TRBV5-1*01 | TRBJ2-7*01 | CASSLTVQGGQYF |
| C149 | clonotype282 | No hit | CD4+ T cells | TCR-Liao-C149-282 | 1 | CD4 | TRAV17*01 | TRAJ10*01 | CATARGFTGGGNKLTF | TRBV18*01 | TRBJ2-1*01 | CASSRGTGGGNEQFF |
| C149 | clonotype003 | No hit | CD8+ T cells | TCR-Liao-C149-003 | 9 | CD8 | TRAV21*01 | TRAJ32*01 | CAVRPSGATNKLIF | TRBV7-6*01 | TRBJ1-2*01 | CASSSEPGWAYGYTF |
| C149 | clonotype004 | No hit | CD8+ T cells | TCR-Liao-C149-004 | 7 | CD8 | TRAV26-1*01 | TRAJ53*01 | CIVRAYSGGSNYKLTF | TRBV11-2*01 | TRBJ1-5*01 | CASSYHSNQPQHF |
| C149 | clonotype006 | No hit | CD8+ T cells | TCR-Liao-C149-006 | 5 | CD8 | TRAV1-2*01 | TRAJ34*01 | CAVRDRDTDKLIF | TRBV4-1*01 | TRBJ1-5*01 | CASSQDPAETRVQPQHF |
| C149 | clonotype007c | No hit | CD8+ T cells | TCR-Liao-C149-007c | 5 | CD8 | TRAV3*01 | TRAJ41*01 | CAVRWNANSGYALNF | TRBV7-6*01 | TRBJ2-7*01 | CASSPETGIHYEQYF |
| C149 | clonotype008 | Hit | CD8+ T cells | TCR-Liao-C149-008 | 4 | CD8 | TRAV30*01 | TRAJ54*01 | CGTWEDQGAQKLVF | TRBV7-2*01 | TRBJ2-1*01 | CASSLAQGPYNEQFF |
| C149 | clonotype010 | No hit | CD8+ T cells | TCR-Liao-C149-010 | 3 | CD8 | TRAV38-2/DV8*01 | TRAJ45*01 | CAYILPGGGADGLTF | TRBV7-9*01 | TRBJ2-6*01 | CASDYSGANVLTF |
| C149 | clonotype007a | No hit | CD8+ T cells | TCR-Liao-C149-007a | 5 | CD8 | TRAV12-3*01 | TRAJ18*01 | CAPNFDRGSTLGRLYF | TRBV7-6*01 | TRBJ2-7*01 | CASSPETGIHYEQYF |

| Patient | total CD4 | total CD8 | hits CD4 | hits CD8 |
|---|---|---|---|---|
| C141 | 10 | 7 | 1 | 0 |
| C142 | 5 | 9 | 1 | 0 |
| C143 | 8 | 8 | 2 | 1 |
| C144 | 4 | 15 | 0 | 0 |
| C146 | 5 | 10 | 0 | 1 |
| C148 | 10 | 8 | 3 | 0 |
| C149 | 5 | 7 | 1 | 1 |
| total | 47 | 64 | 8 | 3 |
