## Supplemental file S1 for "High resolution profiling of MHC-II peptide presentation capacity, by Mammalian Epitope Display, reveals SARS-CoV-2 targets for CD4 T cells and mechanisms of immune-escape"

**DRB10701**

| Position | Protein | Combined 20aa Peptide |
|---|---|---|
| 7 | spike glycoprotein - | LLPLVSSQCVNLTTRTQL |
| 26 | spike glycoprotein - | PAYTNSFTRGVYYPDKVFR |
| 37 | spike glycoprotein - | YYPDKVFRSSVLHSTQDLFL |
| 43 | spike glycoprotein - | FRSSVLHSTQDLFLPFFSN |
| 57 | spike glycoprotein - | PFFSNVTWFHAIHVSGTNG |
| 64 | spike glycoprotein - | WFHAIHVSGTNGTKRF |
| 88 | spike glycoprotein - | DGVYFASTEKSNIIRGW |
| 106 | spike glycoprotein - | FGTTLDSKTQSLLIVNNATN |
| 112 | spike glycoprotein - | SKTQSLLIVNNATNVVIKV |
| 200 | spike glycoprotein - | YFKIYSKHTPINLVRDLPQ |
| 228 | spike glycoprotein - | DLPIGINITRFQTLLALH |
| 239 | spike glycoprotein - | QTLLALHRSYLTPGDSS |
| 267 | spike glycoprotein - | VGYLQPRTFLLKYNENGT |
| 307 | spike glycoprotein - | TVEKGIYQTSNFRVQPTES |
| 314 | spike glycoprotein - | QTSNFRVQPTESIVRFPNIT |
| 320 | spike glycoprotein - | VQPTESIVRFPNITN |
| 391 | spike glycoprotein - | CFTNVYADSFVIRGDEVRQ |
| 433 | spike glycoprotein - | VIAWNSNNLDSKVGG |
| 454 | spike glycoprotein - | RLFRKSNLKPFERDIS |
| 461 | spike glycoprotein - | LKPFERDISTEIYQAG |
| 470 | spike glycoprotein - | TEIYQAGSTPCNGVEGF |
| 489 | spike glycoprotein - | YFPLQSYGFQPTNGV |
| 495 | spike glycoprotein - | YGFQPTNGVGYQPYR |
| 530 | spike glycoprotein - | STNLVKNKCVNFNFNGLT |
| 541 | spike glycoprotein - | FNFNGLTGTGVLTESNK |
| 632 | spike glycoprotein - | TWRVYSTGSNVFQTRAG |
| 677 | spike glycoprotein - | QTNSPRRARSVASQSIIAYT |
| 683 | spike glycoprotein - | RARSVASQSIIAYTMSLGAE |
| 689 | spike glycoprotein - | SQSIIAYTMSLGAENSVAYS |
| 695 | spike glycoprotein - | YTMSLGAENSVAYSNNS |
| 700 | spike glycoprotein - | GAENSVAYSNNSIAIPTNFT |
| 715 | spike glycoprotein - | PTNFTISVTTEILPVS |
| 722 | spike glycoprotein - | VTTEILPVSMTKTSVD |
| 781 | spike glycoprotein - | VFAQVKQIYKTPPIKDFGG |
| 822 | spike glycoprotein - | LFNKVTLADAGFIKQYG |
| 851 | spike glycoprotein - | CAQKFNGLTVLPPLLTDE |
| 869 | spike glycoprotein - | MIAQYTSALLAGTITSGWTF |
| 880 | spike glycoprotein - | GTITSGWTFGAGAAL |
| 895 | spike glycoprotein - | QIPFAMQMAYRFNGIGV |
| 905 | spike glycoprotein - | RFNGIGVTQNVLYENQKL |
| 921 | spike glycoprotein - | KLIANQFNSAIGKIQDSLSS |

| | | |
|---|---|---|
| 933 | spike glycoprotein - | KIQDSLSSTASALGKLQDVV |
| 1007 | spike glycoprotein - | YVTQQLIRAAEIRASANLAA |
| 1013 | spike glycoprotein - | IRAAEIRASANLAATKMSEC |
| 1019 | spike glycoprotein - | RASANLAATKMSECVLGQ |
| 1025 | spike glycoprotein - | AATKMSECVLGQSKR |
| 1046 | spike glycoprotein - | GYHLMSFPQSAPHGV |
| 1066 | spike glycoprotein - | TYVPAQEKNFTTAPAI |
| 1072 | spike glycoprotein - | EKNFTTAPAICHDGKA |
| 1131 | spike glycoprotein - | GIVNNTVYDPLQPELD |
| 1153 | spike glycoprotein - | DKYFKNHTSPDVDLGDISG |
| 9 | ORF8 - | IITTVAAFHQECSLQSC |
| 18 | ORF8 - | QECSLQSCTQHQPYVVD |
| 24 | ORF8 - | SCTQHQPYVVDDPCPIH |
| 44 | ORF8 - | KWYIRVGARKSAPLIELCVD |
| 82 | ORF8 - | SCLPFTINCQEPKLGSL |
| 8 | ORF7b - | DFYLCFLAFLLFLVLIML |
| 28 | ORF7b - | FWFSLELQDHNETCHA |
| 1 | ORF7a - | MKIILFLALITLATCEL |
| 33 | ORF7a - | EPCSSGTYEGNSPFHPL |
| 59 | ORF7a - | FSTQFAFACPDGVKH |
| 69 | ORF7a - | DGVKHVYQLRARSVSPKLFI |
| 75 | ORF7a - | YQLRARSVSPKLFIRQ |
| 80 | ORF7a - | RSVSPKLFIRQEEVQ |
| 35 | ORF6 - | LIIKNLSKSLTENKYSQLDE |
| 41 | ORF6 - | SKSLTENKYSQLDEEQP |
| 47 | ORF6 - | NKYSQLDEEQPMEID |
| 5 | ORF3a - | MRIFTIGTVTLKQGE |
| 10 | ORF3a - | IGTVTLKQGEIKDATPSDF |
| 23 | ORF3a - | ATPSDFVRATATIPIQASLP |
| 29 | ORF3a - | VRATATIPIQASLPFGWL |
| 54 | ORF3a - | AVFQSASKIITLKKRWQLAL |
| 62 | ORF3a - | IITLKKRWQLALSKGVHFVC |
| 68 | ORF3a - | RWQLALSKGVHFVCN |
| 166 | ORF3a - | SIVITSGDGTTSPISEHD |
| 209 | ORF3a - | SDYYQLYSTQLSTDTG |
| 233 | ORF3a - | YNKIVDEPEEHVQIH |
| 251 | ORF3a - | GSSGVVNPVMEPIYDEPTT |
| 4462 | ORF1b polyprotein - | FVVKRHTFSNYQHEE |
| 4496 | ORF1b polyprotein - | FRIDGDMVPHISRQR |
| 4499 | ORF1b polyprotein - | DGDMVPHISRQRLTKYTMAD |
| 4505 | ORF1b polyprotein - | HISRQRLTKYTMADL |
| 4561 | ORF1b polyprotein - | PDILRVYANLGERVRQ |
| 4611 | ORF1b polyprotein - | FGDFIQTTPGSGVPVVDSY |

| | | |
|---|---|---|
| 4633 | ORF1b polyprotein - | LMPILTLTRALTAESHVDTD |
| 4639 | ORF1b polyprotein - | LTRALTAESHVDTDL |
| 4679 | ORF1b polyprotein - | FKYWDQTYHPNCVNCL |
| 4727 | ORF1b polyprotein - | VDGVPFVVSTGYHFRELGVV |
| 4733 | ORF1b polyprotein - | VVSTGYHFRELGVVHNQDV |
| 4747 | ORF1b polyprotein - | HNQDVNLHSSRLSFKELL |
| 4767 | ORF1b polyprotein - | AADPAMHAASGNLLLDKRTT |
| 4773 | ORF1b polyprotein - | HAASGNLLLDKRTTC |
| 4791 | ORF1b polyprotein - | AALTNNVAFQTVKPGNFNKD |
| 4797 | ORF1b polyprotein - | VAFQTVKPGNFNKDFY |
| 4801 | ORF1b polyprotein - | TVKPGNFNKDFYDFA |
| 4811 | ORF1b polyprotein - | FYDFAVSKGFFKEGSSVELK |
| 4817 | ORF1b polyprotein - | SKGFFKEGSSVELKHFF |
| 4887 | ORF1b polyprotein - | VNNLDKSAGFPFNKWGK |
| 4913 | ORF1b polyprotein - | YEDQDALFAYTKRNVIPTIT |
| 4919 | ORF1b polyprotein - | LFAYTKRNVIPTITQMN |
| 4934 | ORF1b polyprotein - | MNLKYAISAKNRARTVAG |
| 4944 | ORF1b polyprotein - | NRARTVAGVSICSTM |
| 4952 | ORF1b polyprotein - | VSICSTMTNRQFHQKLL |
| 4966 | ORF1b polyprotein - | KLLKSIAATRGATVVIGTSK |
| 4972 | ORF1b polyprotein - | AATRGATVVIGTSKFY |
| 5023 | ORF1b polyprotein - | RIMASLVLARKHTTC |
| 5075 | ORF1b polyprotein - | GDATTAYANSVFNICQAVT |
| 5085 | ORF1b polyprotein - | VFNICQAVTANVNALLSTDG |
| 5091 | ORF1b polyprotein - | AVTANVNALLSTDGNKIADK |
| 5097 | ORF1b polyprotein - | NALLSTDGNKIADKYVRN |
| 5137 | ORF1b polyprotein - | FYAYLRKHFSMMILSDDAVV |
| 5155 | ORF1b polyprotein - | VVCFNSTYASQGLVASIKNF |
| 5167 | ORF1b polyprotein - | LVASIKNFKSVLYYQNNVF |
| 5202 | ORF1b polyprotein - | HEFCSQHTMLVKQGD |
| 5237 | ORF1b polyprotein - | DDIVKTDGTLMIERFV |
| 5246 | ORF1b polyprotein - | LMIERFVSLAIDAYPLTKH |
| 5254 | ORF1b polyprotein - | LAIDAYPLTKHPNQEYADVF |
| 5288 | ORF1b polyprotein - | TGHMLDMYSVMLTNDNTSR |
| 5353 | ORF1b polyprotein - | CCYDHVISTSHKLVLSVNPY |
| 5359 | ORF1b polyprotein - | ISTSHKLVLSVNPYVC |
| 5437 | ORF1b polyprotein - | DWTNAGDYILANTCT |
| 5452 | ORF1b polyprotein - | ERLKLFAAETLKATEETF |
| 5465 | ORF1b polyprotein - | TEETFKLSYGIATVREVLSD |
| 5471 | ORF1b polyprotein - | LSYGIATVREVLSDRE |
| 5505 | ORF1b polyprotein - | VFTGYRVTKNSKVQIGEY |
| 5519 | ORF1b polyprotein - | IGEYTFEKGDYGDAV |
| 5523 | ORF1b polyprotein - | TFEKGDYGDAVVYRG |

| | | |
|---|---|---|
| 5530 | ORF1b polyprotein - | GDAVVYRGTTTYKLNVGDY |
| 5544 | ORF1b polyprotein - | NVGDYFVLTSHTVMPLSAPT |
| 5550 | ORF1b polyprotein - | VLTSHTVMPLSAPTLVPQE |
| 5568 | ORF1b polyprotein - | EHYVRITGLYPTLNISDEFS |
| 5576 | ORF1b polyprotein - | LYPTLNISDEFSSNV |
| 5583 | ORF1b polyprotein - | SDEFSSNVANYQKVGM |
| 5593 | ORF1b polyprotein - | YQKVGMQKYSTLQGPPG |
| 5609 | ORF1b polyprotein - | GTGKSHFAIGLALYYPS |
| 5619 | ORF1b polyprotein - | LALYYPSARIVYTACS |
| 5657 | ORF1b polyprotein - | IIPARARVECFDKFK |
| 5696 | ORF1b polyprotein - | VFDEISMATNYDLSVVNARL |
| 5702 | ORF1b polyprotein - | MATNYDLSVVNARLR |
| 5715 | ORF1b polyprotein - | LRAKHYVYIGDPAQLPAPR |
| 5727 | ORF1b polyprotein - | AQLPAPRTLLTKGTL |
| 5735 | ORF1b polyprotein - | LLTKGTLEPEYFNSV |
| 5770 | ORF1b polyprotein - | AEIVDTVSALVYDNKLKA |
| 5795 | ORF1b polyprotein - | CFKMFYKGVITHDVSSAIN |
| 5843 | ORF1b polyprotein - | NAVASKILGLPTQTV |
| 5846 | ORF1b polyprotein - | ASKILGLPTQTVDSSQG |
| 5906 | ORF1b polyprotein - | YDKLQFTSLEIPRRNVATL |
| 5946 | ORF1b polyprotein - | TQAPTHLSVDTKFKT |
| 5964 | ORF1b polyprotein - | CVDIPGIPKDMTYRR |
| 5978 | ORF1b polyprotein - | RLISMMGFKMNYQVN |
| 6003 | ORF1b polyprotein - | EAIRHVRAWIGFDVEGCHA |
| 6029 | ORF1b polyprotein - | NLPLQLGFSTGVNLVAV |
| 6040 | ORF1b polyprotein - | VNLVAVPTGYVDTPNNT |
| 6058 | ORF1b polyprotein - | FSRVSAKPPPGDQFKH |
| 6111 | ORF1b polyprotein - | WAHGFELTSMKYFVKI |
| 6155 | ORF1b polyprotein - | SIGFDYVVYNPFMIDVQ |
| 6168 | ORF1b polyprotein - | DVQQWGFTGNLQSNHDLY |
| 6185 | ORF1b polyprotein - | YCQVHGNAHVASCDAIM |
| 6213 | ORF1b polyprotein - | KRVDWTIEYPIIGDEL |
| 6221 | ORF1b polyprotein - | YPIIGDELKINAACR |
| 6235 | ORF1b polyprotein - | RKVQHMVVKAALLADKFPVL |
| 6341 | ORF1b polyprotein - | GGSLYVNKHAFHTPA |
| 6354 | ORF1b polyprotein - | PAFDKSAFVNLKQLPFFY |
| 6418 | ORF1b polyprotein - | LYLDAYNMMISAGFSL |
| 6424 | ORF1b polyprotein - | NMMISAGFSLWVYKQFD |
| 6446 | ORF1b polyprotein - | NTFTRLQSLENVAFNVVNK |
| 6462 | ORF1b polyprotein - | VNKGHFDGQQGEVPVSII |
| 6478 | ORF1b polyprotein - | IINNTVYTKVDGVDVELFE |
| 6510 | ORF1b polyprotein - | WAKRNIKPVPEVKILNNLGV |
| 6516 | ORF1b polyprotein - | KPVPEVKILNNLGVDIAANT |

| | | |
|---|---|---|
| 6522 | ORF1b polyprotein - | KILNNLGVDIAANTVI |
| 6583 | ORF1b polyprotein - | VDLFRNARNGVLITEGSV |
| 6592 | ORF1b polyprotein - | GVLITEGSVKGLQPS |
| 6601 | ORF1b polyprotein - | KGLQPSVGPKQASLNGVT |
| 6619 | ORF1b polyprotein - | LIGEAVKTQFNYYKK |
| 6627 | ORF1b polyprotein - | QFNYYKKVDGVVQQLPETY |
| 6639 | ORF1b polyprotein - | QQLPETYFTQSRNLQEFKPR |
| 6645 | ORF1b polyprotein - | YFTQSRNLQEFKPRSQ |
| 6686 | ORF1b polyprotein - | HIVYGDFSHSQLGGLHLLIG |
| 6752 | ORF1b polyprotein - | DDFVEIIKSQDLSVVSKVV |
| 6795 | ORF1b polyprotein - | PKLQSSQAWQPGVAMPNL |
| 6848 | ORF1b polyprotein - | LCQYLNTLTLAVPYNMRVI |
| 6862 | ORF1b polyprotein - | NMRVIHFGAGSDKGVAP |
| 6880 | ORF1b polyprotein - | TAVLRQWLPTGTLLVDSDLN |
| 6886 | ORF1b polyprotein - | WLPTGTLLVDSDLNDFVS |
| 6924 | ORF1b polyprotein - | LIISDMYDPKTKNVTKEND |
| 6955 | ORF1b polyprotein - | IQQKLALGGSVAIKITEHSW |
| 6961 | ORF1b polyprotein - | LGGSVAIKITEHSWNADLYK |
| 6967 | ORF1b polyprotein - | IKITEHSWNADLYKL |
| 6985 | ORF1b polyprotein - | FAWWTAFVTNVNASS |
| 6991 | ORF1b polyprotein - | FVTNVNASSSEAFLIGCN |
| 7018 | ORF1b polyprotein - | DGYVMHANYIFWRNTNPIQL |
| 7024 | ORF1b polyprotein - | ANYIFWRNTNPIQLSSYS |
| 7045 | ORF1b polyprotein - | MSKFPLKLRGTAVMSLKEGQ |
| 7051 | ORF1b polyprotein - | KLRGTAVMSLKEGQI |
| 7069 | ORF1b polyprotein - | ILSLLSKGRLIIRENNRVV |
| 7080 | ORF1b polyprotein - | IRENNRVVISSDVLVNN |
| 16 | ORF1a polyprotein - | LSLPVLQVRDVLVRG |
| 68 | ORF1a polyprotein - | YVFIKRSDARTAPHGHVM |
| 104 | ORF1a polyprotein - | LGVLVPHVGEIPVAYRKV |
| 114 | ORF1a polyprotein - | IPVAYRKVLLRKNGNKGA |
| 121 | ORF1a polyprotein - | VLLRKNGNKGAGGHS |
| 124 | ORF1a polyprotein - | RKNGNKGAGGHSYGAD |
| 159 | ORF1a polyprotein - | ENWNTKHSSGVTREL |
| 204 | ORF1a polyprotein - | LLARAGKASCTLSEQL |
| 246 | ORF1a polyprotein - | EKSYELQTPFEIKLAKKFDT |
| 252 | ORF1a polyprotein - | QTPFEIKLAKKFDTF |
| 255 | ORF1a polyprotein - | FEIKLAKKFDTFNGE |
| 270 | ORF1a polyprotein - | CPNFVFPLNSIIKTIQPRVE |
| 276 | ORF1a polyprotein - | PLNSIIKTIQPRVEKKKLD |
| 295 | ORF1a polyprotein - | GFMGRIRSVYPVASPNECNQ |
| 301 | ORF1a polyprotein - | RSVYPVASPNECNQM |
| 337 | ORF1a polyprotein - | VKATCEFCGTENLTKEG |

| | | |
|---|---|---|
| 390 | ORF1a polyprotein - | ESGLKTILRKGGRTI |
| 415 | ORF1a polyprotein - | GCHNKCAYWVPRASA |
| 459 | ORF1a polyprotein - | VNINIVGDFKLNEEIAII |
| 475 | ORF1a polyprotein - | IILASFSASTSAFVETVKGL |
| 481 | ORF1a polyprotein - | SASTSAFVETVKGLD |
| 484 | ORF1a polyprotein - | TSAFVETVKGLDYKA |
| 501 | ORF1a polyprotein - | QIVESCGNFKVTKGK |
| 504 | ORF1a polyprotein - | ESCGNFKVTKGKAKK |
| 514 | ORF1a polyprotein - | GKAKKGAWNIGEQKSILSPL |
| 539 | ORF1a polyprotein - | EAARVVRSIFSRTLETAQN |
| 558 | ORF1a polyprotein - | SVRVLQKAAITILDGISQYS |
| 564 | ORF1a polyprotein - | KAAITILDGISQYSLRLID |
| 576 | ORF1a polyprotein - | YSLRLIDAMMFTSDLATNNL |
| 584 | ORF1a polyprotein - | MMFTSDLATNNLVVM |
| 597 | ORF1a polyprotein - | VMAYITGGVVQLTSQ |
| 604 | ORF1a polyprotein - | GVVQLTSQWLTNIFG |
| 620 | ORF1a polyprotein - | VYEKLKPVLDWLEEKF |
| 756 | ORF1a polyprotein - | VVLKTGDLQPLEQPT |
| 759 | ORF1a polyprotein - | KTGDLQPLEQPTSEA |
| 796 | ORF1a polyprotein - | TEKYCALAPNMMVTNNTFTL |
| 802 | ORF1a polyprotein - | LAPNMMVTNNTFTLKGGAPT |
| 815 | ORF1a polyprotein - | LKGGAPTKVTFGDDTVI |
| 828 | ORF1a polyprotein - | DTVIEVQGYKSVNITFELDE |
| 834 | ORF1a polyprotein - | QGYKSVNITFELDER |
| 846 | ORF1a polyprotein - | DERIDKVLNEKCSAY |
| 871 | ORF1a polyprotein - | FACVVADAVIKTLQPVS |
| 957 | ORF1a polyprotein - | GKPLEFGATSAALQPEEEQE |
| 963 | ORF1a polyprotein - | GATSAALQPEEEQEED |
| 1010 | ORF1a polyprotein - | PQLEMELTPVVQTIEVNS |
| 1030 | ORF1a polyprotein - | GYLKLTDNVYIKNADIVEEA |
| 1036 | ORF1a polyprotein - | DNVYIKNADIVEEAKKVKPT |
| 1050 | ORF1a polyprotein - | KKVKPTVVVNAANVYLKHGG |
| 1056 | ORF1a polyprotein - | VVVNAANVYLKHGGGVAG |
| 1065 | ORF1a polyprotein - | LKHGGGVAGALNKAT |
| 1071 | ORF1a polyprotein - | VAGALNKATNNAMQVESDDY |
| 1077 | ORF1a polyprotein - | KATNNAMQVESDDYI |
| 1089 | ORF1a polyprotein - | DYIATNGPLKVGGSCV |
| 1132 | ORF1a polyprotein - | KSAYENFNQHEVLLAPLLSA |
| 1144 | ORF1a polyprotein - | LLAPLLSAGIFGADPIHS |
| 1169 | ORF1a polyprotein - | VRTNVYLAVFDKNLYDKLV |
| 1188 | ORF1a polyprotein - | SSFLEMKSEKQVEQKI |
| 1207 | ORF1a polyprotein - | PKEEVKPFITESKPSVEQRK |
| 1213 | ORF1a polyprotein - | PFITESKPSVEQRKQD |

| | | |
|---|---|---|
| 1262 | ORF1a polyprotein - | NLHPDSATLVSDIDITFL |
| 1275 | ORF1a polyprotein - | DITFLKKDAPYIVGDVVQEG |
| 1281 | ORF1a polyprotein - | KDAPYIVGDVVQEGV |
| 1300 | ORF1a polyprotein - | VIPTKKAGGTTEMLA |
| 1314 | ORF1a polyprotein - | AKALRKVPTDNYITTYPGQG |
| 1337 | ORF1a polyprotein - | YTVEEAKTVLKKCKSAF |
| 1350 | ORF1a polyprotein - | KSAFYILPSIISNEKQEILG |
| 1356 | ORF1a polyprotein - | LPSIISNEKQEILGTV |
| 1367 | ORF1a polyprotein - | ILGTVSWNLREMLAHA |
| 1391 | ORF1a polyprotein - | VCVETKAIVSTIQRKYK |
| 1396 | ORF1a polyprotein - | KAIVSTIQRKYKGIK |
| 1399 | ORF1a polyprotein - | VSTIQRKYKGIKIQEGVVDY |
| 1405 | ORF1a polyprotein - | KYKGIKIQEGVVDYG |
| 1422 | ORF1a polyprotein - | FYFYTSKTTVASLINTLNDL |
| 1428 | ORF1a polyprotein - | KTTVASLINTLNDLNE |
| 1448 | ORF1a polyprotein - | MPLGYVTHGLNLEEAA |
| 1458 | ORF1a polyprotein - | NLEEAARYMRSLKVPATVSV |
| 1464 | ORF1a polyprotein - | RYMRSLKVPATVSVSSPDA |
| 1486 | ORF1a polyprotein - | YNGYLTSSSKTPEEHFIE |
| 1494 | ORF1a polyprotein - | SKTPEEHFIETISLAGSYKD |
| 1500 | ORF1a polyprotein - | HFIETISLAGSYKDWSYSGQ |
| 1506 | ORF1a polyprotein - | SLAGSYKDWSYSGQS |
| 1509 | ORF1a polyprotein - | GSYKDWSYSGQSTQLGIE |
| 1530 | ORF1a polyprotein - | RGDKSVYYTSNPTTFHLDGE |
| 1536 | ORF1a polyprotein - | YYTSNPTTFHLDGEVI |
| 1560 | ORF1a polyprotein - | LSLREVRTIKVFTTVDNINL |
| 1566 | ORF1a polyprotein - | RTIKVFTTVDNINLHTQVVD |
| 1572 | ORF1a polyprotein - | TTVDNINLHTQVVDM |
| 1582 | ORF1a polyprotein - | QVVDMSMTYGQQFGPTYL |
| 1632 | ORF1a polyprotein - | FEYYHTTDPSFLGRYMSA |
| 1724 | ORF1a polyprotein - | ELGDVRETMSYLFQHANLDS |
| 1746 | ORF1a polyprotein - | RVLNVVCKTCGQQQTT |
| 1758 | ORF1a polyprotein - | QQTTLKGVEAVMYMGTL |
| 1764 | ORF1a polyprotein - | GVEAVMYMGTLSYEQFKKG |
| 1781 | ORF1a polyprotein - | KGVQIPCTCGKQATKY |
| 1794 | ORF1a polyprotein - | TKYLVQQESPFVMMS |
| 1797 | ORF1a polyprotein - | LVQQESPFVMMSAPP |
| 1801 | ORF1a polyprotein - | ESPFVMMSAPPAQYEL |
| 1848 | ORF1a polyprotein - | IDGALLTKSSEYKGPITDV |
| 1859 | ORF1a polyprotein - | YKGPITDVFYKENSY |
| 1911 | ORF1a polyprotein - | PIDLVPNQPYPNASF |
| 1944 | ORF1a polyprotein - | LTGYKKPASRELKVT |
| 1982 | ORF1a polyprotein - | GAKLLHKPIVWHVNNATNKA |

| | | |
|---|---|---|
| 1988 | ORF1a polyprotein - | KPIVWHVNNATNKATY |
| 1996 | ORF1a polyprotein - | NATNKATYKPNTWCIR |
| 2012 | ORF1a polyprotein - | CLWSTKPVETSNSFDVL |
| 2017 | ORF1a polyprotein - | KPVETSNSFDVLKSEDA |
| 2073 | ORF1a polyprotein - | GDIILKPANNSLKITEEVG |
| 2082 | ORF1a polyprotein - | NSLKITEEVGHTDLMAAY |
| 2108 | ORF1a polyprotein - | KKPNELSRVLGLKTLATHGL |
| 2114 | ORF1a polyprotein - | SRVLGLKTLATHGLA |
| 2123 | ORF1a polyprotein - | ATHGLAAVNSVPWDTIA |
| 2141 | ORF1a polyprotein - | YAKPFLNKVVSTTTNIVTR |
| 2180 | ORF1a polyprotein - | CTFTRSTNSRIKASM |
| 2187 | ORF1a polyprotein - | NSRIKASMPTTIAKNTVKS |
| 2196 | ORF1a polyprotein - | TTIAKNTVKSVGKFC |
| 2213 | ORF1a polyprotein - | ASFNYLKSPNFSKLIN |
| 2243 | ORF1a polyprotein - | LIYSTAALGVLMSNL |
| 2366 | ORF1a polyprotein - | WLIINLVQMAPISAMVRM |
| 2471 | ORF1a polyprotein - | RDLSLQFKRPINPTDQ |
| 2480 | ORF1a polyprotein - | PINPTDQSSYIVDSV |
| 2509 | ORF1a polyprotein - | GQKTYERHSLSHFVNLDNLR |
| 2537 | ORF1a polyprotein - | PINVIVFDGKSKCEE |
| 2569 | ORF1a polyprotein - | ILLLDQALVSDVGDSA |
| 2575 | ORF1a polyprotein - | ALVSDVGDSAEVAVKM |
| 2591 | ORF1a polyprotein - | FDAYVNTFSSTFNVPMEKLK |
| 2597 | ORF1a polyprotein - | TFSSTFNVPMEKLKT |
| 2608 | ORF1a polyprotein - | KLKTLVATAEAELAKNVSLD |
| 2632 | ORF1a polyprotein - | TFISAARQGFVDSDVETK |
| 2653 | ORF1a polyprotein - | ECLKLSHQSDIEVTGDSCNN |
| 2659 | ORF1a polyprotein - | HQSDIEVTGDSCNNY |
| 2671 | ORF1a polyprotein - | NNYMLTYNKVENMTPRDLG |
| 2695 | ORF1a polyprotein - | SARHINAQVAKSHNIALIWN |
| 2701 | ORF1a polyprotein - | AQVAKSHNIALIWNV |
| 2746 | ORF1a polyprotein - | TTRQVVNVVTTKIALKGGKI |
| 2752 | ORF1a polyprotein - | NVVTTKIALKGGKIVN |
| 2756 | ORF1a polyprotein - | TKIALKGGKIVNNWL |
| 2794 | ORF1a polyprotein - | HVMSKHTDFSSEIIGYKAI |
| 2806 | ORF1a polyprotein - | IIGYKAIDGGVTRDI |
| 2852 | ORF1a polyprotein - | PLIAAVITREVGFVVPGLPG |
| 2858 | ORF1a polyprotein - | ITREVGFVVPGLPGT |
| 2866 | ORF1a polyprotein - | VPGLPGTILRTTNGDF |
| 2886 | ORF1a polyprotein - | PRVFSAVGNICYTPSKLIEY |
| 2892 | ORF1a polyprotein - | VGNICYTPSKLIEYTDFATS |
| 2898 | ORF1a polyprotein - | TPSKLIEYTDFATSA |
| 2957 | ORF1a polyprotein - | MDGSIIQFPNTYLEGSVRV |

| | | |
|---|---|---|
| 2967 | ORF1a polyprotein - | TYLEGSVRVVTTFDS |
| 3029 | ORF1a polyprotein - | NMFTPLIQPIGALDISASIV |
| 3035 | ORF1a polyprotein - | IQPIGALDISASIVAGGIVA |
| 3041 | ORF1a polyprotein - | LDISASIVAGGIVAIV |
| 3065 | ORF1a polyprotein - | MRFRRAFGEYSHVVAFNTL |
| 3121 | ORF1a polyprotein - | SFLAHIQWMVMFTPLVPF |
| 3154 | ORF1a polyprotein - | YWFFSNYLKRRVVFN |
| 3185 | ORF1a polyprotein - | LNKEMYLKLRSDVLL |
| 3188 | ORF1a polyprotein - | EMYLKLRSDVLLPLTQYNR |
| 3209 | ORF1a polyprotein - | ALYNKYKYFSGAMDTTSY |
| 3239 | ORF1a polyprotein - | NDFSNSGSDVLYQPPQTSI |
| 3246 | ORF1a polyprotein - | SDVLYQPPQTSITSA |
| 3250 | ORF1a polyprotein - | YQPPQTSITSAVLQSGFRKM |
| 3256 | ORF1a polyprotein - | SITSAVLQSGFRKMA |
| 3265 | ORF1a polyprotein - | GFRKMAFPSGKVEGC |
| 3283 | ORF1a polyprotein - | VTCGTTTLNGLWLDD |
| 3324 | ORF1a polyprotein - | KSNHNFLVQAGNVQLRVIGH |
| 3330 | ORF1a polyprotein - | LVQAGNVQLRVIGHS |
| 3362 | ORF1a polyprotein - | PKYKFVRIQPGQTFSVL |
| 3391 | ORF1a polyprotein - | CAMRPNFTIKGSFLN |
| 3469 | ORF1a polyprotein - | AWLYAAVINGDRWFLN |
| 3484 | ORF1a polyprotein - | NRFTTTLNDFNLVAMK |
| 3511 | ORF1a polyprotein - | DILGPLSAQTGIAVLDMC |
| 3570 | ORF1a polyprotein - | SAVKRTIKGTHHWLL |
| 3631 | ORF1a polyprotein - | HKHAFLCLFLLPSLATVA |
| 3691 | ORF1a polyprotein - | LLILMTARTVYDDGARR |
| 3735 | ORF1a polyprotein - | ALIISVTSNYSGVVT |
| 3738 | ORF1a polyprotein - | ISVTSNYSGVVTTVMF |
| 3794 | ORF1a polyprotein - | FGLFCLLNRYFRLTL |
| 3812 | ORF1a polyprotein - | DYLVSTQEFRYMNSQGLLPP |
| 3818 | ORF1a polyprotein - | QEFRYMNSQGLLPPKNSID |
| 3841 | ORF1a polyprotein - | NIKLLGVGGKPCIKVATV |
| 3862 | ORF1a polyprotein - | MSDVKCTSVVLLSVLQ |
| 3891 | ORF1a polyprotein - | CVQLHNDILLAKDTT |
| 3941 | ORF1a polyprotein - | LQAIASEFSSLPSYA |
| 3945 | ORF1a polyprotein - | ASEFSSLPSYAAFATAQEAY |
| 3976 | ORF1a polyprotein - | VLKKLKKSLNVAKSEFDRDA |
| 3982 | ORF1a polyprotein - | KSLNVAKSEFDRDAAMQRKL |
| 3997 | ORF1a polyprotein - | MQRKLEKMADQAMTQM |
| 4003 | ORF1a polyprotein - | KMADQAMTQMYKQARSED |
| 4009 | ORF1a polyprotein - | MTQMYKQARSEDKRAKVTS |
| 4028 | ORF1a polyprotein - | AMQTMLFTMLRKLDNDA |
| 4055 | ORF1a polyprotein - | GCVPLNIIPLTTAAKLMVVI |

| | | |
|---|---|---|
| 4061 | ORF1a polyprotein - | IIPLTTAAKLMVVIPDYNTY |
| 4069 | ORF1a polyprotein - | KLMVVIPDYNTYKNT |
| 4085 | ORF1a polyprotein - | DGTTFTYASALWEIQQVV |
| 4100 | ORF1a polyprotein - | QVVDADSKIVQLSEI |
| 4123 | ORF1a polyprotein - | AWPLIVTALRANSAVKLQNN |
| 4129 | ORF1a polyprotein - | TALRANSAVKLQNNEL |
| 4168 | ORF1a polyprotein - | ALAYYNTTKGGRFVLALLS |
| 4228 | ORF1a polyprotein - | LYFIKGLNNLNRGMVLGSL |
| 4239 | ORF1a polyprotein - | RGMVLGSLAATVRLQAGNA |
| 4247 | ORF1a polyprotein - | AATVRLQAGNATEVPANSTV |
| 4278 | ORF1a polyprotein - | KAYKDYLASGGQPITNCVKM |
| 4284 | ORF1a polyprotein - | LASGGQPITNCVKMLC |
| 4304 | ORF1a polyprotein - | TGQAITVTPEANMDQES |
| 4310 | ORF1a polyprotein - | VTPEANMDQESFGGASCC |
| 1 | ORF10 | MGYINVFAFPFTIYSLL |
| 6 | ORF10 | VFAFPFTIYSLLLCR |
| 20 | ORF10 | RMNSRNYIAQVDVVN |
| 1 | nucleocapsid phosphoprotein - | MSDNGPQNQRNAPRI |
| 16 | nucleocapsid phosphoprotein - | TFGGPSDSTGSNQNGERS |
| 61 | nucleocapsid phosphoprotein - | KEDLKFPRGQGVPINTNSS |
| 130 | nucleocapsid phosphoprotein - | IIWVATEGALNTPKDHI |
| 138 | nucleocapsid phosphoprotein - | ALNTPKDHIGTRNPA |
| 154 | nucleocapsid phosphoprotein - | NAAIVLQLPQGTTLP |
| 157 | nucleocapsid phosphoprotein - | IVLQLPQGTTLPKGF |
| 184 | nucleocapsid phosphoprotein - | SRSSSRSRNSSRNSTP |
| 188 | nucleocapsid phosphoprotein - | SRSRNSSRNSTPGSS |
| 202 | nucleocapsid phosphoprotein - | SRGTSPARMAGNGGDAA |
| 249 | nucleocapsid phosphoprotein - | KSAAEASKKPRQKRTA |
| 260 | nucleocapsid phosphoprotein - | QKRTATKAYNVTQAFGRRGP |
| 266 | nucleocapsid phosphoprotein - | KAYNVTQAFGRRGPEQ |
| 302 | nucleocapsid phosphoprotein - | PQIAQFAPSASAFFGMSRIG |
| 308 | nucleocapsid phosphoprotein - | APSASAFFGMSRIGM |
| 321 | nucleocapsid phosphoprotein - | GMEVTPSGTWLTYTGAIKLD |
| 327 | nucleocapsid phosphoprotein - | SGTWLTYTGAIKLDDKDPNF |
| 333 | nucleocapsid phosphoprotein - | YTGAIKLDDKDPNFKDQ |
| 355 | nucleocapsid phosphoprotein - | KHIDAYKTFPPTEPK |
| 374 | nucleocapsid phosphoprotein - | KKADETQALPQRQKKQQTVT |
| 99 | membrane glycoprotein - | SFRLFARTRSMWSFNPET |
| 110 | membrane glycoprotein - | WSFNPETNILLNVPLHG |
| 122 | membrane glycoprotein - | VPLHGTILTRPLLESEL |
| 134 | membrane glycoprotein - | LESELVIGAVILRGH |
| 163 | membrane glycoprotein - | DLPKEITVATSRTLSYYKLG |
| 169 | membrane glycoprotein - | TVATSRTLSYYKLGASQRV |

| | | |
|---|---|---|
| 176 | membrane glycoprotein - | LSYYKLGASQRVAGDSGFAA |
| 201 | membrane glycoprotein - | IGNYKLNTDHSSSSDNIA |
| 45 | envelope protein - | NIVNVSLVKPSFYVYS |
| 54 | envelope protein - | PSFYVYSRVKNLNSSRVPDL |
| 60 | envelope protein - | SRVKNLNSSRVPDLLV |

**DRB10404**

| Position | Protein | Combined 20aa Peptide |
|---|---|---|
| 14 | spike glycoprotein - | QCVNLTTRTQLPPAY |
| 23 | spike glycoprotein - | QLPPAYTNSFTRGVYYP |
| 39 | spike glycoprotein - | PDKVFRSSVLHSTQDL |
| 66 | spike glycoprotein - | HAIHVSGTNGTKRFDNPV |
| 87 | spike glycoprotein - | NDGVYFASTEKSNIIRGWIF |
| 111 | spike glycoprotein - | DSKTQSLLIVNNATN |
| 122 | spike glycoprotein - | NATNVVIKVCEFQFC |
| 141 | spike glycoprotein - | LGVYYHKNNKSWMESE |
| 175 | spike glycoprotein - | FLMDLEGKQGNFKNLR |
| 187 | spike glycoprotein - | KNLREFVFKNIDGYFKI |
| 208 | spike glycoprotein - | TPINLVRDLPQGFSALE |
| 229 | spike glycoprotein - | LPIGINITRFQTLLA |
| 233 | spike glycoprotein - | INITRFQTLLALHRS |
| 301 | spike glycoprotein - | CTLKSFTVEKGIYQTS |
| 313 | spike glycoprotein - | YQTSNFRVQPTESIVRFPN |
| 342 | spike glycoprotein - | FNATRFASVYAWNRK |
| 350 | spike glycoprotein - | VYAWNRKRISNCVADYSVL |
| 366 | spike glycoprotein - | SVLYNSASFSTFKCY |
| 372 | spike glycoprotein - | ASFSTFKCYGVSPTKLNDL |
| 400 | spike glycoprotein - | FVIRGDEVRQIAPGQT |
| 432 | spike glycoprotein - | CVIAWNSNNLDSKVGGN |
| 447 | spike glycoprotein - | GNYNYLYRLFRKSNLK |
| 509 | spike glycoprotein - | RVVVLSFELLHAPATVCGP |
| 528 | spike glycoprotein - | KKSTNLVKNKCVNFNFNGLT |
| 534 | spike glycoprotein - | VKNKCVNFNFNGLTGT |
| 579 | spike glycoprotein - | PQTLEILDITPCSFGGVS |
| 593 | spike glycoprotein - | GGVSVITPGTNTSNQVA |
| 614 | spike glycoprotein - | DVNCTEVPVAIHADQL |
| 634 | spike glycoprotein - | RVYSTGSNVFQTRAGC |
| 648 | spike glycoprotein - | GCLIGAEHVNNSYEC |
| 651 | spike glycoprotein - | IGAEHVNNSYECDIPI |
| 675 | spike glycoprotein - | QTQTNSPRRARSVASQSIIA |
| 689 | spike glycoprotein - | SQSIIAYTMSLGAENSVAY |
| 702 | spike glycoprotein - | ENSVAYSNNSIAIPTN |

| | | |
|---|---|---|
| 726 | spike glycoprotein - | ILPVSMTKTSVDCTMYI |
| 759 | spike glycoprotein - | FCTQLNRALTGIAVEQDK |
| 765 | spike glycoprotein - | RALTGIAVEQDKNTQ |
| 768 | spike glycoprotein - | TGIAVEQDKNTQEVF |
| 795 | spike glycoprotein - | KDFGGFNFSQILPDPSK |
| 802 | spike glycoprotein - | FSQILPDPSKPSKRSF |
| 817 | spike glycoprotein - | FIEDLLFNKVTLADAGFIKQ |
| 823 | spike glycoprotein - | FNKVTLADAGFIKQY |
| 853 | spike glycoprotein - | QKFNGLTVLPPLLTDEMI |
| 899 | spike glycoprotein - | AMQMAYRFNGIGVTQ |
| 903 | spike glycoprotein - | AYRFNGIGVTQNVLY |
| 913 | spike glycoprotein - | QNVLYENQKLIANQFNSAI |
| 920 | spike glycoprotein - | QKLIANQFNSAIGKIQDSLS |
| 926 | spike glycoprotein - | QFNSAIGKIQDSLSSTA |
| 934 | spike glycoprotein - | IQDSLSSTASALGKLQDVV |
| 942 | spike glycoprotein - | ASALGKLQDVVNQNAQALNT |
| 948 | spike glycoprotein - | LQDVVNQNAQALNTLVKQLS |
| 954 | spike glycoprotein - | QNAQALNTLVKQLSSNFGAI |
| 960 | spike glycoprotein - | NTLVKQLSSNFGAISS |
| 984 | spike glycoprotein - | LDKVEAEVQIDRLIT |
| 991 | spike glycoprotein - | VQIDRLITGRLQSLQTYVTQ |
| 997 | spike glycoprotein - | ITGRLQSLQTYVTQQLIRAA |
| 1010 | spike glycoprotein - | QQLIRAAEIRASANLAATK |
| 1044 | spike glycoprotein - | GKGYHLMSFPQSAPHGVVFL |
| 1050 | spike glycoprotein - | MSFPQSAPHGVVFLHVTYV |
| 1102 | spike glycoprotein - | WFVTQRNFYEPQIIT |
| 1105 | spike glycoprotein - | TQRNFYEPQIITTDN |
| 1168 | spike glycoprotein - | DISGINASVVNIQKE |
| 1222 | spike glycoprotein - | AGLIAIVMVTIMLCCMTS |
| 6 | ORF8 - | FLGIITTVAAFHQECSL |
| 47 | ORF8 - | IRVGARKSAPLIELCV |
| 53 | ORF8 - | KSAPLIELCVDEAGS |
| 78 | ORF8 - | NYTVSCLPFTINCQEPKLGS |
| 84 | ORF8 - | LPFTINCQEPKLGSLVVR |
| 8 | ORF7b - | DFYLCFLAFLLFLVLI |
| 25 | ORF7b - | LIIFWFSLELQDHNE |
| 28 | ORF7b - | FWFSLELQDHNETCHA |
| 8 | ORF7a - | ALITLATCELYHYQECV |
| 69 | ORF7a - | DGVKHVYQLRARSVSPK |
| 78 | ORF7a - | RARSVSPKLFIRQEEVQELY |
| 84 | ORF7a - | PKLFIRQEEVQELYSPIFL |
| 7 | ORF6 - | FQVTIAEILLIIMRTFKV |
| 34 | ORF6 - | NLIIKNLSKSLTENKYSQ |

| | | |
|---|---|---|
| 5 | ORF3a - | MRIFTIGTVTLKQGEIKDAT |
| 11 | ORF3a - | GTVTLKQGEIKDATP |
| 23 | ORF3a - | ATPSDFVRATATIPI |
| 26 | ORF3a - | SDFVRATATIPIQASL |
| 47 | ORF3a - | IVGVALLAVFQSASKIITLK |
| 53 | ORF3a - | LAVFQSASKIITLKKR |
| 64 | ORF3a - | TLKKRWQLALSKGVH |
| 116 | ORF3a - | QSINFVRIIMRLWLC |
| 195 | ORF3a - | SGVKDCVVLHSYFTSDY |
| 209 | ORF3a - | SDYYQLYSTQLSTDTGVEH |
| 232 | ORF3a - | IYNKIVDEPEEHVQI |
| 241 | ORF3a - | EEHVQIHTIDGSSGVVNP |
| 256 | ORF3a - | VNPVMEPIYDEPTTTTSVPL |
| 4411 | ORF1b polyprotein - | LTPCGTGTSTDVVYR |
| 4446 | ORF1b polyprotein - | CRFQEKDEDDNLIDS |
| 4465 | ORF1b polyprotein - | KRHTFSNYQHEETIYN |
| 4474 | ORF1b polyprotein - | HEETIYNLLKDCPAVAKHDF |
| 4480 | ORF1b polyprotein - | NLLKDCPAVAKHDFFK |
| 4499 | ORF1b polyprotein - | DGDMVPHISRQRLTK |
| 4545 | ORF1b polyprotein - | DDDYFNKKDWYDFVENPDIL |
| 4560 | ORF1b polyprotein - | NPDILRVYANLGERVRQALL |
| 4566 | ORF1b polyprotein - | VYANLGERVRQALLKTV |
| 4578 | ORF1b polyprotein - | LLKTVQFCDAMRNAGIVG |
| 4612 | ORF1b polyprotein - | GDFIQTTPGSGVPVVDS |
| 4624 | ORF1b polyprotein - | PVVDSYYSLLMPILTLTRA |
| 4633 | ORF1b polyprotein - | LMPILTLTRALTAESHVDT |
| 4682 | ORF1b polyprotein - | WDQTYHPNCVNCLDD |
| 4685 | ORF1b polyprotein - | TYHPNCVNCLDDRCI |
| 4723 | ORF1b polyprotein - | RKIFVDGVPFVVSTGYHF |
| 4741 | ORF1b polyprotein - | RELGVVHNQDVNLHSSRLSF |
| 4757 | ORF1b polyprotein - | RLSFKELLVYAADPA |
| 4783 | ORF1b polyprotein - | KRTTCFSVAALTNNVAFQT |
| 4792 | ORF1b polyprotein - | ALTNNVAFQTVKPGNFNKDF |
| 4798 | ORF1b polyprotein - | AFQTVKPGNFNKDFY |
| 4804 | ORF1b polyprotein - | PGNFNKDFYDFAVSKG |
| 4924 | ORF1b polyprotein - | KRNVIPTITQMNLKYAISAK |
| 4930 | ORF1b polyprotein - | TITQMNLKYAISAKNRARTV |
| 4936 | ORF1b polyprotein - | LKYAISAKNRARTVAGV |
| 4961 | ORF1b polyprotein - | RQFHQKLLKSIAATRGATVV |
| 4967 | ORF1b polyprotein - | LLKSIAATRGATVVIG |
| 4987 | ORF1b polyprotein - | YGGWHNMLKTVYSDV |
| 5015 | ORF1b polyprotein - | DRAMPNMLRIMASLV |
| 5041 | ORF1b polyprotein - | SHRFYRLANECAQVLS |

| | | |
|---|---|---|
| 5065 | ORF1b polyprotein - | LYVKPGGTSSGDATTAY |
| 5073 | ORF1b polyprotein - | SSGDATTAYANSVFNI |
| 5083 | ORF1b polyprotein - | NSVFNICQAVTANVNALLST |
| 5089 | ORF1b polyprotein - | CQAVTANVNALLSTD |
| 5148 | ORF1b polyprotein - | MILSDDAVVCFNSTY |
| 5152 | ORF1b polyprotein - | DDAVVCFNSTYASQGLVASI |
| 5158 | ORF1b polyprotein - | FNSTYASQGLVASIKN |
| 5166 | ORF1b polyprotein - | GLVASIKNFKSVLYYQNNV |
| 5176 | ORF1b polyprotein - | SVLYYQNNVFMSEAK |
| 5201 | ORF1b polyprotein - | PHEFCSQHTMLVKQGDD |
| 5234 | ORF1b polyprotein - | CFVDDIVKTDGTLMI |
| 5237 | ORF1b polyprotein - | DDIVKTDGTLMIERFVSLAI |
| 5243 | ORF1b polyprotein - | DGTLMIERFVSLAIDAYPL |
| 5253 | ORF1b polyprotein - | SLAIDAYPLTKHPNQ |
| 5260 | ORF1b polyprotein - | PLTKHPNQEYADVFH |
| 5358 | ORF1b polyprotein - | VISTSHKLVLSVNPYVCNAP |
| 5364 | ORF1b polyprotein - | KLVLSVNPYVCNAPGCD |
| 5431 | ORF1b polyprotein - | NAIATCDWTNAGDYI |
| 5444 | ORF1b polyprotein - | YILANTCTERLKLFAAETLK |
| 5450 | ORF1b polyprotein - | CTERLKLFAAETLKAT |
| 5498 | ORF1b polyprotein - | PPLNRNYVFTGYRVT |
| 5504 | ORF1b polyprotein - | YVFTGYRVTKNSKVQIG |
| 5551 | ORF1b polyprotein - | LTSHTVMPLSAPTLVPQEHY |
| 5568 | ORF1b polyprotein - | EHYVRITGLYPTLNI |
| 5596 | ORF1b polyprotein - | VGMQKYSTLQGPPGTG |
| 5626 | ORF1b polyprotein - | ARIVYTACSHAAVDAL |
| 5640 | ORF1b polyprotein - | ALCEKALKYLPIDKCS |
| 5661 | ORF1b polyprotein - | RARVECFDKFKVNSTLE |
| 5681 | ORF1b polyprotein - | FCTVNALPETTADIVVFD |
| 5697 | ORF1b polyprotein - | FDEISMATNYDLSVVNAR |
| 5718 | ORF1b polyprotein - | KHYVYIGDPAQLPAPR |
| 5747 | ORF1b polyprotein - | NSVCRLMKTIGPDMFL |
| 5769 | ORF1b polyprotein - | PAEIVDTVSALVYDNKLKA |
| 5794 | ORF1b polyprotein - | QCFKMFYKGVITHDV |
| 5797 | ORF1b polyprotein - | KMFYKGVITHDVSSA |
| 5801 | ORF1b polyprotein - | KGVITHDVSSAINRPQIG |
| 5807 | ORF1b polyprotein - | DVSSAINRPQIGVVRE |
| 5828 | ORF1b polyprotein - | PAWRKAVFISPYNSQNAVAS |
| 5834 | ORF1b polyprotein - | VFISPYNSQNAVASKIL |
| 5840 | ORF1b polyprotein - | NSQNAVASKILGLPTQTVDS |
| 5846 | ORF1b polyprotein - | ASKILGLPTQTVDSS |
| 5850 | ORF1b polyprotein - | LGLPTQTVDSSQGSEY |
| 5854 | ORF1b polyprotein - | TQTVDSSQGSEYDYV |

| | | |
|---|---|---|
| 5870 | ORF1b polyprotein - | FTQTTETAHSCNVNR |
| 5887 | ORF1b polyprotein - | VAITRAKVGILCIMS |
| 5895 | ORF1b polyprotein - | GILCIMSDRDLYDKL |
| 5902 | ORF1b polyprotein - | DRDLYDKLQFTSLEIPR |
| 5923 | ORF1b polyprotein - | TLQAENVTGLFKDCSK |
| 5927 | ORF1b polyprotein - | ENVTGLFKDCSKVIT |
| 5937 | ORF1b polyprotein - | SKVITGLHPTQAPTHLSVDT |
| 5943 | ORF1b polyprotein - | LHPTQAPTHLSVDTKF |
| 5973 | ORF1b polyprotein - | DMTYRRLISMMGFKM |
| 5976 | ORF1b polyprotein - | YRRLISMMGFKMNYQVNGYP |
| 5982 | ORF1b polyprotein - | MMGFKMNYQVNGYPNMFITR |
| 5988 | ORF1b polyprotein - | NYQVNGYPNMFITRE |
| 6000 | ORF1b polyprotein - | TREEAIRHVRAWIGFDVEGC |
| 6034 | ORF1b polyprotein - | LGFSTGVNLVAVPTGYVDTP |
| 6040 | ORF1b polyprotein - | VNLVAVPTGYVDTPNNTDF |
| 6055 | ORF1b polyprotein - | NTDFSRVSAKPPPGD |
| 6094 | ORF1b polyprotein - | MLSDTLKNLSDRVVFV |
| 6154 | ORF1b polyprotein - | HSIGFDYVYNPFMIDVQQW |
| 6193 | ORF1b polyprotein - | HVASCDAIMTRCLAV |
| 6249 | ORF1b polyprotein - | DKFPVLHDIGNPKAIKCVPQ |
| 6259 | ORF1b polyprotein - | NPKAIKCVPQADVEWKFYD |
| 6292 | ORF1b polyprotein - | FYSYATHSDKFTDGV |
| 6327 | ORF1b polyprotein - | DTRVLSNLNLPGCDGGS |
| 6353 | ORF1b polyprotein - | TPAFDKSAFVNLKQLPFFYY |
| 6359 | ORF1b polyprotein - | SAFVNLKQLPFFYYS |
| 6373 | ORF1b polyprotein - | SDSPCESHGKQVVSD |
| 6440 | ORF1b polyprotein - | DTYNLWNTFTRLQSLENVAF |
| 6446 | ORF1b polyprotein - | NTFTRLQSLENVAFNVVNKG |
| 6452 | ORF1b polyprotein - | QSLENVAFNVVNKGH |
| 6455 | ORF1b polyprotein - | ENVAFNVVNKGHFDGQQ |
| 6471 | ORF1b polyprotein - | QGEVPVSIINNTVYT |
| 6477 | ORF1b polyprotein - | SIINNTVYTKVDGVD |
| 6483 | ORF1b polyprotein - | VYTKVDGVDVELFENKT |
| 6514 | ORF1b polyprotein - | NIKPVPEVKILNNLGVDIAA |
| 6520 | ORF1b polyprotein - | EVKILNNLGVDIAANT |
| 6531 | ORF1b polyprotein - | IAANTVIWDYKRDAP |
| 6542 | ORF1b polyprotein - | RDAPAHISTIGVCSMTDIAK |
| 6548 | ORF1b polyprotein - | ISTIGVCSMTDIAKKPTETI |
| 6576 | ORF1b polyprotein - | DGRVDGQVDLFRNARN |
| 6595 | ORF1b polyprotein - | ITEGSVKGLQPSVGPKQASL |
| 6601 | ORF1b polyprotein - | KGLQPSVGPKQASLNGVTLI |
| 6607 | ORF1b polyprotein - | VGPKQASLNGVTLIGE |
| 6625 | ORF1b polyprotein - | KTQFNYYKKVDGVVQQ |

| | | |
|---|---|---|
| 6641 | ORF1b polyprotein - | LPETYFTQSRNLQEFKP |
| 6654 | ORF1b polyprotein - | EFKPRSQMEIDFLEL |
| 6664 | ORF1b polyprotein - | DFLELAMDEFIERYKLE |
| 6699 | ORF1b polyprotein - | GLHLLIGLAKRFKESPF |
| 6724 | ORF1b polyprotein - | DSTVKNYFITDAQTGSSKCV |
| 6730 | ORF1b polyprotein - | YFITDAQTGSSKCVCS |
| 6753 | ORF1b polyprotein - | DFVEIIKSQDLSVVSKVVK |
| 6829 | ORF1b polyprotein - | GDSATLPKGIMMNVAKYT |
| 6850 | ORF1b polyprotein - | QYLNTLTLAVPYNMRVIHF |
| 6902 | ORF1b polyprotein - | VSDADSTLIGDCATVHT |
| 6914 | ORF1b polyprotein - | ATVHTANKWDLIISDMY |
| 6938 | ORF1b polyprotein - | TKENDSKEGFFTYIC |
| 6956 | ORF1b polyprotein - | QQKLALGGSVAIKITEHSWN |
| 6962 | ORF1b polyprotein - | GGSVAIKITEHSWNAD |
| 6999 | ORF1b polyprotein - | SSEAFLIGCNYLGKPREQI |
| 7022 | ORF1b polyprotein - | MHANYIFWRNTNPIQLSSY |
| 7044 | ORF1b polyprotein - | DMSKFPLKLRGTAVMSLKEG |
| 7050 | ORF1b polyprotein - | LKLRGTAVMSLKEGQI |
| 7055 | ORF1b polyprotein - | TAVMSLKEGQINDMI |
| 7073 | ORF1b polyprotein - | LSKGRLIIRENNRVVISSDV |
| 7079 | ORF1b polyprotein - | IIRENNRVVISSDVLVNN |
| 6 | ORF1a polyprotein - | PGFNEKTHVQLSLPV |
| 17 | ORF1a polyprotein - | SLPVLQVRDVLVRGFGDSVE |
| 23 | ORF1a polyprotein - | VRDVLVRGFGDSVEE |
| 57 | ORF1a polyprotein - | EKGVLPQLEQPYVFI |
| 65 | ORF1a polyprotein - | EQPYVFIKRSDARTA |
| 92 | ORF1a polyprotein - | LEGIQYGRSGETLGVLVP |
| 100 | ORF1a polyprotein - | SGETLGVLVPHVGEIPVAYR |
| 119 | ORF1a polyprotein - | RKVLLRKNGNKGAGGH |
| 136 | ORF1a polyprotein - | YGADLKSFDLGDELGTDPY |
| 147 | ORF1a polyprotein - | DELGTDPYEDFQENW |
| 169 | ORF1a polyprotein - | VTRELMRELNGGAYTRYVD |
| 195 | ORF1a polyprotein - | GYPLECIKDLLARAGKASC |
| 251 | ORF1a polyprotein - | LQTPFEIKLAKKFDT |
| 254 | ORF1a polyprotein - | PFEIKLAKKFDTFNGE |
| 281 | ORF1a polyprotein - | IKTIQPRVEKKKLDG |
| 290 | ORF1a polyprotein - | KKKLDGFMGRIRSVY |
| 296 | ORF1a polyprotein - | FMGRIRSVYPVASPNECN |
| 387 | ORF1a polyprotein - | YHNESGLKTILRKGG |
| 390 | ORF1a polyprotein - | ESGLKTILRKGGRTI |
| 442 | ORF1a polyprotein - | GSEGLNDNLLEILQK |
| 459 | ORF1a polyprotein - | VNINIVGDFKLNEEI |
| 476 | ORF1a polyprotein - | ILASFSASTSAFVETVK |

| | | |
|---|---|---|
| 488 | ORF1a polyprotein - | VETVKGLDYKAFKQIV |
| 501 | ORF1a polyprotein - | QIVESCGNFKVTKGKAKKGA |
| 507 | ORF1a polyprotein - | GNFKVTKGKAKKGAWN |
| 554 | ORF1a polyprotein - | TAQNSVRVLQKAAITI |
| 562 | ORF1a polyprotein - | LQKAAITILDGISQYSLR |
| 571 | ORF1a polyprotein - | DGISQYSLRLIDAMMFTSDL |
| 577 | ORF1a polyprotein - | SLRLIDAMMFTSDLA |
| 594 | ORF1a polyprotein - | NLVVMAYITGGVVQLT |
| 646 | ORF1a polyprotein - | WEIVKFISTCACEIVGGQ |
| 682 | ORF1a polyprotein - | VNKFLALCADSIIIGGAKL |
| 699 | ORF1a polyprotein - | KLKALNLGETFVTHSKGLYR |
| 705 | ORF1a polyprotein - | LGETFVTHSKGLYRK |
| 730 | ORF1a polyprotein - | LMPLKAPKEIIFLEG |
| 740 | ORF1a polyprotein - | IFLEGETLPTEVLTEE |
| 751 | ORF1a polyprotein - | VLTEEVVLKTGDLQP |
| 834 | ORF1a polyprotein - | QGYKSVNITFELDERID |
| 858 | ORF1a polyprotein - | SAYTVELGTEVNEFAC |
| 867 | ORF1a polyprotein - | EVNEFACVVADAVIKT |
| 879 | ORF1a polyprotein - | VIKTLQPVSELLTPLGIDLD |
| 955 | ORF1a polyprotein - | YQGKPLEFGATSAALQPEEE |
| 961 | ORF1a polyprotein - | EFGATSAALQPEEEQ |
| 994 | ORF1a polyprotein - | SEDNQTTTIQTIVEVQ |
| 1011 | ORF1a polyprotein - | QLEMELTPVVQTIEVNSF |
| 1032 | ORF1a polyprotein - | LKLTDNVYIKNADIV |
| 1043 | ORF1a polyprotein - | ADIVEEAKKVKPTVVVNAAN |
| 1049 | ORF1a polyprotein - | AKKVKPTVVVNAANVYLK |
| 1114 | ORF1a polyprotein - | CLHVVGPNVNKGEDIQLLKS |
| 1120 | ORF1a polyprotein - | PNVNKGEDIQLLKSA |
| 1126 | ORF1a polyprotein - | EDIQLLKSAYENFNQHEV |
| 1142 | ORF1a polyprotein - | EVLLAPLLSAGIFGADPIHS |
| 1148 | ORF1a polyprotein - | LLSAGIFGADPIHSLR |
| 1183 | ORF1a polyprotein - | YDKLVSSFLEMKSEKQVE |
| 1216 | ORF1a polyprotein - | TESKPSVEQRKQDDKK |
| 1228 | ORF1a polyprotein - | DDKKIKACVEEVTTT |
| 1276 | ORF1a polyprotein - | ITFLKKDAPYIVGDVVQE |
| 1285 | ORF1a polyprotein - | YIVGDVVQEGVLTAVVIPT |
| 1293 | ORF1a polyprotein - | EGVLTAVVIPTKKAGGTTEM |
| 1299 | ORF1a polyprotein - | VVIPTKKAGGTTEML |
| 1308 | ORF1a polyprotein - | GTTEMLAKALRKVPTDNYI |
| 1346 | ORF1a polyprotein - | LKKCKSAFYILPSIISNEKQ |
| 1352 | ORF1a polyprotein - | AFYILPSIISNEKQEILGTV |
| 1358 | ORF1a polyprotein - | SIISNEKQEILGTVSWNLR |
| 1377 | ORF1a polyprotein - | EMLAHAEETRKLMPVCVETK |

| | | |
|---|---|---|
| 1383 | ORF1a polyprotein - | EETRKLMPVCVETKAI |
| 1391 | ORF1a polyprotein - | VCVETKAIVSTIQRKYKGIK |
| 1397 | ORF1a polyprotein - | AIVSTIQRKYKGIKIQEG |
| 1422 | ORF1a polyprotein - | FYFYTSKTTVASLINTLNDL |
| 1428 | ORF1a polyprotein - | KTTVASLINTLNDLNETLVT |
| 1453 | ORF1a polyprotein - | VTHGLNLEEAARYMRS |
| 1460 | ORF1a polyprotein - | EEAARYMRSLKVPATVSVSS |
| 1466 | ORF1a polyprotein - | MRSLKVPATVSVSSP |
| 1498 | ORF1a polyprotein - | EEHFIETISLAGSYKDWSY |
| 1527 | ORF1a polyprotein - | FLKRGDKSVYYTSNP |
| 1534 | ORF1a polyprotein - | SVYYTSNPTTFHLDGE |
| 1561 | ORF1a polyprotein - | SLREVRTIKVFTTVDNINL |
| 1577 | ORF1a polyprotein - | INLHTQVVDMSMTYGQQ |
| 1583 | ORF1a polyprotein - | VVDMSMTYGQQFGPT |
| 1631 | ORF1a polyprotein - | AFEYYHTTDPSFLGRYM |
| 1639 | ORF1a polyprotein - | DPSFLGRYMSALNHTKKWKY |
| 1645 | ORF1a polyprotein - | RYMSALNHTKKWKYP |
| 1655 | ORF1a polyprotein - | KWKYPQVNGLTSIKWA |
| 1711 | ORF1a polyprotein - | CALILAYCNKTVGELGDV |
| 1773 | ORF1a polyprotein - | TLSYEQFKKGVQIPC |
| 1795 | ORF1a polyprotein - | KYLVQQESPFVMMSAPPAQY |
| 1801 | ORF1a polyprotein - | ESPFVMMSAPPAQYELKH |
| 1826 | ORF1a polyprotein - | EYTGNYQCGHYKHIT |
| 1839 | ORF1a polyprotein - | ITSKETLYCIDGALL |
| 1875 | ORF1a polyprotein - | TTIKPVTYKLDGVVCTEID |
| 1894 | ORF1a polyprotein - | PKLDNYYKKDNSYFTE |
| 1900 | ORF1a polyprotein - | YKKDNSYFTEQPIDL |
| 1914 | ORF1a polyprotein - | LVPNQPYPNASFDNF |
| 1926 | ORF1a polyprotein - | DNFKFVCDNIKFADD |
| 1961 | ORF1a polyprotein - | PDLNGDVVAIDYKHY |
| 2016 | ORF1a polyprotein - | TKPVETSNSFDVLKSEDAQG |
| 2022 | ORF1a polyprotein - | SNSFDVLKSEDAQGMD |
| 2026 | ORF1a polyprotein - | DVLKSEDAQGMDNLAC |
| 2043 | ORF1a polyprotein - | DLKPVSEEVVENPTIQKDVL |
| 2066 | ORF1a polyprotein - | VKTTEVVGDIILKPA |
| 2071 | ORF1a polyprotein - | VVGDIILKPANNSLKITEEV |
| 2077 | ORF1a polyprotein - | LKPANNSLKITEEVGHT |
| 2093 | ORF1a polyprotein - | TDLMAAYVDNSSLTI |
| 2109 | ORF1a polyprotein - | KPNELSRVLGLKTLATHGLA |
| 2115 | ORF1a polyprotein - | RVLGLKTLATHGLAAVNSVP |
| 2136 | ORF1a polyprotein - | DTIANYAKPFLNKVV |
| 2143 | ORF1a polyprotein - | KPFLNKVVSTTTNIVTRC |
| 2175 | ORF1a polyprotein - | LLLQLCTFTRSTNSRIKASM |

| | | |
|---|---|---|
| 2181 | ORF1a polyprotein - | TFTRSTNSRIKASMPTTI |
| 2240 | ORF1a polyprotein - | LGSLIYSTAALGVLMSNLGM |
| 2246 | ORF1a polyprotein - | STAALGVLMSNLGMPSY |
| 2266 | ORF1a polyprotein - | YREGYLNSTNVTIAT |
| 2274 | ORF1a polyprotein - | TNVTIATYCTGSIPCS |
| 2290 | ORF1a polyprotein - | VCLSGLDSLDTYPSL |
| 2299 | ORF1a polyprotein - | DTYPSLETIQITISS |
| 2310 | ORF1a polyprotein - | TISSFKWDLTAFGLVAE |
| 2418 | ORF1a polyprotein - | ATRVECTTIVNGVRRSF |
| 2470 | ORF1a polyprotein - | ARDLSLQFKRPINPTDQSSY |
| 2476 | ORF1a polyprotein - | QFKRPINPTDQSSYI |
| 2513 | ORF1a polyprotein - | YERHSLSHFVNLDNLRANNT |
| 2519 | ORF1a polyprotein - | SHFVNLDNLRANNTK |
| 2564 | ORF1a polyprotein - | LMCQPILLLDQALVSDVGDS |
| 2570 | ORF1a polyprotein - | LLLDQALVSDVGDSAE |
| 2584 | ORF1a polyprotein - | AEVAVKMFDAYVNTFSSTFN |
| 2590 | ORF1a polyprotein - | MFDAYVNTFSSTFNVPM |
| 2607 | ORF1a polyprotein - | EKLKTLVATAEAELAKNVSL |
| 2613 | ORF1a polyprotein - | VATAEAELAKNVSLD |
| 2650 | ORF1a polyprotein - | DVVECLKLSHQSDIEVTG |
| 2695 | ORF1a polyprotein - | SARHINAQVAKSHNIALI |
| 2708 | ORF1a polyprotein - | NIALIWNVKDFMSLSEQLRK |
| 2714 | ORF1a polyprotein - | NVKDFMSLSEQLRKQ |
| 2743 | ORF1a polyprotein - | TCATTRQVVNVVTTKI |
| 2747 | ORF1a polyprotein - | TRQVVNVVTTKIALKGGKIV |
| 2755 | ORF1a polyprotein - | TTKIALKGGKIVNNW |
| 2851 | ORF1a polyprotein - | CPLIAAVITREVGFVVP |
| 2884 | ORF1a polyprotein - | FLPRVFSAVGNICYT |
| 2891 | ORF1a polyprotein - | AVGNICYTPSKLIEYTDFA |
| 2904 | ORF1a polyprotein - | EYTDFATSACVLAAECT |
| 2954 | ORF1a polyprotein - | YVLMDGSIIQFPNTYLEGSV |
| 3013 | ORF1a polyprotein - | SLPGVFCGVDAVNLL |
| 3022 | ORF1a polyprotein - | DAVNLLTNMFTPLIQPIGA |
| 3032 | ORF1a polyprotein - | TPLIQPIGALDISASIVAGG |
| 3038 | ORF1a polyprotein - | IGALDISASIVAGGIVAI |
| 3066 | ORF1a polyprotein - | RFRRAFGEYSHVVAFNTLL |
| 3129 | ORF1a polyprotein - | MVMFTPLVPFWITIA |
| 3157 | ORF1a polyprotein - | FSNYLKRRVVFNGVSFSTFE |
| 3163 | ORF1a polyprotein - | RRVVFNGVSFSTFEE |
| 3189 | ORF1a polyprotein - | MYLKLRSDVLLPLTQYN |
| 3211 | ORF1a polyprotein - | YNKYKYFSGAMDTTSYRE |
| 3232 | ORF1a polyprotein - | CHLAKALNDFSNSGSDVLYQ |
| 3238 | ORF1a polyprotein - | LNDFSNSGSDVLYQPP |

| | | |
|---|---|---|
| 3248 | ORF1a polyprotein - | VLYQPPQTSITSAVLQSG |
| 3346 | ORF1a polyprotein - | QNCVLKLKVDTANPKTPKY |
| 3361 | ORF1a polyprotein - | TPKYKFVRIQPGQTFSVLAC |
| 3367 | ORF1a polyprotein - | VRIQPGQTFSVLACYN |
| 3422 | ORF1a polyprotein - | FCYMHHMELPTGVHAG |
| 3446 | ORF1a polyprotein - | GPFVDRQTAQAAGTDT |
| 3490 | ORF1a polyprotein - | LNDFNLVAMKYNYEPLTQDH |
| 3496 | ORF1a polyprotein - | VAMKYNYEPLTQDHV |
| 3508 | ORF1a polyprotein - | DHVDILGPLSAQTGIAVLD |
| 3522 | ORF1a polyprotein - | IAVLDMCASLKELLQ |
| 3535 | ORF1a polyprotein - | LQNGMNGRTILGSAL |
| 3547 | ORF1a polyprotein - | SALLEDEFTPFDVVRQCSGV |
| 3556 | ORF1a polyprotein - | PFDVVRQCSGVTFQSAVK |
| 3640 | ORF1a polyprotein - | LLPSLATVAYFNMVY |
| 3643 | ORF1a polyprotein - | SLATVAYFNMVYMPASWVMR |
| 3649 | ORF1a polyprotein - | YFNMVYMPASWVMRIM |
| 3662 | ORF1a polyprotein - | RIMTWLDMVDTSLSG |
| 3665 | ORF1a polyprotein - | TWLDMVDTSLSGFKL |
| 3724 | ORF1a polyprotein - | GNALDQAISMWALIISVTS |
| 3840 | ORF1a polyprotein - | LNIKLLGVGGKPCIK |
| 3846 | ORF1a polyprotein - | GVGGKPCIKVATVQSKMS |
| 3870 | ORF1a polyprotein - | VVLLSVLQQLRVESSS |
| 3891 | ORF1a polyprotein - | CVQLHNDILLAKDTT |
| 3897 | ORF1a polyprotein - | DILLAKDTTEAFEKMV |
| 3943 | ORF1a polyprotein - | AIASEFSSLPSYAAFATAQE |
| 3949 | ORF1a polyprotein - | SSLPSYAAFATAQEAYEQ |
| 3960 | ORF1a polyprotein - | AQEAYEQAVANGDSEVV |
| 3971 | ORF1a polyprotein - | GDSEVVLKKLKKSLN |
| 4001 | ORF1a polyprotein - | LEKMADQAMTQMYKQARSE |
| 4026 | ORF1a polyprotein - | TSAMQTMLFTMLRKLDNDAL |
| 4032 | ORF1a polyprotein - | MLFTMLRKLDNDALNNIIN |
| 4056 | ORF1a polyprotein - | CVPLNIIPLTTAAKLMVVIP |
| 4073 | ORF1a polyprotein - | VIPDYNTYKNTCDGTTFTY |
| 4097 | ORF1a polyprotein - | EIQQVVDADSKIVQLS |
| 4106 | ORF1a polyprotein - | SKIVQLSEISMDNSP |
| 4126 | ORF1a polyprotein - | LIVTALRANSAVKLQNNELS |
| 4134 | ORF1a polyprotein - | NSAVKLQNNELSPVALRQMS |
| 4140 | ORF1a polyprotein - | QNNELSPVALRQMSCA |
| 4144 | ORF1a polyprotein - | LSPVALRQMSCAAGTTQTAC |
| 4150 | ORF1a polyprotein - | RQMSCAAGTTQTACTDDN |
| 4165 | ORF1a polyprotein - | DDNALAYYNTTKGGRFVLAL |
| 4171 | ORF1a polyprotein - | YYNTTKGGRFVLALL |
| 4181 | ORF1a polyprotein - | VLALLSDLQDLKWARFP |

| | | |
|---|---|---|
| 4204 | ORF1a polyprotein - | TIYTELEPPCRFVTDTPKG |
| 4216 | ORF1a polyprotein - | VTDTPKGPKVKYLYFI |
| 4229 | ORF1a polyprotein - | YFIKGLNNLNRGMVLGSL |
| 4240 | ORF1a polyprotein - | GMVLGSLAATVRLQAG |
| 4278 | ORF1a polyprotein - | KAYKDYLASGGQPIT |
| 4294 | ORF1a polyprotein - | CVKMLCTHTGTGQAIT |
| 4384 | ORF1a polyprotein - | DQLREPMLQSADAQSFLNRV |
| 13 | ORF10 | IYSLLLCRMNSRNYIAQ |
| 23 | ORF10 | SRNYIAQVDVVNFNLT |
| 16 | nucleocapsid phosphoprotein - | TFGGPSDSTGSNQNGER |
| 41 | nucleocapsid phosphoprotein - | RPQGLPNNTASWFTALTQ |
| 50 | nucleocapsid phosphoprotein - | ASWFTALTQHGKEDL |
| 70 | nucleocapsid phosphoprotein - | QGVPINTNSSPDDQI |
| 83 | nucleocapsid phosphoprotein - | QIGYYRRATRRIRGG |
| 113 | nucleocapsid phosphoprotein - | LGTGPEAGLPYGANKD |
| 121 | nucleocapsid phosphoprotein - | LPYGANKDGIIWVATEGALN |
| 127 | nucleocapsid phosphoprotein - | KDGIIWVATEGALNTPKDHI |
| 133 | nucleocapsid phosphoprotein - | VATEGALNTPKDHIGTRN |
| 163 | nucleocapsid phosphoprotein - | QGTTLPKGFYAEGSRGGSQ |
| 221 | nucleocapsid phosphoprotein - | LLLLDRLNQLESKMSG |
| 225 | nucleocapsid phosphoprotein - | DRLNQLESKMSGKGQ |
| 228 | nucleocapsid phosphoprotein - | NQLESKMSGKGQQQQ |
| 262 | nucleocapsid phosphoprotein - | RTATKAYNVTQAFGRRGPE |
| 300 | nucleocapsid phosphoprotein - | HWPQIAQFAPSASAFFGMSR |
| 306 | nucleocapsid phosphoprotein - | QFAPSASAFFGMSRI |
| 309 | nucleocapsid phosphoprotein - | PSASAFFGMSRIGMEVTPSG |
| 315 | nucleocapsid phosphoprotein - | FGMSRIGMEVTPSGTWLTYT |
| 321 | nucleocapsid phosphoprotein - | GMEVTPSGTWLTYTGAIK |
| 396 | nucleocapsid phosphoprotein - | PAADLDDFSKQLQQS |
| 19 | membrane glycoprotein - | QWNLVIGFLFLTWICLLQF |
| 44 | membrane glycoprotein - | RFLYIIKLIFLWLLW |
| 98 | membrane glycoprotein - | ASFRLFARTRSMWSFNPE |
| 114 | membrane glycoprotein - | PETNILLNVPLHGTILTRPL |
| 120 | membrane glycoprotein - | LNVPLHGTILTRPLLES |
| 144 | membrane glycoprotein - | ILRGHLRIAGHHLGR |
| 147 | membrane glycoprotein - | GHLRIAGHHLGRCDIK |
| 161 | membrane glycoprotein - | IKDLPKEITVATSRTLSY |
| 186 | membrane glycoprotein - | RVAGDSGFAAYSRYRIGNYK |
| 201 | membrane glycoprotein - | IGNYKLNTDHSSSSDN |
| 35 | envelope protein - | TALRLCAYCCNIVNV |
| 43 | envelope protein - | CCNIVNVSLVKPSFY |
| 54 | envelope protein - | PSFYVYSRVKNLNSSRVPDL |
| 60 | envelope protein - | SRVKNLNSSRVPDLLV |

**DRB11501**

| Position in Protein | | Combined 20aa Peptide |
|---|---|---|
| 21 | spike glycoprotein - | RTQLPPAYTNSFTRG |
| 40 | spike glycoprotein - | DKVFRSSVLHSTQDLF |
| 59 | spike glycoprotein - | FSNVTWFHAIHVSGTNGT |
| 69 | spike glycoprotein - | HVSGTNGTKRFDNPVLP |
| 194 | spike glycoprotein - | FKNIDGYFKIYSKHTPINLV |
| 200 | spike glycoprotein - | YFKIYSKHTPINLVRDLP |
| 210 | spike glycoprotein - | INLVRDLPQGFSALE |
| 230 | spike glycoprotein - | PIGINITRFQTLLALHRSYL |
| 236 | spike glycoprotein - | TRFQTLLALHRSYLTPGDSS |
| 242 | spike glycoprotein - | LALHRSYLTPGDSSSGW |
| 304 | spike glycoprotein - | KSFTVEKGIYQTSNF |
| 315 | spike glycoprotein - | TSNFRVQPTESIVRFPNIT |
| 337 | spike glycoprotein - | PFGEVFNATRFASVYAWNRK |
| 343 | spike glycoprotein - | NATRFASVYAWNRKRISNCV |
| 349 | spike glycoprotein - | SVYAWNRKRISNCVADY |
| 399 | spike glycoprotein - | SFVIRGDEVRQIAPGQTGK |
| 453 | spike glycoprotein - | YRLFRKSNLKPFERDI |
| 546 | spike glycoprotein - | LTGTGVLTESNKKFLPFQQF |
| 552 | spike glycoprotein - | LTESNKKFLPFQQFGRDIAD |
| 558 | spike glycoprotein - | KFLPFQQFGRDIADT |
| 632 | spike glycoprotein - | TWRVYSTGSNVFQTR |
| 682 | spike glycoprotein - | RRARSVASQSIIAYTMSLGA |
| 688 | spike glycoprotein - | ASQSIIAYTMSLGAENSVAY |
| 722 | spike glycoprotein - | VTTEILPVSMTKTSVDC |
| 769 | spike glycoprotein - | GIAVEQDKNTQEVFAQ |
| 773 | spike glycoprotein - | EQDKNTQEVFAQVKQ |
| 778 | spike glycoprotein - | TQEVFAQVKQIYKTPPIKDF |
| 784 | spike glycoprotein - | QVKQIYKTPPIKDFGGFN |
| 802 | spike glycoprotein - | FSQILPDPSKPSKRSFIEDL |
| 868 | spike glycoprotein - | EMIAQYTSALLAGTI |
| 894 | spike glycoprotein - | LQIPFAMQMAYRFNGIGVTQ |
| 900 | spike glycoprotein - | MQMAYRFNGIGVTQNVL |
| 910 | spike glycoprotein - | GVTQNVLYENQKLIANQFNS |
| 919 | spike glycoprotein - | NQKLIANQFNSAIGKIQDSL |
| 925 | spike glycoprotein - | NQFNSAIGKIQDSLS |
| 928 | spike glycoprotein - | NSAIGKIQDSLSSTA |
| 936 | spike glycoprotein - | DSLSSTASALGKLQDVVNQN |
| 942 | spike glycoprotein - | ASALGKLQDVVNQNAQAL |
| 949 | spike glycoprotein - | QDVVNQNAQALNTLVKQLS |

| | | |
|---|---|---|
| 959 | spike glycoprotein - | LNTLVKQLSSNFGAISSVL |
| 983 | spike glycoprotein - | RLDKVEAEVQIDRLITG |
| 994 | spike glycoprotein - | DRLITGRLQSLQTYVTQQLI |
| 1000 | spike glycoprotein - | RLQSLQTYVTQQLIRAAEIR |
| 1006 | spike glycoprotein - | TYVTQQLIRAAEIRASANLA |
| 1012 | spike glycoprotein - | LIRAAEIRASANLAATKMSE |
| 1064 | spike glycoprotein - | HVTYVPAQEKNFTTA |
| 1173 | spike glycoprotein - | NASVVNIQKEIDRLN |
| 1219 | spike glycoprotein - | GFIAGLIAIVMVTIM |
| 1222 | spike glycoprotein - | AGLIAIVMVTIMLCCMTSC |
| 1 | ORF8 - | MKFLVFLGIITTVAA |
| 33 | ORF8 - | VDDPCPIHFYSKWYIRVGAR |
| 39 | ORF8 - | IHFYSKWYIRVGARKSAPLI |
| 45 | ORF8 - | WYIRVGARKSAPLIELCV |
| 6 | ORF7b - | LIDFYLCFLAFLLFL |
| 18 | ORF7b - | LFLVLIMLIIFWFSLELQ |
| 1 | ORF7a - | MKIILFLALITLATCE |
| 68 | ORF7a - | PDGVKHVYQLRARSVSPKLF |
| 74 | ORF7a - | VYQLRARSVSPKLFIRQEEV |
| 80 | ORF7a - | RSVSPKLFIRQEEVQE |
| 31 | ORF6 - | YIINLIIKNLSKSLTENKYS |
| 37 | ORF6 - | IKNLSKSLTENKYSQLD |
| 5 | ORF3a - | MRIFTIGTVTLKQGEI |
| 10 | ORF3a - | IGTVTLKQGEIKDAT |
| 24 | ORF3a - | TPSDFVRATATIPIQASLPF |
| 30 | ORF3a - | RATATIPIQASLPFG |
| 39 | ORF3a - | ASLPFGWLIVGVALL |
| 48 | ORF3a - | VGVALLAVFQSASKIITLKK |
| 54 | ORF3a - | AVFQSASKIITLKKRWQLAL |
| 60 | ORF3a - | SKIITLKKRWQLALSKGVHF |
| 66 | ORF3a - | KKRWQLALSKGVHFV |
| 73 | ORF3a - | LSKGVHFVCNLLLLFVT |
| 87 | ORF3a - | FVTVYSHLLLVAAGLEA |
| 179 | ORF3a - | ISEHDYQIGGYTEKW |
| 213 | ORF3a - | QLYSTQLSTDTGVEHV |
| 4457 | ORF1b polyprotein - | LIDSYFVVKRHTFSNYQHEE |
| 4463 | ORF1b polyprotein - | VVKRHTFSNYQHEETI |
| 4497 | ORF1b polyprotein - | RIDGDMVPHISRQRLTKYTM |
| 4503 | ORF1b polyprotein - | VPHISRQRLTKYTMADLVY |
| 4560 | ORF1b polyprotein - | NPDILRVYANLGERVRQALL |
| 4633 | ORF1b polyprotein - | LMPILTLTRALTAESHVDTD |
| 4639 | ORF1b polyprotein - | LTRALTAESHVDTDLT |
| 4748 | ORF1b polyprotein - | NQDVNLHSSRLSFKEL |

| | | |
|---|---|---|
| 4892 | ORF1b polyprotein - | KSAGFPFNKWGKARLYYDSM |
| 4898 | ORF1b polyprotein - | FNKWGKARLYYDSMS |
| 4912 | ORF1b polyprotein - | SYEDQDALFAYTKRNVIPTI |
| 4918 | ORF1b polyprotein - | ALFAYTKRNVIPTITQMNL |
| 4925 | ORF1b polyprotein - | RNVIPTITQMNLKYAISAKN |
| 4931 | ORF1b polyprotein - | ITQMNLKYAISAKNRARTVA |
| 4937 | ORF1b polyprotein - | KYAISAKNRARTVAGVSI |
| 4954 | ORF1b polyprotein - | ICSTMTNRQFHQKLLKS |
| 4964 | ORF1b polyprotein - | HQKLLKSIAATRGATVVIGT |
| 4970 | ORF1b polyprotein - | SIAATRGATVVIGTSKF |
| 4979 | ORF1b polyprotein - | VVIGTSKFYGGWHNMLKT |
| 5016 | ORF1b polyprotein - | RAMPNMLRIMASLVLARKHT |
| 5022 | ORF1b polyprotein - | LRIMASLVLARKHTTCCSLS |
| 5064 | ORF1b polyprotein - | SLYVKPGGTSSGDATT |
| 5104 | ORF1b polyprotein - | GNKIADKYVRNLQHRLY |
| 5162 | ORF1b polyprotein - | YASQGLVASIKNFKSVLYYQ |
| 5168 | ORF1b polyprotein - | VASIKNFKSVLYYQNNVF |
| 5244 | ORF1b polyprotein - | GTLMIERFVSLAIDA |
| 5250 | ORF1b polyprotein - | RFVSLAIDAYPLTKHPNQEY |
| 5256 | ORF1b polyprotein - | IDAYPLTKHPNQEYADV |
| 5278 | ORF1b polyprotein - | QYIRKLHDELTGHML |
| 5355 | ORF1b polyprotein - | YDHVISTSHKLVLSVNPY |
| 5363 | ORF1b polyprotein - | HKLVLSVNPYVCNAPGC |
| 5453 | ORF1b polyprotein - | RLKLFAAETLKATEETFK |
| 5486 | ORF1b polyprotein - | ELHLSWEVGKPRPPLNRN |
| 5531 | ORF1b polyprotein - | DAVVYRGTTTYKLNVGD |
| 5550 | ORF1b polyprotein - | VLTSHTVMPLSAPTLVPQEH |
| 5556 | ORF1b polyprotein - | VMPLSAPTLVPQEHYVR |
| 5572 | ORF1b polyprotein - | RITGLYPTLNISDEFSSNVA |
| 5586 | ORF1b polyprotein - | FSSNVANYQKVGMQKYSTLQ |
| 5592 | ORF1b polyprotein - | NYQKVGMQKYSTLQGP |
| 5596 | ORF1b polyprotein - | VGMQKYSTLQGPPGT |
| 5612 | ORF1b polyprotein - | KSHFAIGLALYYPSARIVYT |
| 5618 | ORF1b polyprotein - | GLALYYPSARIVYTAC |
| 5647 | ORF1b polyprotein - | KYLPIDKCSRIIPARA |
| 5701 | ORF1b polyprotein - | SMATNYDLSVVNARLRAKHY |
| 5707 | ORF1b polyprotein - | DLSVVNARLRAKHYVYIG |
| 5724 | ORF1b polyprotein - | GDPAQLPAPRTLLTK |
| 5775 | ORF1b polyprotein - | TVSALVYDNKLKAHKDKSAQ |
| 5781 | ORF1b polyprotein - | YDNKLKAHKDKSAQCF |
| 5792 | ORF1b polyprotein - | SAQCFKMFYKGVITHDVSSA |
| 5813 | ORF1b polyprotein - | NRPQIGVVREFLTRNPAWRK |
| 5819 | ORF1b polyprotein - | VVREFLTRNPAWRKAVFISP |

| | | |
|---|---|---|
| 5825 | ORF1b polyprotein - | TRNPAWRKAVFISPYNS |
| 5831 | ORF1b polyprotein - | RKAVFISPYNSQNAVASKIL |
| 5837 | ORF1b polyprotein - | SPYNSQNAVASKILG |
| 5845 | ORF1b polyprotein - | VASKILGLPTQTVDSSQGSE |
| 5851 | ORF1b polyprotein - | GLPTQTVDSSQGSEYD |
| 5881 | ORF1b polyprotein - | NVNRFNVAITRAKVGI |
| 5919 | ORF1b polyprotein - | RNVATLQAENVTGLF |
| 5967 | ORF1b polyprotein - | IPGIPKDMTYRRLISMMGFK |
| 5973 | ORF1b polyprotein - | DMTYRRLISMMGFKM |
| 5976 | ORF1b polyprotein - | YRRLISMMGFKMNYQVNGYP |
| 5982 | ORF1b polyprotein - | MMGFKMNYQVNGYPN |
| 5989 | ORF1b polyprotein - | YQVNGYPNMFITREEAIRH |
| 6025 | ORF1b polyprotein - | AVGTNLPLQLGFSTGV |
| 6053 | ORF1b polyprotein - | PNNTDFSRVSAKPPPGDQF |
| 6066 | ORF1b polyprotein - | PPGDQFKHLIPLMYKGLPWN |
| 6072 | ORF1b polyprotein - | KHLIPLMYKGLPWNVVRI |
| 6089 | ORF1b polyprotein - | IKIVQMLSDTLKNLSD |
| 6111 | ORF1b polyprotein - | WAHGFELTSMKYFVKIGPER |
| 6117 | ORF1b polyprotein - | LTSMKYFVKIGPERTCC |
| 6235 | ORF1b polyprotein - | RKVQHMVVKAALLADKFPVL |
| 6241 | ORF1b polyprotein - | VVKAALLADKFPVLH |
| 6340 | ORF1b polyprotein - | DGGSLYVNKHAFHTP |
| 6344 | ORF1b polyprotein - | LYVNKHAFHTPAFDKSAFV |
| 6454 | ORF1b polyprotein - | LENVAFNVVNKGHFDGQQG |
| 6470 | ORF1b polyprotein - | QQGEVPVSIINNTVY |
| 6474 | ORF1b polyprotein - | VPVSIINNTVYTKVD |
| 6485 | ORF1b polyprotein - | TKVDGVDVELFENKT |
| 6501 | ORF1b polyprotein - | LPVNVAFELWAKRNIKPVPE |
| 6507 | ORF1b polyprotein - | FELWAKRNIKPVPEVKILNN |
| 6513 | ORF1b polyprotein - | RNIKPVPEVKILNNLGVDIA |
| 6519 | ORF1b polyprotein - | PEVKILNNLGVDIAANTVI |
| 6531 | ORF1b polyprotein - | IAANTVIWDYKRDAPAHIST |
| 6537 | ORF1b polyprotein - | IWDYKRDAPAHISTI |
| 6580 | ORF1b polyprotein - | DGQVDLFRNARNGVLIT |
| 6595 | ORF1b polyprotein - | ITEGSVKGLQPSVGPKQAS |
| 6623 | ORF1b polyprotein - | AVKTQFNYYKKVDGVVQQLP |
| 6629 | ORF1b polyprotein - | NYYKKVDGVVQQLPE |
| 6649 | ORF1b polyprotein - | SRNLQEFKPRSQMEIDFLE |
| 6699 | ORF1b polyprotein - | GLHLLIGLAKRFKESPFELE |
| 6757 | ORF1b polyprotein - | IIKSQDLSVVSKVVKVT |
| 6802 | ORF1b polyprotein - | AWQPGVAMPNLYKMQRMLLE |
| 6808 | ORF1b polyprotein - | AMPNLYKMQRMLLEKCDLQ |
| 6831 | ORF1b polyprotein - | SATLPKGIMMNVAKYTQLC |

| | | |
|---|---|---|
| 6850 | ORF1b polyprotein - | QYLNTLTLAVPYNMRVIHFG |
| 6856 | ORF1b polyprotein - | TLAVPYNMRVIHFGAG |
| 6861 | ORF1b polyprotein - | YNMRVIHFGAGSDKGVAPGT |
| 6878 | ORF1b polyprotein - | PGTAVLRQWLPTGTLLVDS |
| 6924 | ORF1b polyprotein - | LIISDMYDPKTKNVTKEND |
| 6931 | ORF1b polyprotein - | DPKTKNVTKENDSKEGFFT |
| 6952 | ORF1b polyprotein - | CGFIQQKLALGGSVAIK |
| 7045 | ORF1b polyprotein - | MSKFPLKLRGTAVMSLKEGQ |
| 7051 | ORF1b polyprotein - | KLRGTAVMSLKEGQI |
| 7068 | ORF1b polyprotein - | MILSLLSKGRLIIRENNRVV |
| 7074 | ORF1b polyprotein - | SKGRLIIRENNRVVISSDVL |
| 7080 | ORF1b polyprotein - | IRENNRVVISSDVLVNN |
| 18 | ORF1a polyprotein - | LPVLQVRDVLVRGFGDS |
| 67 | ORF1a polyprotein - | PYVFIKRSDARTAPHGHVM |
| 105 | ORF1a polyprotein - | GVLVPHVGEIPVAYRKVLLR |
| 111 | ORF1a polyprotein - | VGEIPVAYRKVLLRKNGNKG |
| 117 | ORF1a polyprotein - | AYRKVLLRKNGNKGAGGHSY |
| 123 | ORF1a polyprotein - | LRKNGNKGAGGHSYGA |
| 249 | ORF1a polyprotein - | YELQTPFEIKLAKKFDTFNG |
| 255 | ORF1a polyprotein - | FEIKLAKKFDTFNGE |
| 275 | ORF1a polyprotein - | FPLNSIIKTIQPRVEKKKLD |
| 281 | ORF1a polyprotein - | IKTIQPRVEKKKLDGFMGRI |
| 287 | ORF1a polyprotein - | RVEKKKLDGFMGRIRSVYPV |
| 293 | ORF1a polyprotein - | LDGFMGRIRSVYPVASPNEC |
| 299 | ORF1a polyprotein - | RIRSVYPVASPNECNQ |
| 388 | ORF1a polyprotein - | HNESGLKTILRKGGRTIAFG |
| 394 | ORF1a polyprotein - | KTILRKGGRTIAFGGCV |
| 418 | ORF1a polyprotein - | NKCAYWVPRASANIG |
| 437 | ORF1a polyprotein - | GVVGEGSEGLNDNLLE |
| 441 | ORF1a polyprotein - | EGSEGLNDNLLEILQ |
| 449 | ORF1a polyprotein - | NLLEILQKEKVNINI |
| 476 | ORF1a polyprotein - | ILASFSASTSAFVETVK |
| 485 | ORF1a polyprotein - | SAFVETVKGLDYKAFKQIVE |
| 491 | ORF1a polyprotein - | VKGLDYKAFKQIVES |
| 507 | ORF1a polyprotein - | GNFKVTKGKAKKGAWNIGEQ |
| 530 | ORF1a polyprotein - | LSPLYAFASEAARVVRS |
| 536 | ORF1a polyprotein - | FASEAARVVRSIFSRTLETA |
| 542 | ORF1a polyprotein - | RVVRSIFSRTLETAQN |
| 550 | ORF1a polyprotein - | RTLETAQNSVRVLQKAAITI |
| 556 | ORF1a polyprotein - | QNSVRVLQKAAITILDGISQ |
| 691 | ORF1a polyprotein - | DSIIIGGAKLKALNLGETFV |
| 697 | ORF1a polyprotein - | GAKLKALNLGETFVTH |
| 754 | ORF1a polyprotein - | EEVVLKTGDLQPLEQPTSE |

| | | |
|---|---|---|
| 809 | ORF1a polyprotein - | TNNTFTLKGGAPTKVTF |
| 829 | ORF1a polyprotein - | TVIEVQGYKSVNITFELDE |
| 963 | ORF1a polyprotein - | GATSAALQPEEEQEED |
| 1036 | ORF1a polyprotein - | DNVYIKNADIVEEAKKV |
| 1050 | ORF1a polyprotein - | KKVKPTVVVNAANVYLKHGG |
| 1056 | ORF1a polyprotein - | VVVNAANVYLKHGGGVAGAL |
| 1062 | ORF1a polyprotein - | NVYLKHGGGVAGALNKA |
| 1069 | ORF1a polyprotein - | GGVAGALNKATNNAMQVESD |
| 1075 | ORF1a polyprotein - | LNKATNNAMQVESDDYI |
| 1188 | ORF1a polyprotein - | SSFLEMKSEKQVEQKIAEIP |
| 1209 | ORF1a polyprotein - | EEVKPFITESKPSVEQRKQ |
| 1218 | ORF1a polyprotein - | SKPSVEQRKQDDKKI |
| 1234 | ORF1a polyprotein - | ACVEEVTTTLEETKF |
| 1276 | ORF1a polyprotein - | ITFLKKDAPYIVGDVV |
| 1296 | ORF1a polyprotein - | LTAVVIPTKKAGGTTEMLA |
| 1310 | ORF1a polyprotein - | TEMLAKALRKVPTDN |
| 1324 | ORF1a polyprotein - | NYITTYPGQGLNGYTVE |
| 1392 | ORF1a polyprotein - | CVETKAIVSTIQRKYKGIKI |
| 1398 | ORF1a polyprotein - | IVSTIQRKYKGIKIQ |
| 1418 | ORF1a polyprotein - | YGARFYFYTSKTTVASLINT |
| 1424 | ORF1a polyprotein - | FYTSKTTVASLINTL |
| 1458 | ORF1a polyprotein - | NLEEAARYMRSLKVPATVSV |
| 1464 | ORF1a polyprotein - | RYMRSLKVPATVSVSSPD |
| 1476 | ORF1a polyprotein - | SVSSPDAVTAYNGYLTSS |
| 1485 | ORF1a polyprotein - | AYNGYLTSSSKTPEE |
| 1554 | ORF1a polyprotein - | DNLKTLLSLREVRTIKVFTT |
| 1560 | ORF1a polyprotein - | LSLREVRTIKVFTTVDNINL |
| 1566 | ORF1a polyprotein - | RTIKVFTTVDNINLHTQ |
| 1572 | ORF1a polyprotein - | TTVDNINLHTQVVDM |
| 1603 | ORF1a polyprotein - | DVTKIKPHNSHEGKTFYVLP |
| 1641 | ORF1a polyprotein - | SFLGRYMSALNHTKKWKYP |
| 1685 | ORF1a polyprotein - | QIELKFNPPALQDAYYR |
| 1690 | ORF1a polyprotein - | FNPPALQDAYYRARA |
| 1749 | ORF1a polyprotein - | NVVCKTCGQQQTTLK |
| 1759 | ORF1a polyprotein - | QTTLKGVEAVMYMGTLSY |
| 1798 | ORF1a polyprotein - | VQQESPFVMMSAPPAQYELK |
| 1804 | ORF1a polyprotein - | FVMMSAPPAQYELKHGT |
| 1873 | ORF1a polyprotein - | YTTTIKPVTYKLDGVVC |
| 1911 | ORF1a polyprotein - | PIDLVPNQPYPNASFDNFK |
| 1938 | ORF1a polyprotein - | ADDLNQLTGYKKPASRELKV |
| 1944 | ORF1a polyprotein - | LTGYKKPASRELKVTF |
| 1964 | ORF1a polyprotein - | NGDVVAIDYKHYTPSFKKGA |
| 1970 | ORF1a polyprotein - | IDYKHYTPSFKKGAKLLHKP |

| | | |
|---|---|---|
| 1976 | ORF1a polyprotein - | TPSFKKGAKLLHKPIVW |
| 1988 | ORF1a polyprotein - | KPIVWHVNNATNKAT |
| 2102 | ORF1a polyprotein - | NSSLTIKKPNELSRVLGL |
| 2112 | ORF1a polyprotein - | ELSRVLGLKTLATHGLAAVN |
| 2118 | ORF1a polyprotein - | GLKTLATHGLAAVNSVP |
| 2134 | ORF1a polyprotein - | PWDTIANYAKPFLNKVVSTT |
| 2140 | ORF1a polyprotein - | NYAKPFLNKVVSTTTNIVT |
| 2176 | ORF1a polyprotein - | LLQLCTFTRSTNSRIKASM |
| 2188 | ORF1a polyprotein - | SRIKASMPTTIAKNTVK |
| 2194 | ORF1a polyprotein - | MPTTIAKNTVKSVGKFCL |
| 2223 | ORF1a polyprotein - | FSKLINIIIWFLLLSVCLG |
| 2233 | ORF1a polyprotein - | FLLLSVCLGSLIYST |
| 2295 | ORF1a polyprotein - | LDSLDTYPSLETIQI |
| 2367 | ORF1a polyprotein - | LIINLVQMAPISAMVRMYI |
| 2404 | ORF1a polyprotein - | CNSSTCMMCYKRNRATRVEC |
| 2410 | ORF1a polyprotein - | MMCYKRNRATRVECT |
| 2424 | ORF1a polyprotein - | TTIVNGVRRSFYVYANG |
| 2468 | ORF1a polyprotein - | EVARDLSLQFKRPINPTDQS |
| 2474 | ORF1a polyprotein - | SLQFKRPINPTDQSSYI |
| 2499 | ORF1a polyprotein - | GSIHLYFDKAGQKTYER |
| 2537 | ORF1a polyprotein - | PINVIVFDGKSKCEESS |
| 2567 | ORF1a polyprotein - | QPILLLDQALVSDVG |
| 2575 | ORF1a polyprotein - | ALVSDVGDSAEVAVKM |
| 2592 | ORF1a polyprotein - | DAYVNTFSSTFNVPMEKLK |
| 2611 | ORF1a polyprotein - | TLVATAEAELAKNVSLD |
| 2627 | ORF1a polyprotein - | DNVLSTFISAARQGFVDSDV |
| 2655 | ORF1a polyprotein - | LKLSHQSDIEVTGDSCNNY |
| 2672 | ORF1a polyprotein - | NYMLTYNKVENMTPRDLGAC |
| 2691 | ORF1a polyprotein - | CIDCSARHINAQVAKSHNIA |
| 2697 | ORF1a polyprotein - | RHINAQVAKSHNIALIW |
| 2718 | ORF1a polyprotein - | FMSLSEQLRKQIRSAAKKNN |
| 2724 | ORF1a polyprotein - | QLRKQIRSAAKKNNLPFKL |
| 2751 | ORF1a polyprotein - | VNVVTTKIALKGGKIVNNWL |
| 2757 | ORF1a polyprotein - | KIALKGGKIVNNWLKQL |
| 2769 | ORF1a polyprotein - | WLKQLIKVTLVFLFV |
| 2791 | ORF1a polyprotein - | TPVHVMSKHTDFSSE |
| 2804 | ORF1a polyprotein - | SEIIGYKAIDGGVTRDIAST |
| 2853 | ORF1a polyprotein - | LIAAVITREVGFVVPGLPG |
| 2878 | ORF1a polyprotein - | NGDFLHFLPRVFSAV |
| 2936 | ORF1a polyprotein - | TNVLEGSVAYESLRPDTRY |
| 2956 | ORF1a polyprotein - | LMDGSIIQFPNTYLEGSV |
| 3033 | ORF1a polyprotein - | PLIQPIGALDISASI |
| 3061 | ORF1a polyprotein - | AYYFMRFRRAFGEYSHVVA |

| | | |
|---|---|---|
| 3100 | ORF1a polyprotein - | FLPGVYSVIYLYLTFY |
| 3122 | ORF1a polyprotein - | FLAHIQWMVMFTPLVPFW |
| 3156 | ORF1a polyprotein - | FFSNYLKRRVVFNGVSFS |
| 3193 | ORF1a polyprotein - | LRSDVLLPLTQYNRYLALYN |
| 3199 | ORF1a polyprotein - | LPLTQYNRYLALYNKYKYFS |
| 3205 | ORF1a polyprotein - | NRYLALYNKYKYFSGAMDTT |
| 3246 | ORF1a polyprotein - | SDVLYQPPQTSITSAVLQSG |
| 3252 | ORF1a polyprotein - | PPQTSITSAVLQSGFR |
| 3256 | ORF1a polyprotein - | SITSAVLQSGFRKMAFPSGK |
| 3262 | ORF1a polyprotein - | LQSGFRKMAFPSGKVEGCM |
| 3316 | ORF1a polyprotein - | NYEDLLIRKSNHNFLVQAGN |
| 3322 | ORF1a polyprotein - | IRKSNHNFLVQAGNVQL |
| 3328 | ORF1a polyprotein - | NFLVQAGNVQLRVIGHSMQN |
| 3334 | ORF1a polyprotein - | GNVQLRVIGHSMQNCVLK |
| 3361 | ORF1a polyprotein - | TPKYKFVRIQPGQTFSV |
| 3447 | ORF1a polyprotein - | PFVDRQTAQAAGTDTTIT |
| 3511 | ORF1a polyprotein - | DILGPLSAQTGIAVL |
| 3532 | ORF1a polyprotein - | KELLQNGMNGRTILGSALLE |
| 3559 | ORF1a polyprotein - | VVRQCSGVTFQSAVKRTIKG |
| 3565 | ORF1a polyprotein - | GVTFQSAVKRTIKGTHHW |
| 3579 | ORF1a polyprotein - | THHWLLLTILTSLLVLV |
| 3618 | ORF1a polyprotein - | IIAMSAFAMMFVKHKHAFL |
| 3633 | ORF1a polyprotein - | HAFLCLFLLPSLATVA |
| 3692 | ORF1a polyprotein - | LILMTARTVYDDGARRVW |
| 3716 | ORF1a polyprotein - | TLVYKVYYGNALDQAISM |
| 3798 | ORF1a polyprotein - | CLLNRYFRLTLGVYDYL |
| 3816 | ORF1a polyprotein - | STQEFRYMNSQGLLPPKNSI |
| 3822 | ORF1a polyprotein - | YMNSQGLLPPKNSID |
| 3834 | ORF1a polyprotein - | SIDAFKLNIKLLGVGGKPCI |
| 3840 | ORF1a polyprotein - | LNIKLLGVGGKPCIKVAT |
| 3849 | ORF1a polyprotein - | GKPCIKVATVQSKMSDVKCT |
| 3874 | ORF1a polyprotein - | SVLQQLRVESSSKLW |
| 3877 | ORF1a polyprotein - | QQLRVESSSKLWAQC |
| 3933 | ORF1a polyprotein - | EMLDNRATLQAIASEFSS |
| 3970 | ORF1a polyprotein - | NGDSEVVLKKLKKSLNVAKS |
| 3976 | ORF1a polyprotein - | VLKKLKKSLNVAKSEF |
| 4001 | ORF1a polyprotein - | LEKMADQAMTQMYKQARSED |
| 4007 | ORF1a polyprotein - | QAMTQMYKQARSEDKRAKV |
| 4018 | ORF1a polyprotein - | SEDKRAKVTSAMQTMLFT |
| 4036 | ORF1a polyprotein - | MLRKLDNDALNNIIN |
| 4058 | ORF1a polyprotein - | PLNIIPLTTAAKLMVVI |
| 4109 | ORF1a polyprotein - | VQLSEISMDNSPNLA |
| 4124 | ORF1a polyprotein - | WPLIVTALRANSAVKLQNNE |

| | | |
|---|---|---|
| 4130 | ORF1a polyprotein - | ALRANSAVKLQNNELS |
| 4146 | ORF1a polyprotein - | PVALRQMSCAAGTTQTA |
| 4165 | ORF1a polyprotein - | DDNALAYYNTTKGGRFVLA |
| 4184 | ORF1a polyprotein - | LLSDLQDLKWARFPKSD |
| 4230 | ORF1a polyprotein - | FIKGLNNLNRGMVLGSLA |
| 4247 | ORF1a polyprotein - | AATVRLQAGNATEVPAN |
| 4253 | ORF1a polyprotein - | QAGNATEVPANSTVLS |
| 6 | ORF10 | VFAFPFTIYSLLLCRMN |
| 21 | ORF10 | MNSRNYIAQVDVVNF |
| 35 | nucleocapsid phosphoprotein - | ARSKQRRPQGLPNNTASWFT |
| 41 | nucleocapsid phosphoprotein - | RPQGLPNNTASWFTA |
| 65 | nucleocapsid phosphoprotein - | KFPRGQGVPINTNSS |
| 76 | nucleocapsid phosphoprotein - | TNSSPDDQIGYYRRATRRIR |
| 82 | nucleocapsid phosphoprotein - | DQIGYYRRATRRIRGGDGKM |
| 88 | nucleocapsid phosphoprotein - | RRATRRIRGGDGKMKDLSPR |
| 159 | nucleocapsid phosphoprotein - | LQLPQGTTLPKGFYAEGS |
| 188 | nucleocapsid phosphoprotein - | SRSRNSSRNSTPGSSR |
| 192 | nucleocapsid phosphoprotein - | NSSRNSTPGSSRGTS |
| 198 | nucleocapsid phosphoprotein - | TPGSSRGTSPARMAG |
| 240 | nucleocapsid phosphoprotein - | QQQGQTVTKKSAAEASKKPR |
| 246 | nucleocapsid phosphoprotein - | VTKKSAAEASKKPRQ |
| 265 | nucleocapsid phosphoprotein - | TKAYNVTQAFGRRGPEQTQG |
| 271 | nucleocapsid phosphoprotein - | TQAFGRRGPEQTQGN |
| 301 | nucleocapsid phosphoprotein - | WPQIAQFAPSASAFFGM |
| 349 | nucleocapsid phosphoprotein - | QVILLNKHIDAYKTFPPTEP |
| 355 | nucleocapsid phosphoprotein - | KHIDAYKTFPPTEPKKD |
| 389 | nucleocapsid phosphoprotein - | QQTVTLLPAADLDDFSKQ |
| 395 | nucleocapsid phosphoprotein - | LPAADLDDFSKQLQQ |
| 33 | membrane glycoprotein - | CLLQFAYANRNRFLYI |
| 95 | membrane glycoprotein - | YFIASFRLFARTRSMWSFNP |
| 101 | membrane glycoprotein - | RLFARTRSMWSFNPE |
| 139 | membrane glycoprotein - | VIGAVILRGHLRIAGHHLGR |
| 145 | membrane glycoprotein - | LRGHLRIAGHHLGRCDIK |
| 166 | membrane glycoprotein - | KEITVATSRTLSYYKLGAS |
| 186 | membrane glycoprotein - | RVAGDSGFAAYSRYRIGNYK |
| 192 | membrane glycoprotein - | GFAAYSRYRIGNYKLN |
| 48 | envelope protein - | NVSLVKPSFYVYSRVKNLNS |
| 54 | envelope protein - | PSFYVYSRVKNLNSSRV |
